# Discovery of an Antiviral PROTAC Targeting the SARS-CoV-2 Main Protease Using an Allosteric Warhead

**DOI:** 10.1101/2025.05.03.652023

**Authors:** Christopher Veeck, Lennart Laube, Anke Werner, Wibke Diederich, Stephan Becker

**Affiliations:** Marburg University, Institute of Virology, 35043 Marburg, Germany; Marburg University, Center for Tumor Biology and Immunology, 35043 Marburg, Germany; German Center for Infection Research (DZIF), Partner Site Giessen-Marburg-Langen, 35043 Marburg, Germany

## Abstract

Targeted protein degradation represents a new paradigm in drug development. By hijacking the cellular ubiquitin-proteasome-system pathogenic proteins are degraded via heterobifunctional molecules, referred to as proteolysis targeting chimeras (PROTACs). However, to date, only few PROTACs targeting viral or proviral proteins have been developed. To explore the possibilities and advantages of antiviral PROTACs, we have developed an antiviral PROTAC against the SARS-CoV-2 main protease (M^Pro^) using an allosteric warhead. Here, we present the design of ten M^Pro^ degraders that were developed using pelitinib as a warhead, an allosteric binder of M^Pro^. Among several candidates, LLP019 emerged as the most potent molecule, capable of degrading up to 90% of M^Pro^ after ectopic expression in HEK293F cells. LLP019 displayed significant antiviral activity against several variants of concern of SARS-CoV-2 in infected Calu3 cells. In conclusion, we show that the development of antiviral PROTACs using an allosteric warhead represents a promising antiviral strategy, expanding the range of possible target proteins and ligands.

**SYNPOSIS:** We present a proof-of-concept study to construct a PROTAC targeting the SARS-CoV-2 main protease (M^Pro^) using a warhead that binds outside the catalytic pocket.

## INTRODUCTION

Targeted protein degradation (TPD) employs the cellular ubiquitin-proteasome system (UPS) to force degradation of a target protein by introduction of a heterobifunctional molecule with binding sites for both the target protein and an E3 ligase. Cellular E3 ubiquitin ligases (E3s) are an abundant class of endogenous proteins that poly-ubiquitinate target proteins, which leads to their degradation in the proteasome. To date, more than 600 cellular E3s are known, which underlines the redundancy and importance of this system ^1^. UPS is hijacked by proteolysis targeting chimeras (PROTACs), which are heterobifunctional molecules that consist of an E3-ligand and a ligand of the target protein to be degraded, the warhead joined by a linker ^2^. The forced interaction and formation of a ternary complex between PROTAC, target protein and E3 leads to recruitment of the ubiquitination complex, which ultimately results in the poly-ubiquitination and degradation of the target protein in proteasomes ^3–5^. This approach offers several advantages compared to classical high-affinity inhibitors: 1) It is possible to repurpose inhibitors or ligands of the target protein as warheads ^6^; 2) PROTACs display high specificity for their target ^7^; 3) low dose administration is possible due to their catalytic activity ^8^; 4) PROTACs have shown to be more resistant to mutations of the target protein ^6,9^ than other antivirals. In addition, PROTACs can be used to target otherwise undruggable proteins that lack binding pockets, have high structural flexibility, or are inaccessible due to their intracellular localization ^10^. In sum, TPD through PROTACs carries great potential for the development of antivirals and the therapy of infectious diseases ^11–14^. However, only few antiviral PROTACs have so far been reported ^6,15–17^.

Coronaviruses (CoV) have a wide-ranging ability to infect mammals, including humans. They belong to the order of *Nidovirales* and contain a single-stranded positive-sense RNA genome (+ssRNA). Prominent representatives include the severe acute respiratory syndrome (SARS) CoV and SARS-CoV-2 ^18,19^ as well as the Middle East respiratory syndrome (MERS) CoV ^20^. The polyprotein 1a/b (pp1/ab) of SARS-CoV-2 contains 16 non-structural proteins (NSPs), which are cleaved co- and posttranslationally by the viral papain-like proteinase (PL^Pro^, NSP3) and 3C-like proteinase or main protease (M^Pro^, NSP5) in the cytosol of the infected host cell ^21^. Following the translation of pp1/ab, the viral main protease (M^Pro^) is cleaved autocatalytically from the polyprotein before it cleaves pp1/ab at eleven distinct sites, facilitating the maturation of non-structural proteins essential for viral replication ^22^. Because of its pivotal role for the viral replication cycle M^Pro^ is an often-used target for the development of antivirals against coronaviruses ^23^.

The majority of reported antiviral PROTACs use warheads that address the catalytic cleft of the target protein ^6,17^. Here we analyzed whether ligands, which bind outside of the catalytic cleft of M^Pro^ can be utilized as warheads for antiviral PROTACs, thereby expanding the range of target proteins by including viral proteins without enzymatic activity on the one hand and the spectrum of possible warheads on the other hand. We modified pelitinib, an allosteric binder of the SARS-CoV-2 M^Pro^ ^24^, to design and develop several M^Pro^ degraders via rational linker design and analyzed their degradation ability in cello in detail.

## RESULTS & DISCUSSION

### Design strategy of anti-M^Pro^ PROTACs

For the design of the PROTAC, we selected pelitinib as the warhead. Pelitinib is an allosteric ligand of the SARS-CoV-2 M^Pro^ ^24,25^ (Figure 1A and Figure 1B). Structural data suggested that the interaction between pelitinib and M^Pro^ is primarily mediated by the cyanoquinoline moiety of pelitinib and Ser301 of M^Pro^ [24]. Notably, when pelitinib is bound to M^Pro^ its (dimethylamino)but-2-enamide side chain is solvent-exposed (Figure 1B and Figure 1C highlighted in brown). This orientation renders the side chain a suitable exit vector for the conjugation of the PROTAC’s linker moiety. Therefore, we synthesized DH03, according to a procedure published by Daigneault et al. ^26^ and Carmi et al. ^27^, and outlined in Scheme 1. In brief, commercially available N-(4-chloro-3-cyano-7-ethoxyquinolin-6-yl)acetamide was reacted with 3-chloro-4-fluoroaniline in a nucleophilic aromatic substitution reaction ^28^ to produce intermediate LLP001 in 84% yield. Subsequently, the acetyl group was cleaved with concentrated hydrochloric acid to provide the key intermediate DH03 in a yield of 74% ^27^ (Figure 1C). DH03 provides an amino group at position six which subsequently was used to add a variety of linkers connecting pelitinib to 4-hydroxy-thalidomide, a known ligand for the E3 ubiquitin ligase cereblon (CRBN) ^29,30^. Two synthesis strategies were used. In the first, the linker moiety was coupled with the CRBN-binding group and then with DH03. In the second, DH03 was first coupled with the corresponding linker moiety and then coupled with the CRBN-binding group (Figure 1D) ^31^.

**Figure 1:**
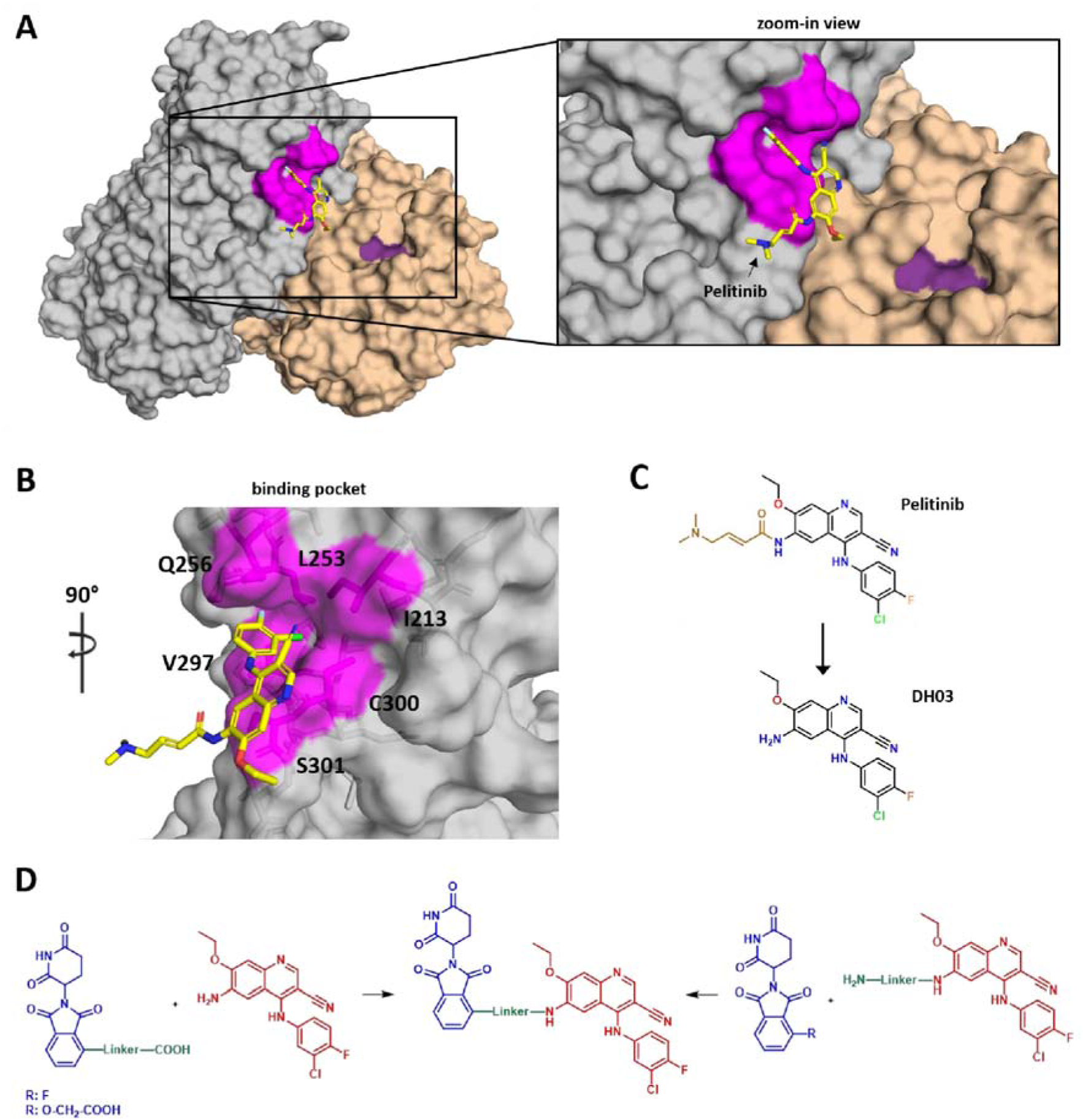
Representation of pelitinib as an allosteric warhead for anti-M^Pro^ PROTACs. (**A**) Pelitinib is a reported allosteric binder of the SARS-CoV-2 M (PDB: 7AXM), binding within a hydrophobic pocket formed by residues Ile213, Leu253, Gln256, Val297, and Cys300 (highlighted in magenta). The binding site is located in the N-terminal domain of M^Pro^ (protomer 1, shown in grey) and is spatially distant from the enzyme’s active site (red), which is part of protomer 2 (wheat). (**B**) Enlargement of pelitinib’s binding site. The (dimethylamino)but-2-enamide sidechain of pelitinib does not appear to interact with M^Pro^ and can therefore serve as a site for chemical modification and linker attachment. (**C**) Chemical structure of pelitinib and development of DH03 as a modified warhead. For the purpose of linker tethering, the (dimethylamino)but-2-enamide sidechain of pelitinib (highlighted in brown) was removed, resulting in the modified allosteric warhead DH03. (**D**) Construction of anti-M^Pro^ PROTACs. DH03 (red) was utilized as the warhead in the design of PROTACs, which were conjugated to thalidomide-derivatives (blue) via various linker modifications (green) to enable recruitment of the CRBN E3 ubiquitin ligase in two strategies. Left: The linker was first tethered to the thalidomide-based CRBN-binding group and subsequently connected to DH03. Right: The linker was first tethered to DH03 and subsequently attached to the thalidomide-derivative.

### Optimization of anti-M^Pro^ Degraders via Rational Linker Design

Using rational linker design, we systematically modified linker properties, such as linker length, hydrophobicity and flexibility to achieve maximal *in cellulo* degradation efficiency of the anti-M^pro^ PROTAC (Figure 2A) ^32–34^. We started with DH06 (Scheme S2), which contains a polyethylene glycol-2 unit (PEG-2) and thus the entire linker backbone consists of ten atoms (always C, N and O, unless otherwise stated). To determine the *in cellulo* degradation profile of M^Pro^ in the presence of DH06, HEK293F cells were transfected with plasmids encoding M^Pro^ or GFP and treated with increasing concentrations of DH06 or DMSO as a control. Expression of GFP served as a control. Cells were lysed at 24 h post transfection (hpt) and subjected to western blot analyses. Quantification of M^Pro^ and GFP signals revealed that treatment with DH06 resulted in the degradation of approximately 75% of M^Pro^. At a concentration of 9.4 µM, 50% of M^Pro^ was degraded (DC_50_ = 9.4 µM, Figure S1). Next, the backbone of the linker was extended to 13 atoms by the addition of an additional PEG unit (LLP037, Scheme S3). Compared to DH06, the extended linker in LLP037 slightly increased the degradation efficiency of M^Pro^ (Figure S2). To further optimize the linker, we replaced the terminal amino-group on the thalidomide-based moiety of LLP037 with a glycolic acid group, reducing the flexibility of this part of the molecule, while maintaining the linker length by removing one PEG group, resulting in LLP019 (Scheme 1 and Scheme S1). LLP019 had a significantly increased degradation rate of almost 85% of M^Pro^ (DC_50_ = 4.7 µM, Figure 2B). Subsequently, by the addition or removal of PEG units linker length was varied, ranging from three to 16 atoms in the backbone and their efficacy was tested (LLP038, Scheme S4; LLP031, Scheme S5; and LLP049, Scheme S6). None of the modifications improved degradation efficiency, suggesting that a linker length of 13 atoms is optimal when a rigid exit group is used. Replacement of the PEG-based linker of LLP019 with an alkyl linker, while maintaining the linker length, resulted in LLP041 (Scheme S7), which had significantly reduced activity. We assume that the increased hydrophobicity of LLP041’s alkyl linker led to a reduced solubility and thus lowered intracellular concentrations. Finally, we investigated linkers comprising short thalidomide-based exit groups and alkyl chains of five to eight atoms in the backbone (LP15, Scheme S8; LP08, Scheme S9; and LP04, Scheme S10). LP04, the PROTAC designed with the shortest linker exhibited a DC_50_-value of 4.4 µM, albeit with a lower D_max_. In summary, none of the latter adaptations led to PROTACs that surpassed the degradation efficiency of LLP019.

**Figure 2:**
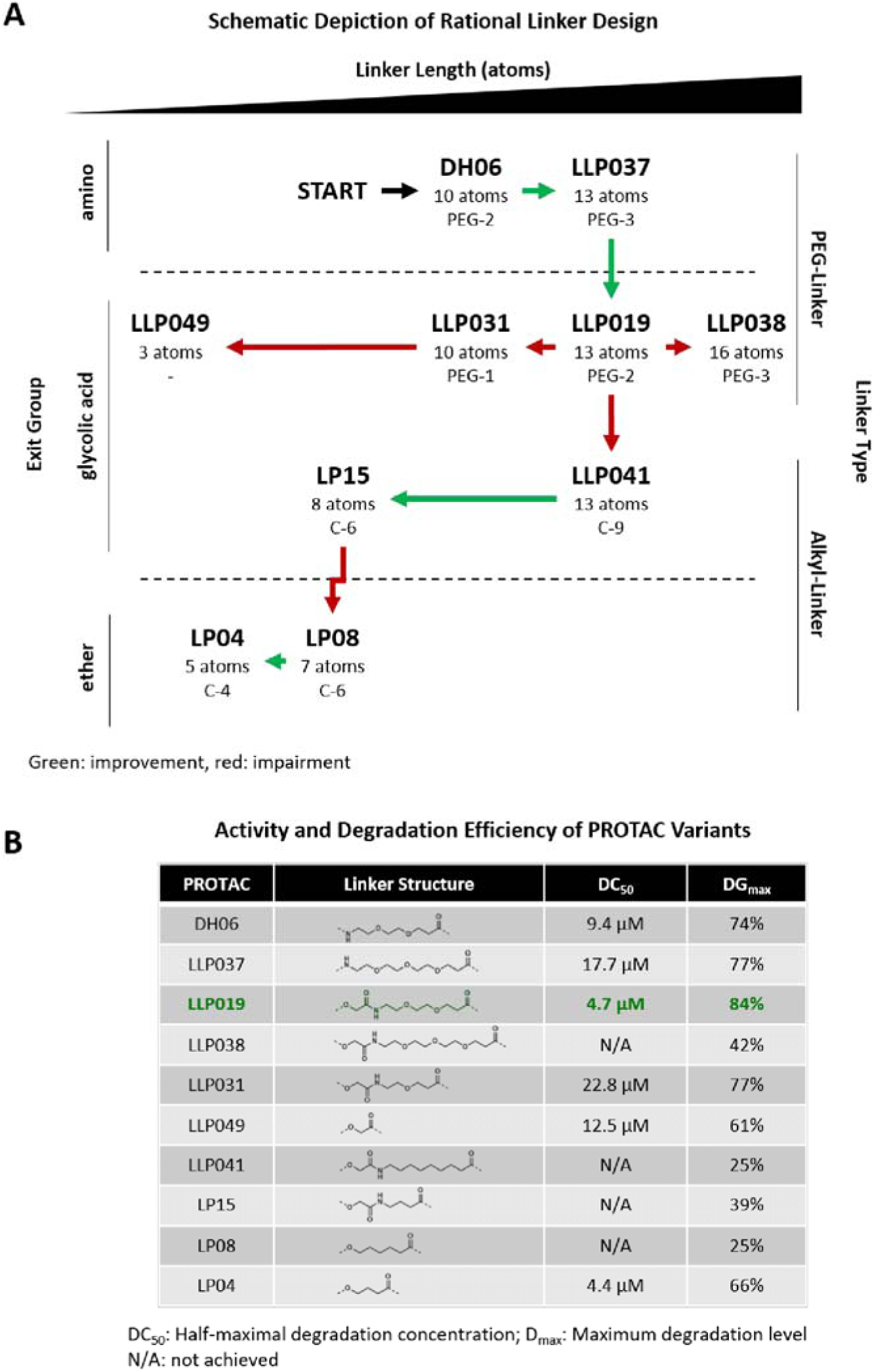
Representation of the rational linker design and summary of all assessed PROTACs. (**A**) Rational linker design strategy. Rational linker optimization was conducted by systematically varying three key properties: length, flexibility (exit group), and composition (PEG or alkyl linker). Starting with the initial PROTAC DH06, modifications in these parameters were explored to identify optimal M^Pro^ degradation efficiency. Beneficial adaptations are indicated by green arrows, while disadvantageous modifications are marked with red arrows. (**B**) Summary of PROTAC performance. A series of PROTAC derivatives (chemical structures provided in Figure S2) were developed and tested for their ability to degrade M^Pro^. Among all variants, LLP019 demonstrated the highest potency, with the lowest DC_50_ (4.7 µM) and the greatest maximal degradation (D_max_ = 84%). Detailed M^Pro^ degradation profiles for each PROTAC are provided in Figure S3.

### Synthesis of LLP019

Because LLP019 was the most potent PROTAC that has been developed and should therefore be further characterized, its synthesis is described in detail in the following paragraph (Scheme 1).

LLP019 was synthesized starting with the intermediate LLP016, which was prepared as described in the literature ^35,36^. Briefly, commercially available 3-hydroxyphthalic anhydride was first reacted with aminopiperidine-2,6-dione hydrochloride to form the intermediate LLP015 in a yield of 77%. This then reacted smoothly with tert-butyl bromoacetate in a substitution reaction to form LLP014 (80%). After deprotection of the carboxylic function with trifluoroacetic acid to give LLP016, this was further reacted with tert-butyl 3-[2-(2-aminoethoxy)ethoxy]propanoate in an amide coupling using HATU to give LLP017 in a yield of 67%. Cleaving off the tert-butyl ester with TFA renders LLP023, which is coupled to DH03, resulting in LLP019, albeit only with a yield of 10%. For all other PROTACs mentioned in this study, detailed experimental procedures and data can be found in the supporting information (Scheme S2-S10).

**Scheme 1:**
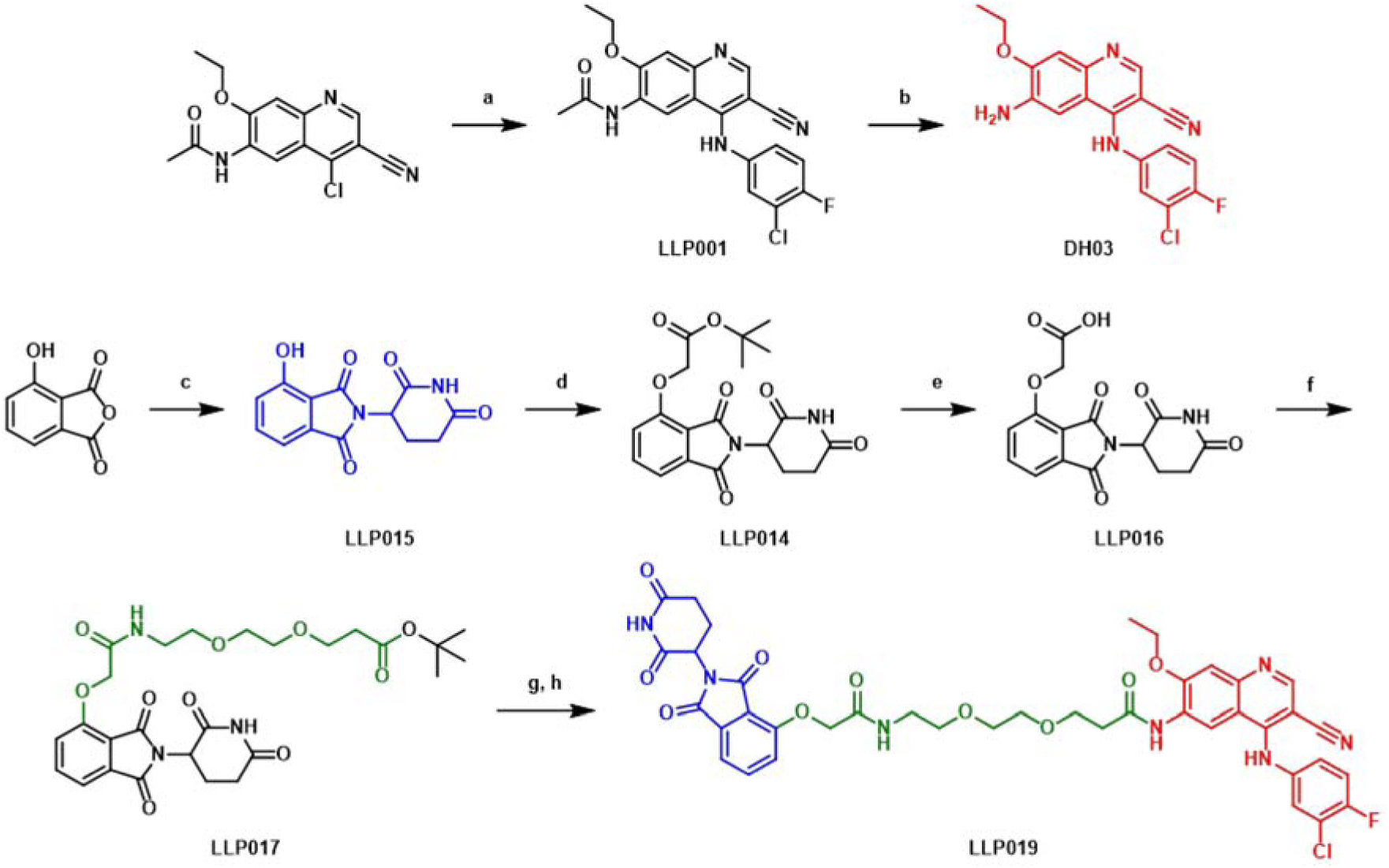
Synthesis of LLP019^a^. ^a^Reagents and conditions: (a) 3-chloro-4-fluoroaniline, methane sulfonic acid, 2-propanol, reflux, 6.5 h, 84% [30]; (b) HCl conc., H_2_O, reflux, overnight, 74% [29]; (c) 3-aminopiperidine-2,6-dione hydrochloride, KOAc, AcOH, reflux, 24 h, 77% [35]; (d) *tert*-butyl 2-bromoacetate, K_2_CO_3_, DMF, r.t., 2 h, 80%; (e) TFA, r.t., 4 h, 96%; (f) *tert*-butyl 3-[2-(2-aminoethoxy)ethoxy]propanoate, HATU, DIPEA, DMF, r.t., 19 h, 67%; (g) TFA, r.t., 4 h [34] (h) 6-amino-4-[(3-chloro-4-fluorophenyl)amino]-7-ethoxy-3-quinolinecarbonitrile (DH03), HATU, DIPEA, DMF, r.t., 19 h, 10%.

### LLP019 is a Potent Degrader of Ectopically Expressed M^Pro^

LLP019 (scheme 1) showed a linear dose-dependent degradation between 1.56 µM and 25 µM, plateauing at 50 µM followed by a slight reversal of activity at 100 µM (Figures 3A and Figure 3B). This characteristic degradation profile is commonly observed with PROTACs and is referred to as the hook effect ^37–42^.

**Figure 3:**
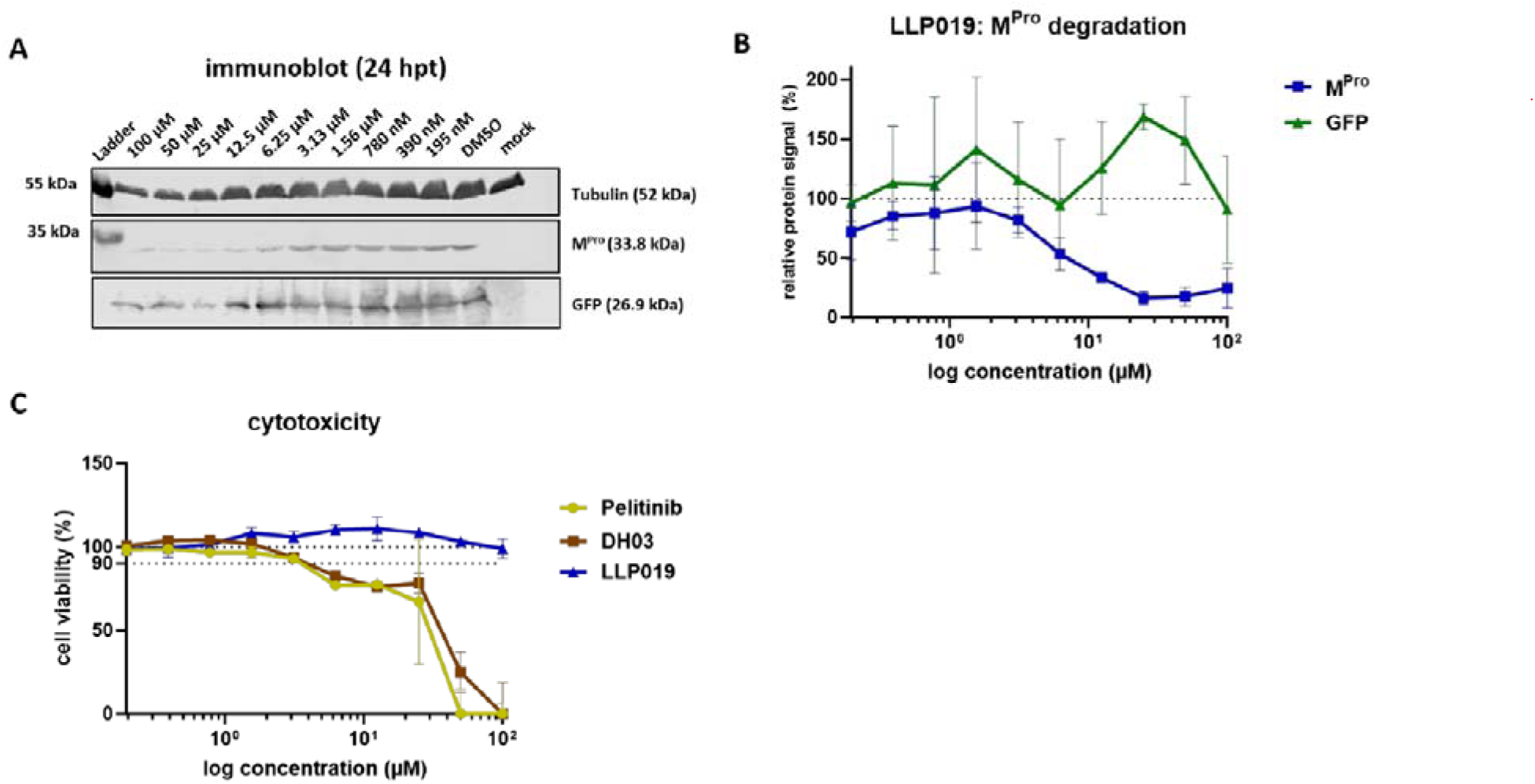
LLP019 is a potent M^Pro^ degrader. (**A**) Western blot analysis of M^Pro^ degradation. HEK293F cells were seeded in 6-well plates, and the medium was replaced with DMEM++ containing 3% FCS and increasing concentrations of LLP019 or DMSO (0.1%) as a control. Cells were subsequently transfected with 500 ng each of pCAGGS-M^Pro^ and pCAGGS-GFP. At 24 hpt, cells were harvested, and lysates were analyzed by SDS-PAGE and western blotting. The panel shows a representative western blot image from three independent experiments. (**B**) Quantification of M^Pro^ degradation. Western blot signals for M^Pro^ and GFP were quantified, normalized to tubulin, and compared to the DMSO (0.1%) control. (C) Cell viability analysis. HEK293F cells were seeded in 96-well plates and treated with increasing concentrations of LLP019, DH03, pelitinib, or DMSO (0.1% v/v final concentration). After 48 hours, ATP-dependent cell viability was measured using the CellTiter-Glo^®^ 2.0 Assay (Promega). Viability data were normalized to the DMSO (0.1%) control.

The hook effect likely arises due to increasing saturation of the PROTAC binding sites on both the E3 ligase cereblon (CRBN) and the target protein M^Pro^. At an excess of PROTAC molecules the formation of ternary complexes between PROTAC, CRBN and M^pro^ is inhibited in favor of binary complexes of the PROTAC with either CRBN or M^Pro^ alone. Therefore, the productive complex required for ubiquitination and proteasomal degradation is no longer formed^43^. This phenomenon has been extensively characterized in PROTAC pharmacology and represents a key consideration for optimizing degrader efficacy.

To assess cytotoxicity, LLP019 was evaluated in HEK293F cells over 48 hours using ATP-based and NADPH/NADH-dependent cell metabolism assays (Figure 3C and Figure S5B). Slight cytotoxicity of LLP019 was observed at 100 µM. In subsequent experiments, the maximum concentration of LLP019 was therefore kept at 50 µM. Interestingly, when tested separately, pelitinib and DH03 caused significant cytotoxicity in HEK293F cells (Figure 3C).

To validate that LLP019 acts by employing the UPS, two compounds with distinct mechanisms of UPS inhibition were tested. The first compound, pevonedistat (MLN4924), blocks the NEDD8-activating enzyme, thereby preventing the activation of cullin ring ligases, which play a key role in degrading numerous regulatory proteins ^44–48^. The second compound, TAK243 (MLN7243), inhibits the formation of the ubiquitin-activating enzyme 1-ubiquitin conjugate, a critical step required for protein degradation via the proteasome ^49^. M^Pro^- and GFP-expressing HEK293F cells were treated at 24 h post transfection with LLP019 (25 µM) and pevonedistat (10 µM) or TAK243 (10 µM) ^44–49^. Both inhibitors exhibit cytotoxic effects after 24 h exposure (not shown) therefeore, cells were harvested already at 5 h after inhibitor treatment to minimize toxic effects (Figure 4A). In addition, to assess degradation efficiency without interference from continuous plasmid-driven M^Pro^ synthesis, cycloheximide (10 µM) was added at the start of the treatment to inhibit translation. We observed that under these conditions treatment with LLP019 resulted in degradation of almost 40% of M^Pro^ after 5 hpt compared to DMSO controls. As anticipated, treatment with pevonedistat or TAK243 inhibited LLP019-induced M^Pro^ degradation (Figure 4A). Pevonedistat completely abolished the effect of LLP019, whereas TAK243 showed a weaker inhibitory effect. Under these conditions, the GFP signal remained virtually unaffected (Figure 4A). This result supports the idea that the cellular UPS is involved in the LLP019-mediated degradation of M^Pro^.

**Figure 4:**
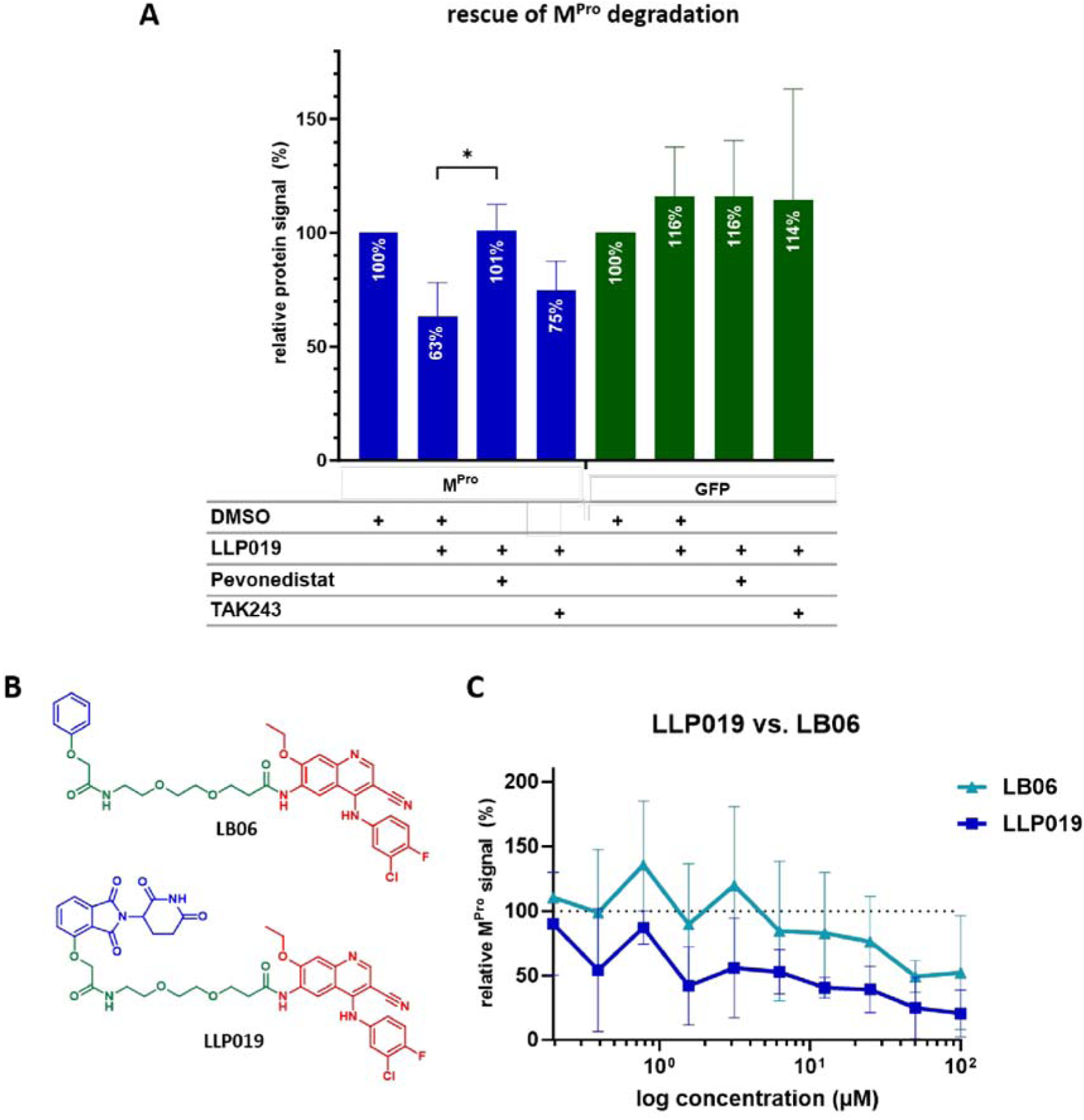
LLP019’s mode of action. (**A**) HEK293F cells were transfected with 500 ng of each pCAGGS-M^Pro^ and pCAGGS-GFP. At 24 hpt, transfection medium was changed and cells treated with cycloheximide (10 µM). In addition, DMSO (0.1% v/v), LLP019 (25 µM) or LLP019 (25 µM) were added as well as pevonedistat (10 µM) or TAK243 (10 µM). After 5 h, cells were harvested and subjected to SDS-PAGE and western blot (Figure S4A). Finally, M^Pro^ and GFP signals were quantified, normalized to tubulin and compared to DMSO control (= 100%). Statistics were performed via one-way ANOVA. Asterisks indicate statistical significance as follows: *p < 0.05. The presented data contains three independent experiments. (B) LB06 is an LLP019 derivative that lacks thalidomide (blue) but contains the warhead DH03 (red) and a linker (green) to allow binding of LB06 to M without being able to recruit CRBN. (C) HEK293F cells were seeded in 6-well plates and transfection with 500 ng of each pCAGGS-M^Pro^ and pCAGGS-GFP. After 4 h, medium was changed to DMEM++ with 3% FCS and increasing concentrations of LLP019, LB06 or DMSO (0.1%). At 24 hpt, cells were harvested and lysates subjected to SDS-PAGE and western blot (Figure S4B). Western blot signals of M and GFP were quantified, normalized to tubulin and compared to DMSO. The presented data contains three independent experiments.

To explore whether the warhead pelitinib alone was capable of inducing the degradation of M^Pro^, we used an LLP019 derivative, LB06 (Scheme S12), in which the thalidomide-based moiety was reduced to a phenyl ring that should be unable to recruit cereblon (Figure 4B). The cytotoxicity of this compound is shown in Figure S5A. We observed, a substantially lower degradation of M^Pro^ in presence of LB06 compared to LLP019 (Figure 4C), most likely because LB06 lacks moieties of thalidomide important for binding to CRBN, while interaction with M^Pro^ is presumably maintained. The residual M^Pro^ degradation activity induced by LB06 might be caused by warhead-induced destabilization of M^Pro 50,51^. Hence, we conclude that LLP019 acts as a heterobifunctional molecule that induces degradation of M^Pro^ via the recruitment of CRBN. Ligands such as LB06 are particularly valuable in the development of antiviral PROTACs as they allow differentiation of the intrinsic antiviral activity of the warhead and the effects mediated by targeted protein degradation. In summary, our data strongly supports that LLP019 induces CUL4A^CRBN^-dependent degradation of M^Pro^ in proteasomes.

LLP019 degraded ca. 40% of ectopically expressed M^Pro^ after 5 hpt (in the presence of cycloheximide) and ca. 85% after 24 hpt. Comparing the DC_50_ of LLP019 to another reported degraders of M^Pro^, it appears to be in a similar range ^17^. Nevertheless, some published degraders, predominantly of endogenous cellular kinases, degraded almost 100% of their target protein within 2-16 h at lower concentrations than determined for LLP019 ^52,53^. The difference might be caused by the significantly lower expression levels and substantially slower expression kinetic of the endogenous target proteins in comparison to the ectopically expressed viral proteins ^36,54–59^.

### LLP019 is a Heterobifunctional Molecule with Antiviral Potency

Next, we proceeded to evaluate LLP019’s antiviral efficacy in the context of SARS-CoV-2 infection. To this end, we pre-incubated Calu3 cells, a lung epithelial cell line which is often used for SARS-CoV-2 infections, with LLP019 for 16 h prior to infection with SARS-CoV-2 (BavPat1/2020 ^60^, henceforth abbreviated with BavPat1). After one hour of infection, virus inoculum was removed and cells were further treated with LLP019, DMSO or nirmatrelvir, which served as an inhibitor control. Samples were taken after 24 and 48 hours post infection (hpi) and viral titers were determined via TCID_50_ to determine half maximal effective concentration (EC_50_) values. After 48 hpi, viral titers in the supernatants of cells treated with 25 µM LLP019 were significantly reduced by ca. 85% (Figure 5A). Interestingly, this effect is reduced at flanking concentrations of 50 µM and 12.5 µM, which we assumed to be part of the hook-effect that was observed before (Figure 3B). To further investigate this phenomenon, the titration of LLP019 was repeated using smaller dilution steps between 50 µM and 10 µM. As expected, we observed a bell-shaped dose-dependence (EC_50_ = 13.03 µM), which resembles a hook, further underlining the presumed mode of action of LLP019 (Figure 5B).

**Figure 5:**
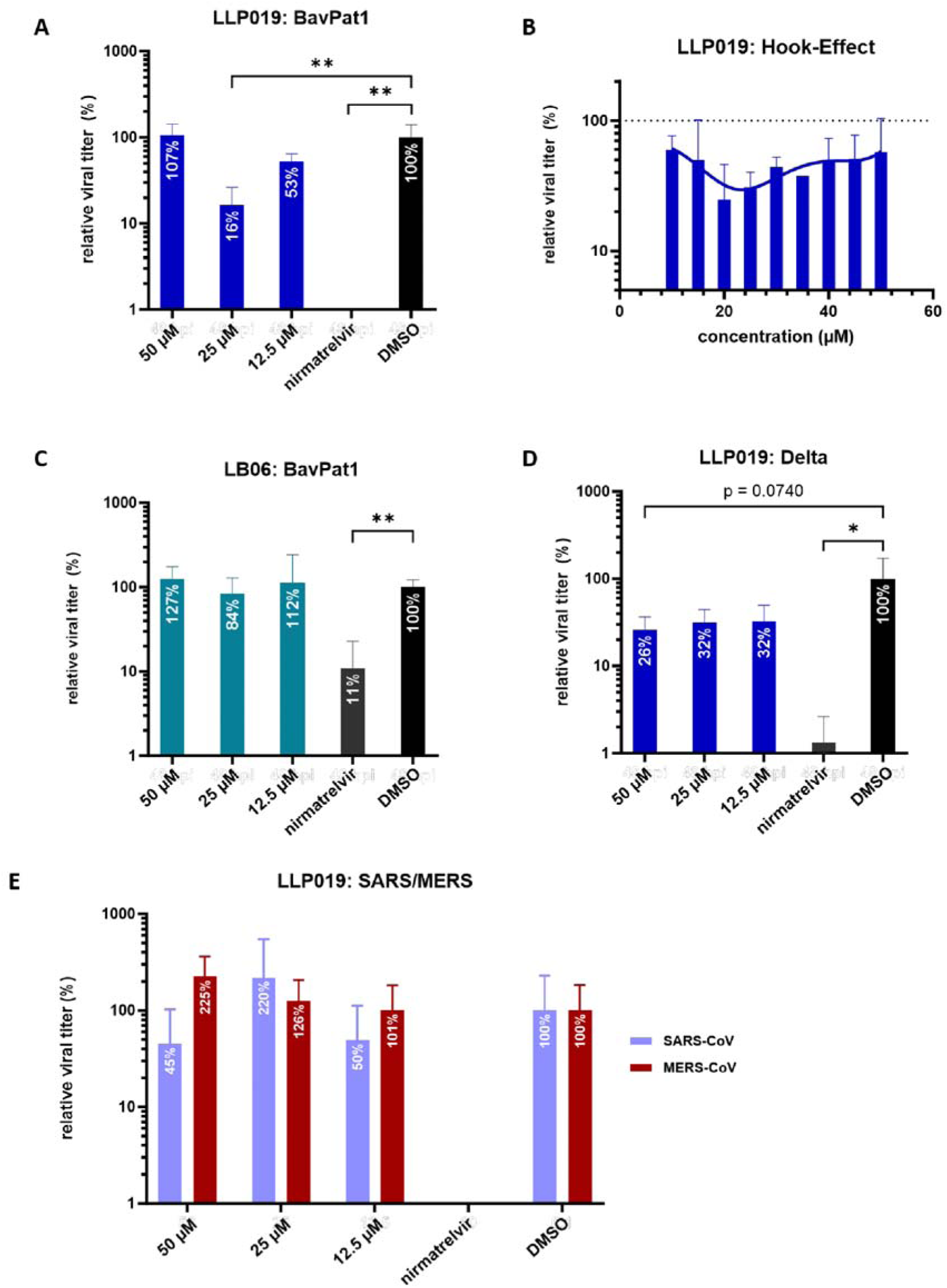
LLP019 displays specific SARS-CoV-2-directed antiviral activity. (**A, B**) At 80-90% confluency, Calu3 cells were pre-incubated with medium containing LLP019 in increasing concentrations, nirmatrelvir (10 µM) or DMSO (0.1% v/v). After 16 h, cells were infected with SARS-CoV-2 BavPat1 (MOI=0.01). After one hour, medium was changed to DMEM++/F12 containing 3% FCS as well as increasing concentrations of LLP019, nirmatrelvir (10 µM) or DMSO (0.01%). Supernatant was collected 48 hpi. Viral titers were assessed via TCID_50_, normalized to DMSO and DMSO-treated cells were set to 100%. (**C**) Calu3 cells were infected with SARS-CoV-2 BavPat1 (MOI=0.1) as described above. Supernatants were harvested after 48 hpi and viral titers were assessed via TCID_50_. Similarly, Calu3 cells were infected with (**D**) SARS-CoV-2 Delta (MOI=0.01) or (**E**) SARS-CoV/MERS-CoV (MOI=0.01) and treated as described. Supernatants were harvested after 48 hpi and viral titers were assessed via TCID_50_, normalized to DMSO and DMSO-treated cells were set to 100%. Statistical analyses were performed via unpaired t-test (**C**) or one-way ANOVA (A, D). Asterisks indicate statistical significance as follows: *p < 0.05, **p < 0.005. All presented experiments (A-E) represent three independent replicates.

To analyze whether the observed antiviral activity of the heterobifunctional LLP019 is in part caused by the warhead only, we treated SARS-COV-2-infected cells with LB06 (Figure 4B). As LB06 did not reduced viral titers, the observed antiviral activity of LLP019 is most likely mediated by TPD and not due to allosteric inhibition of M^Pro^’s enzymatic activity by the warhead (Figure 5C).

Next, we treated cells infected with the Delta variant of SARS-CoV-2 (German isolate FFM-IND8424/2021) ^61^ with LLP019. We observed a 75% titer reduction at 50 µM of LLP019 (Figure 5D). To further assess whether LLP019 displayed cross-species antiviral activity, we infected Calu3 cells with SARS-Co-V and MERS-CoV and treated them with LLP019 under identical conditions as described above. Here we could show that LLP019 had a tendency to reduce titers of SARS-CoV (to 55% at concentrations of 50 µM) although the effect was not statistically significant. MERS-CoV was not inhibited by LLP019 treatment (Figure 5E).

It is described that pelitinib interacts with Ser301 of M^Pro^ via its cyanoquinoline moiety ^24^. Although Ser301 is conserved in SARS-CoV, structural differences at the C-terminus result in masking of the pelitinib binding site via Glu256, which could explain the observed decrease of LLP019 antiviral activity in comparison to SARS-CoV-2 (Figure S6 and Figure S7). In M^Pro^ of MERS-CoV, position 301 is a methionine which results in a significant structural change, explaining the failure of LLP019 to inhibit MERS-CoV (Figure S7).

Our data show that LLP019 is capable to inhibit SARS-CoV-2 BavPat1 (Figure 5A and Figure 5B) and Delta (Figure 5D) in cell culture at non-toxic concentrations via forced M^Pro^ degradation. Using the control derivative LB06, which lacks large moieties of the CRBN-binding group of LLP019, we were able to show that LLP019 acts as an antiviral degrader independent from its warhead DH03 (Figure 5C), further emphasizing that targeted degradation of M^Pro^ is a promising antiviral strategy.

## CONCLUSION

In conclusion, we report the design and synthesis of an antiviral PROTAC, LLP019, targeting the M^pro^ of SARS-CoV-2. LLP019 is a heterobifunctional molecule based on thalidomide, a ligand of CRBN and an M^pro^ ligand that binds outside of the catalytic cleft and did not show inhibitory properties itself. This study demonstrates that warheads that do not inhibit the enzymatic activity of the target protein are also suitable for PROTAC design, thereby expanding the range of warheads available. Furthermore, repurposing ligands of viral proteins lacking druggable enzymatic activity seems to be feasible. Furthermore, we showed that rational linker design focusing on length, solubility and flexibility proved to be a promising strategy to optimize degradation efficiency. Although it must be noted that further studies are needed to evaluate the factors that contribute to successful degradation of a viral target protein and the development of potent antiviral degraders, this work serves as a proof-of-concept for such an approach and highlights TPD as a promising tool for novel antiviral strategies.

## MATERIALS & METHODS

### Cell culture and reagents

Human embryonic kidney 293 freestyle (HEK293F) cells were cultured in Dulbecco’s modified Eagle’s medium (DMEM, Thermo Fisher Scientific, Waltham, MA, USA) supplemented with penicillin (100 U/mL), streptomycin (100 μg/mL) (P/S), 2 mM glutamine (Q) (abbreviated as DMEM++) and 3 or 10% fetal calf serum (FCS). Cultured human airway epithelial 3 (Calu-3) cells were cultured in DMEM/Nutrient Mixture F-12 (DMEM/F12) supplemented with penicillin (50 U/mL), P/S (50 μg/mL), 2 mM Q (abbreviated as DMEM/F12++) and 3 or 10% FCS.

### Plasmids

Codon-optimized (*homo sapiens*) cDNA encoding the Main protease (M^Pro^) of SARS-CoV-2 Wuhan-Hu-1 (GenBank accession number MN908947) with HA-tag its N-terminal domain (YPYDVPDYA) was synthesized at Life Technologies and subcloned into the pCAGGS expression plasmid using EcoRI and XhoI restriction enzymes (pCAGGS-HA-M^Pro^).

### Antibodies

Ectopically expressed M^Pro^ was stained with a monoclonal mouse anti-HA antibody (Sino Biological, #100028-MM10, dilution 1:500). Endogenous tubulin was stained with a monoclonal antibody from Sigma-Aldrich (#T9026, dilution 1:1000). GFP was stained with a polyclonal goat anti-GFP antibody (Rockland, #600-101-215, dilution 1:1000).

IRDye-labeled secondary antibodies were purchased from LI-COR (anti-mouse-IRDye800CW, #926-32210; anti-rabbit-IRDye680, #926-68071) and diluted 1:5000. Alexa Fluor-labeled secondary antibodies were purchased from Thermo Fisher Scientific (anti-goat-Alexa Fluor 680, #A-21084) and diluted 1:5000 for Western blot analyses.

### Transfection of plasmids in HEK293F cells

For ectopic expression of proteins, 60–80% confluent HEK293F cells in 6-well plates were transfected with 500 ng pCAGGS-M^Pro^ and pCAGGS-GFP each, or 1000 ng pCAGGS empty vector (mock) using TransIT transfection reagent (Mirus Bio, Madison, WI, USA, MIR 2300) according to manufacturer’s protocol.

### SDS–PAGE, western Blot analysis and quantification

Cells were harvested at 24 h or 29 h post transfection (hpt), respectively, and centrifuged for 5 min at 5000×g at room temperature. Cells were washed with PBS_def_, lysed in 50 µL passive lysis buffer (Promega, #E1941) for at least 30 min at room temperature and centrifuged again for 10 min at 14,000 rpm. Then, 40 µL of supernatant was supplemented with 14 µL reducing SDS–PAGE sample buffer and heated at 95 °C for 10 min. Proteins were then subjected to SDS–PAGE (12% or 15%, respectively, poly acrylamide gel) and transferred to nitrocellulose membrane (0.45 µm, Cytiva, #10600002) at 25 V for 30 minutes. After 16 h of incubation in blocking buffer (10% skim milk in PBS_def_) at 4°C, detection of target proteins was performed using primary antibodies and species-specific IRDye/Alexa Fluor-coupled secondary antibodies (specified under “Antibodies”). Signals were visualized using the Odyssey CLx Imaging System by LI-COR and quantified by Image studio software version 5.2 (LI-COR).

### Cytotoxicity Assay

HEK293F cells (1×10^4^/well) were seeded in 96-well plates in DMEM++/10%FCS overnight (60-80% confluency). On the next day, medium was changed to 100 µL DMEM++/3%FCS containing decreasing concentrations of PROTACs or DMSO/controls. After 48 h, ATP-dependent cell metabolism was determined using the CellTiter-Glo^®^ 2.0 Cell Viability Assay (Promega, #G9241). To this end, 25 µL of substrate was added to the medium and incubated in the dark at room temperature (RT) for 10 min. Next, 50 µL of the supernatant was transferred to opaque white 96-well plates and luminescence was measured via a CentroLB 960 luminometer (Berthold Technologies). For NADH/NADPH-dependent cell metabolism 96^®^ Aqueous One Solution Cell Proliferation Assay (Promega, #G3582) following the manufacturer’s instructions.

### General Synthesis

Unless otherwise stated, commercially available solvents and reagents were used without further purification. For reactions involving reagents that are sensitive to moisture or oxygen, dried solvents were used and the reactions were carried out in preheated glassware in an inert gas atmosphere (Argon N50, Air Liquide). Reactions were cooled either with ice, ice/salt mixtures, dry ice/acetone or dry ice. Reaction control was performed by means of thin-layer chromatography (TLC) using aluminum-coated TLC plates 60 F_254_ from Merck or via an HPLC-MS system [Agilent 1260 Infinity system equipped with a Poroshell column (120, EC-C18, 2.7 μM, 4.6 mm × 50 mm, Agilent) and an Advion Expression S CMS]. Purification of the crude reaction products was carried out by means of flash chromatography (MPLC) on a Reveleris X2 (Grace, now Büchi) or an Interchim PuriFlash 5.020 system, using pre-packed columns of different sizes from two manufacturers (FlashPure from Büchi, PuriFlash from Interchim). For preparative HPLC, a Büchi Pure C-850 FlashPrep system with a PrepPure column (C18, 100 Å, 10 µm, 250 mm × 20 mm, Büchi) was used, with standard conditions (5-95% MeCN in H_2_O over 1 h) being applied unless otherwise stated. For drying isolated products under reduced pressure in a high vacuum system, optionally at elevated temperature, either a Büchi B-585 glass oven connected to an RC 6 chemistry hybrid pump with a VAP 5 manometer (both vacuum strips) or a Biotage V-10 Touch system was used. The drying time varied from a few hours to several days. The NMR spectra were recorded on either an AV III 300 MHz (Bruker), an ECZ400S or ECA500 (JEOL) spectrometer and the Delta 5.2.1 software (JEOL) was used for data processing and evaluation. The reported chemical shifts (in ppm) were referenced to the residual solvent signal used according to literature as follows: CDCl_3_: 7.26/77.16; DMSO-d_6_: 2.50/39.52 (^1^H/^13^C) ^62^. The following abbreviations were used to indicate spin multiplicities: s =singlet, d =doublet, t =triplet, q =quartet, m =multiplet, (b) =broad signal, sm =symmetric multiplet, dd =doublet of doublets, dt =doublet of triplets. HR-MS spectra were recorded on an AccuTOF-GCv (JEOL) or LTQ-FT (Thermo Fisher Scientific) system. Elemental analysis was determined using a CHN(S) analyzer vario MICRO CUBE (Elementar). The recording of NMR spectra and HR-MS spectra, as well as the determination of the elemental analysis, were carried out in the analytical facilities for NMR, MS and elemental analysis in the Chemistry and Pharmacy Departments.

### SARS-CoV-2 infection of Calu3 cells

Briefly, Calu3 cells were seeded in 6-well plates and grown for one week at 37 °C prior to infection, achieving 60-70% confluency at 24 h prior to infection. At 16 h prior to infection medium was changed to DMEM/F12 (3% FCS) containing PROTAC, nirmatrelvir (10 µM) or DMSO (0.1%). After 16 h, medium was changed to 1 mL DMEM (0% FCS) containing SARS-CoV (German isolate FFM-1 ^63^, GenBank accession number AY310120), MERS-CoV (EMC/2012; NC_019843.3 ^64^), SARS-CoV-2 BavPat1 (NR-52370, European Virus Archive global Cat#026V-03883, ^65^) or Delta B.1.617.2 (FFM-IND8424/2021, GenBank ID: MZ315141 ^61^). After one hour, cells were washed with PBS_def_ and incubated with DMEM/F12 (3% FCS) containing PROTAC, nirmatrelvir or DMSO for 48 h. Viral titers were assessed using tissue culture infectivity dose 50 (TCID_50_) assays after 24 and/or 48 hours post infection (hpi), respectively, as described by Spearman & Kärber ^66^.

## SUPPORTING INFORMATION

M^Pro^ degradation profile of DH06, chemical structures of all developed PROTACs, M^Pro^ degradation profile of all developed PROTACs, western blots of the LLP019 mode of action confirmation, cytotoxic profiles of additional compounds, structural alignment of SARS-CoV-2 and SARS-CoV M^Pro^, Structural alignment of pelitinib’s binding site in SARS-CoV-2, SARS-CoV and MERS-CoV M^Pro^, synthesis schemes of all developed PROTACs, ^1^H and ^13^C NMR spectra of all synthesized PROTACs

## ACKNOWLEDGEMENTS

We thank Marek Widera and the Frankfurt Virus Platform of the Goethe University & University Hospital Frankfurt for providing us virus isolates and associated protocols (for further information contact Denisa Bojkova, Sandra Ciesek). The authors express their deepest gratitude to Georg Winter (Center for Molecular Medicine Vienna) for his invaluable consultation, continuous support throughout the project, and provision of samples. We emphatically express our gratitude for Astrid Herwig for excellent technical support during this work. We also express our gratitude for Gotthard Ludwig, Sebastian Schmidt and Dr. Markus Eickmann for technical support during experiments in the BSL-4 laboratory. We acknowledge Laura Bunt, Justus Happe, Dominique Hagene, and Lukas Plamper for their support in the synthesis. We thank the European Virus Archive Global (EVA-G) for sample provision. Furthermore, we thank the Jürgen Manchot Stiftung, the LOEWE Center DRUID and the Uniscientia Stiftung for funding.

---

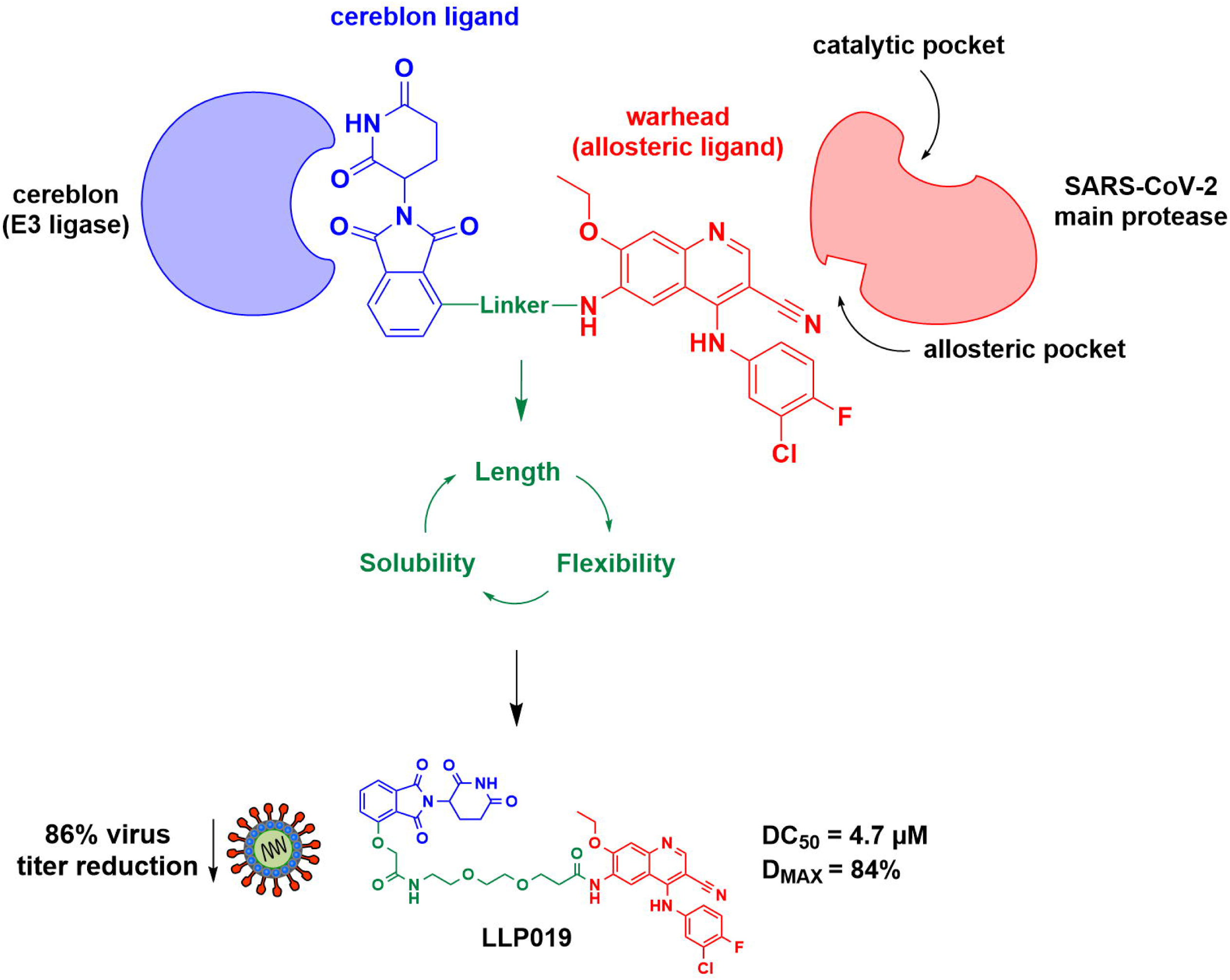

## Notes

### Competing Interest Statement

The authors have declared no competing interest.

