## Supporting Information for "Discovery of an Antiviral PROTAC Targeting the SARS-CoV-2 Main Protease Using an Allosteric Warhead"

*N-{4-[(3-Chloro-4-fluorophenyl)amino]-3-cyano-7-ethoxyquinolin-6-yl}-3-{2-[2-(2-{[2-(2,6-dioxopiperidin-3-yl)-1,3-dioxoisoindolin-4-yl]amino}ethoxy)ethoxy]ethoxy}propenamide (LLP037) 21*

*9-(2-{[2-(2,6-Dioxopiperidin-3-yl)-1,3-dioxoisoindolin-4-yl]oxy}acetamido)nonanoic acid (LLP050) 31*

Table of figures

### Figure S1: DH06 degradation profile

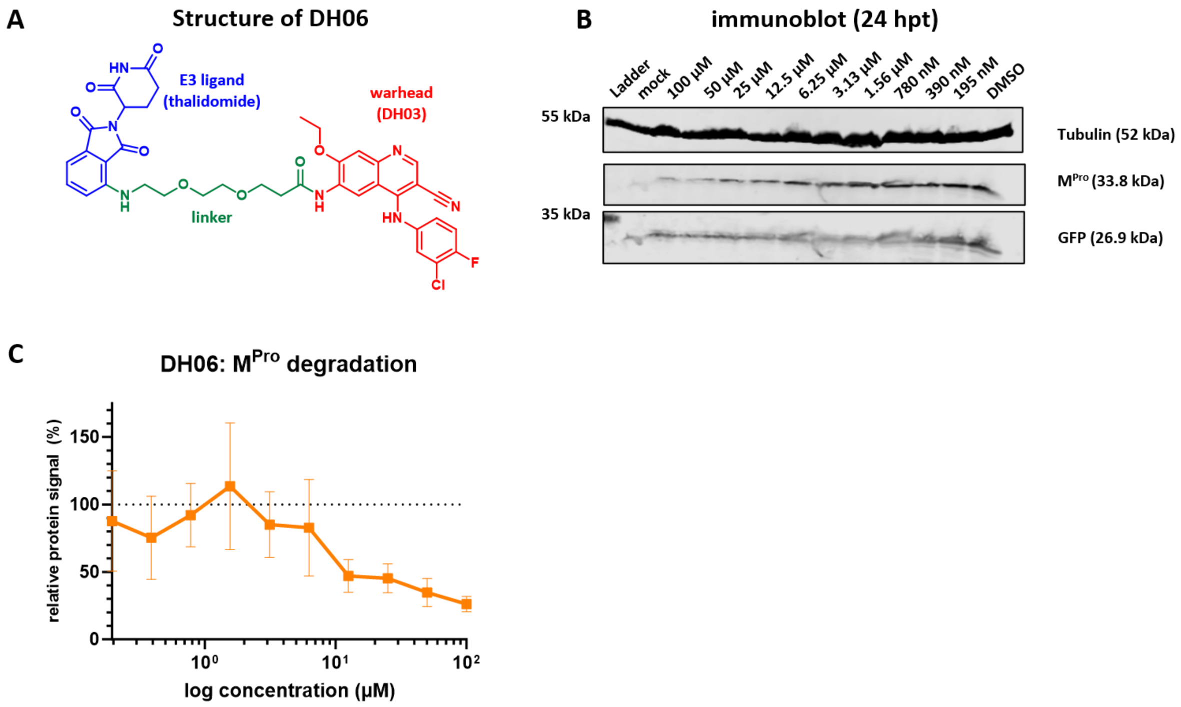

**Figure S1: DH06 is an M$^{Pro}$ degrader.** (**A**) DH06 consists of a flexible amino thalidomide exit group with a hydrophilic PEG-2 linker (green) that adds up to 10 C/N/O atoms between DH03 (red) and thalidomide (blue). (**B**) HEK293F cells were seeded in 6-well plates and medium was changed to DMEM++ with 3% FCS containing increasing concentrations of DH06 or DMSO (0.1%) prior transfection with 500 ng of each pCAGGS-M$^{Pro}$ and pCAGGS-GFP. At 24 hpt, cells were harvested and lysates subjected to SDS-PAGE and western blot. Representative image of three independent experiments. (**C**) Western blot signals of M$^{Pro}$ were quantified, normalized to tubulin and compared to DMSO.

Figure S2: Chemical structures of developed PROTACs

| | |
|---|---|
| **DH06**<br>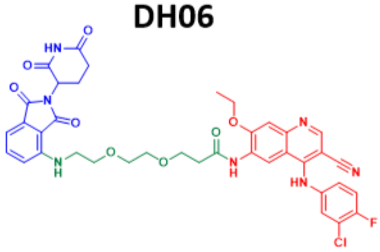 | **LLP041**<br>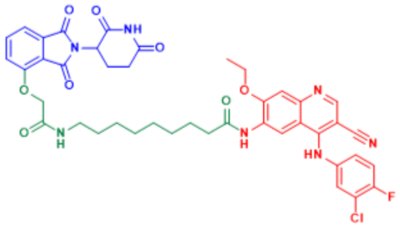 |
| **LLP019**<br>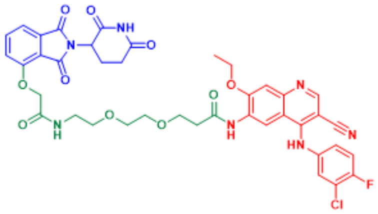 | **LP04**<br>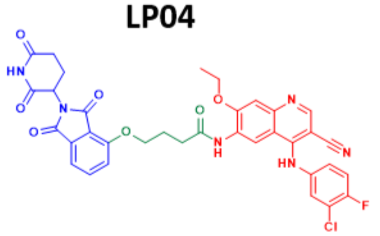 |
| **LLP031**<br>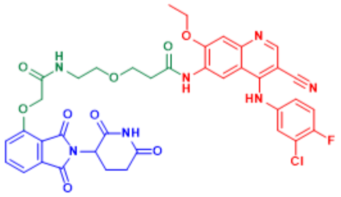 | **LP08**<br>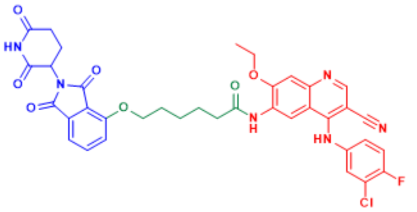 |
| **LLP037**<br>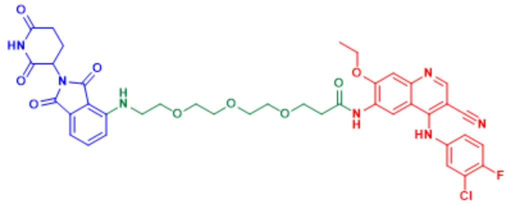 | **LP15**<br>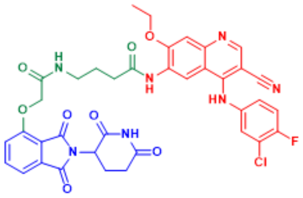 |
| **LLP038**<br>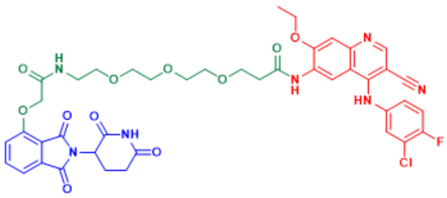 | **LB06**<br>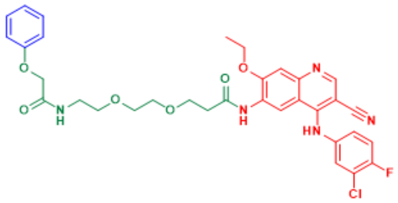 |
| **LLP049**<br>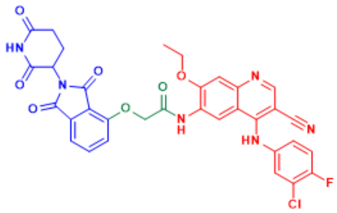 | |

**Figure S2: Chemical structures of developed PROTACs.**

Figure S3: $M^{Pro}$ degradation profiles of all PROTACs

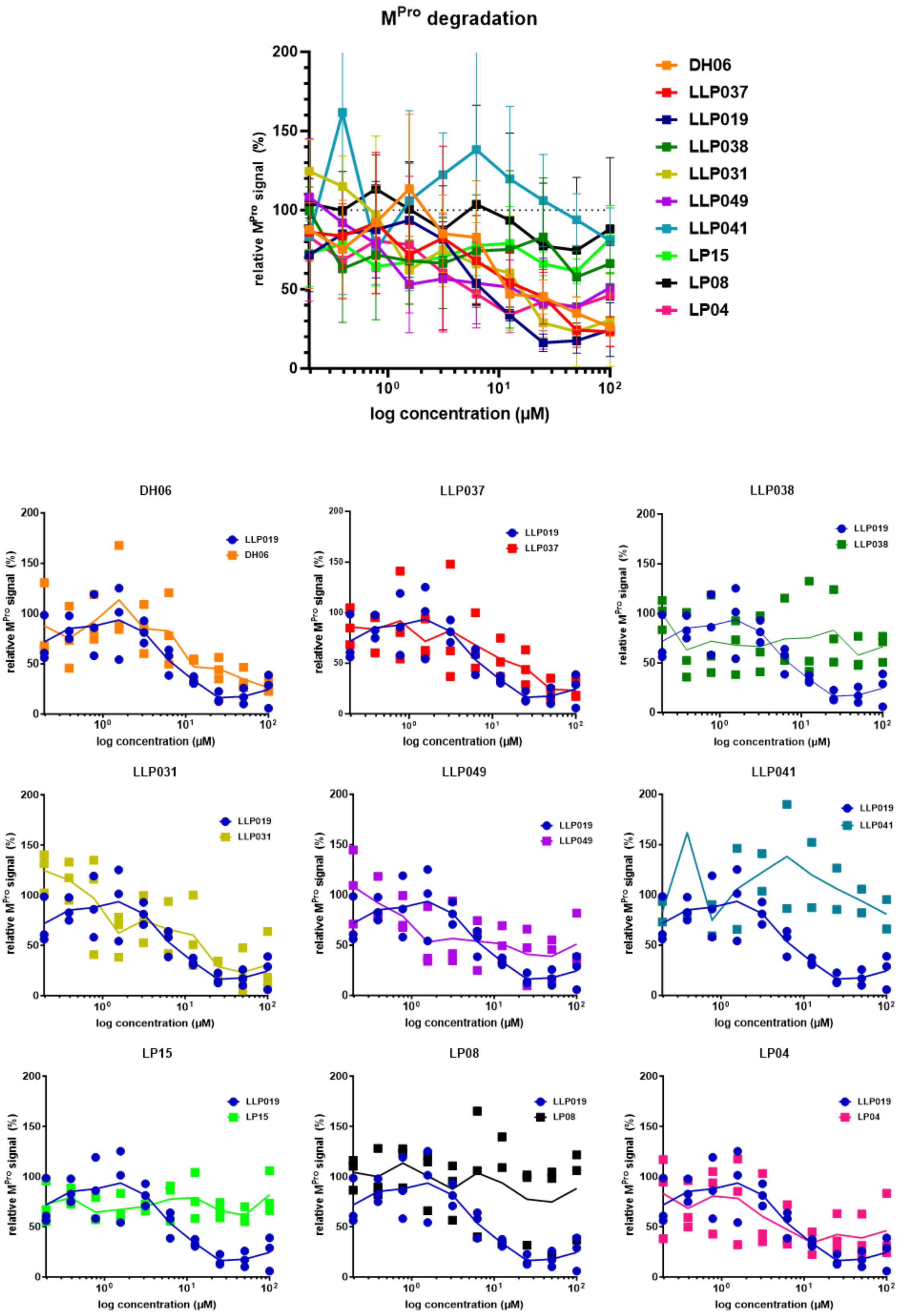

**Figure S3: $M^{Pro}$ degradation profiles of all PROTACs.** Cells were treated as described above in Figure S2 with all listed PROTACs and their degradation profile was compared to LLP019, as depicted below.

Figure S4: Western blots of the LLP019 mode of action confirmation

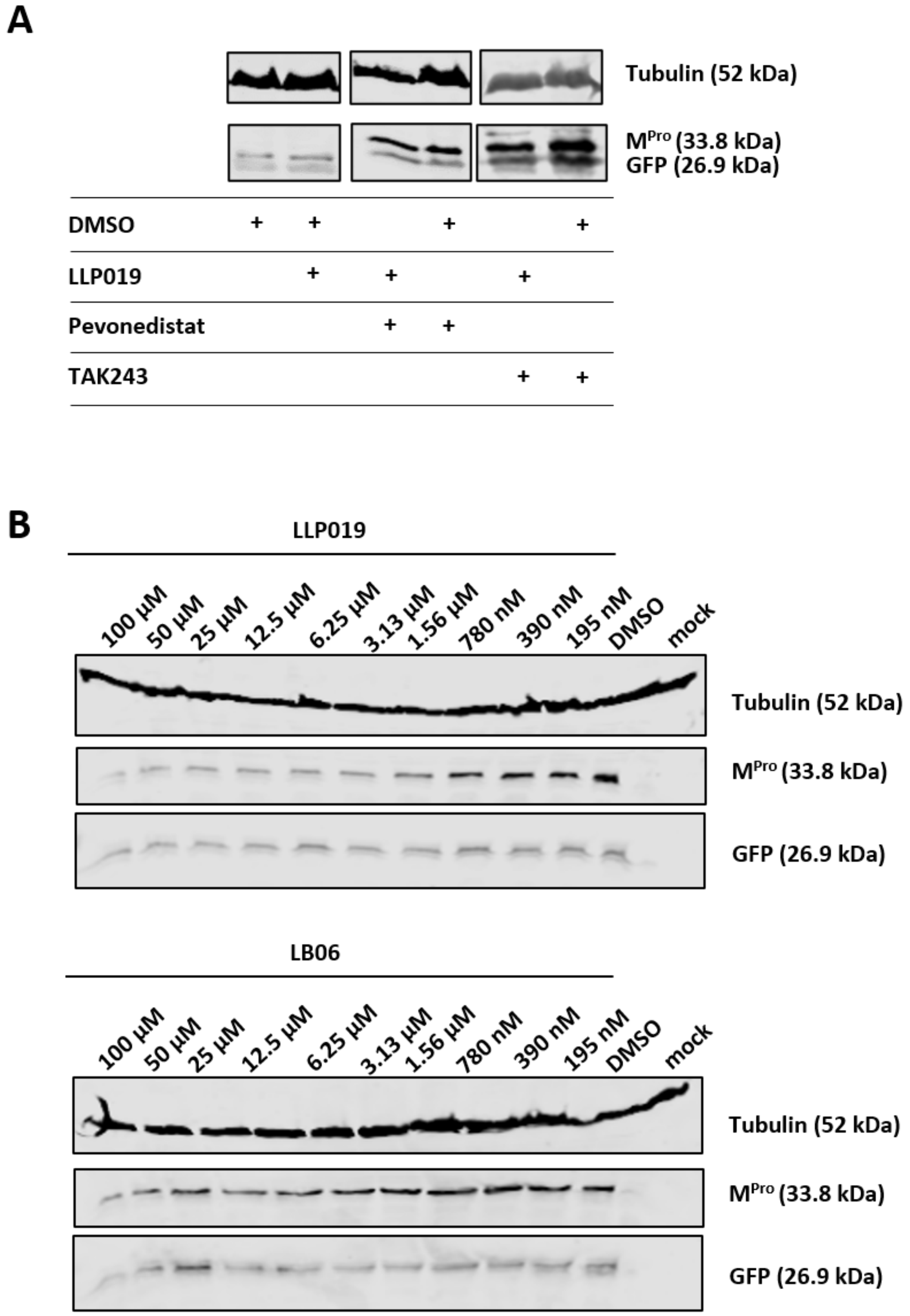

**Figure S4: Western blot of LLP019 or LB06 treated M$^{Pro}$-expressing HEK293F cells.** (**A**) HEK293F cells were transfected with 500 ng of each pCAGGS-M$^{Pro}$ and pCAGGS-GFP. At 24 hpt, transfection medium was changed to medium containing cycloheximide (10 µM) to stop protein expression in addition of either DMSO (0.1%), LLP019 (25 µM) or LLP019 (25 µM) plus pevonedistat (10 µM) or TAK243 (10 µM). After 5 h, cells were harvested and subjected to SDS-PAGE and western blot. Representative image of three independent experiments. (**B**) HEK293F cells were seeded in 6-well plates and transfection with 500 ng of each pCAGGS-M$^{Pro}$ and pCAGGS-GFP. After 4 h, medium was changed to DMEM with 3% FCS containing increasing concentrations of LLP019/LB06 or DMSO (0.1%) prior. At 24 hpt, cells were harvested and lysates subjected to SDS-PAGE and western blot. Representative image of three independent experiments.

Figure S5: Cytotoxic profiles of additional compounds

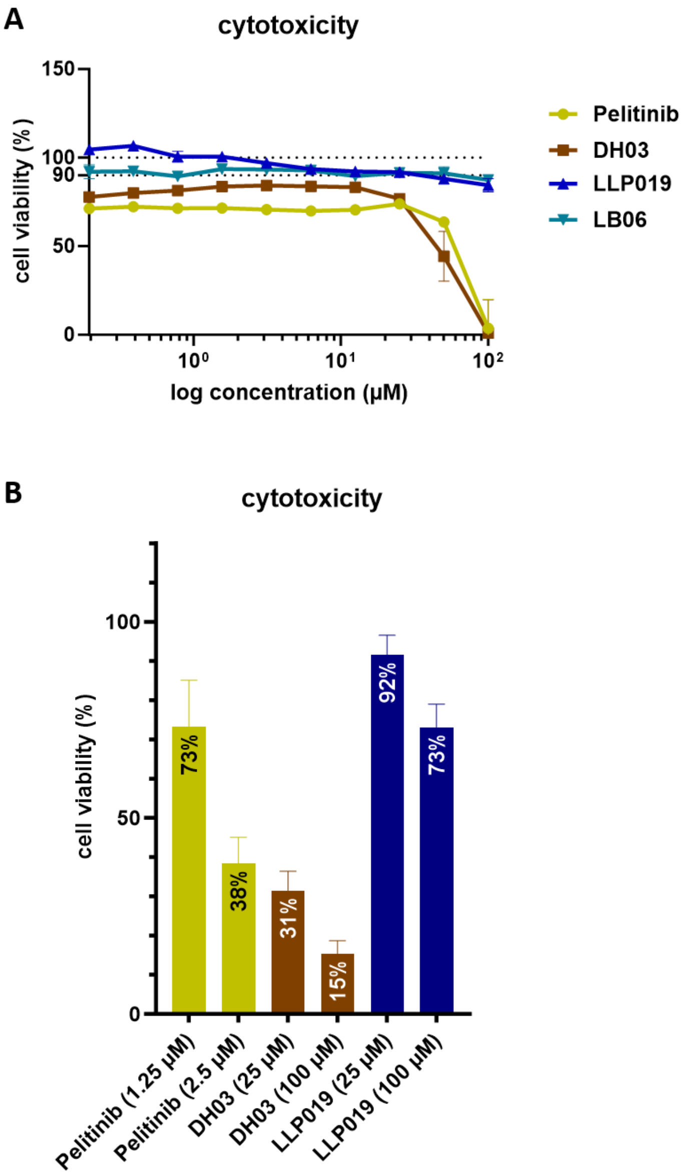

**Figure S5: Cytotoxicity of additional compounds.** (**A**) Calu3 cells were seeded in 96-well plates and incubated with medium containing pelitinib, DH03, LB06, LLP019, or DMSO (0.1% v/v). After 48 h, ATP-dependent cell viability was assessed using the CellTiter-Glo® 2.0 Cell Viability Assay by Promega and normalized to DMSO (set to 100%). (**B**) HEK293F cells were seeded in 96-well plates and incubated with medium containing pelitinib, DH03, LLP019 or DMSO (0.1%). After 48 h, NADH/NADPH-dependent cell viability was assessed using the CellTiter 96® AQueous One Solution Cell Proliferation Assay by Promega and normalized to DMSO (0.1%).

Figure S6: Structural alignment of SARS-CoV-2 and SARS-CoV M^Pro

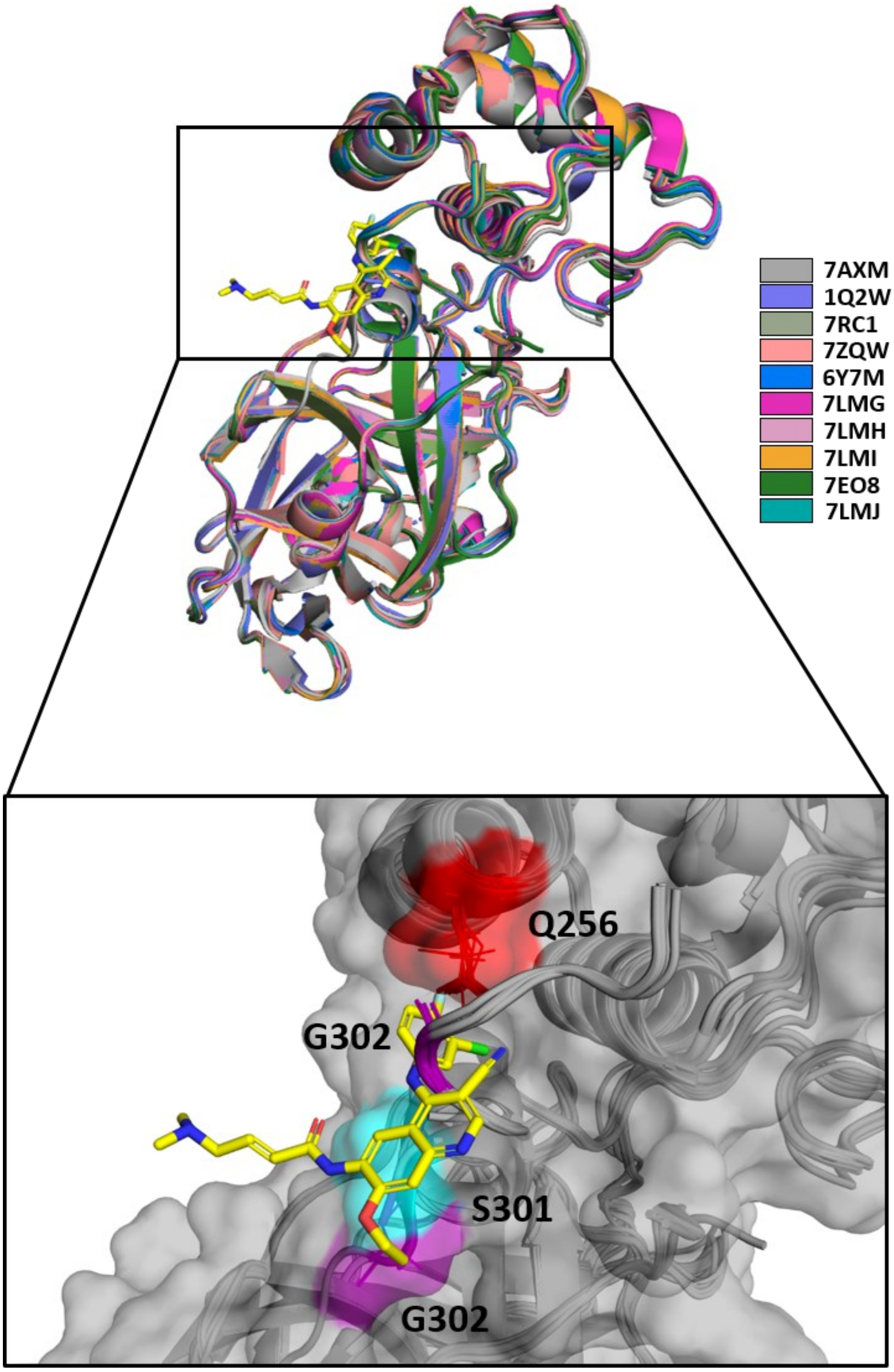

**Figure S6: Structural alignment of SARS-CoV-2 and SARS-CoV M^Pro.** Crystal structures of M^Pro were retrieved from the protein data bank. PDB-IDs of all used structures are listed in the figure and color-coded. All retrieved structures of SARS-CoV M^Pro were aligned to SARS-CoV-2 M^Pro (PDB-ID: 7AXM) using the PyMOL alignment tool.

Figure S7: Structural alignment of pelitinib's binding site in SARS-CoV-2, SARS-CoV, and MERS-CoV $M^{Pro}$

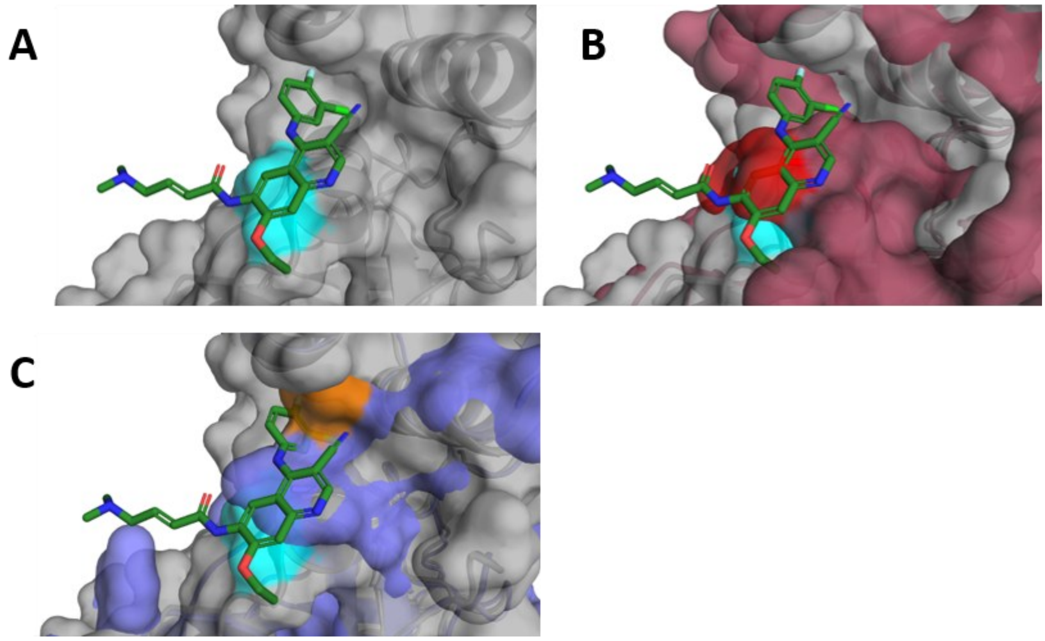

**Figure S7: Pelitinib's binding site is covered in SARS-CoV and MERS-CoV**. To understand the observed absence of antiviral activity of LLP019 against SARS-CoV and MERS-CoV, structural alignment of $M^{Pro}$ was performed using the PyMOL alignment tool. (**A**) Pelitinib's binding towards SARS-CoV-2 $M^{Pro}$ (PDB-ID: 7AXM, grey) is most likely facilitated by Ser301 (cyan). (**B**) At position 301, MERS-CoV (PDB-ID: 5C3N, bordeaux) contains a methionine (red), which most likely disrupts the binding site of pelitinib. (**C**) Similarly, SARS-CoV (PDB-ID: 1Q2W, pale blue) probably disrupts pelitinib's binding site via Glu256 (orange), which can be observed in numerous $M^{Pro}$ structures (Figure S6***: Structural alignment of SARS-CoV-2 and SARS-CoV $M^{Pro}$***

).

Scheme S1: Synthesis of LLP019

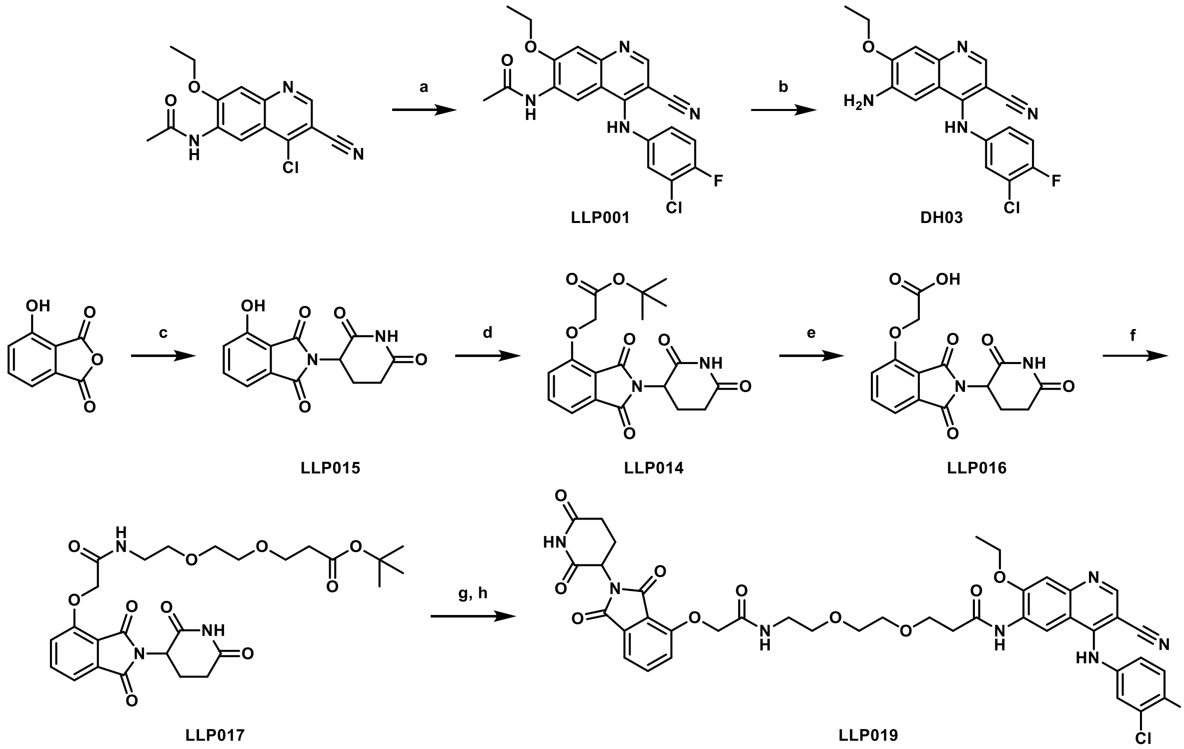

[a] Reagents and conditions: (a) 3-chloro-4-fluoroaniline, methane sulfonic acid, 2-propanol, reflux, 6.5 h, 84% [1]; (b) HCl conc., $H_2O$, reflux, overnight, 74% [2]; (c) 3-aminopiperidine-2,6-dione hydrochloride, KOAc, AcOH, reflux, 24 h, 77% [3]; (d) *tert*-butyl 2-bromoacetate, $K_2CO_3$, DMF, r.t., 2 h, 80%; (e) TFA, r.t., 4 h, 96% [4]; (f) *tert*-butyl 3-[2-(2-aminoethoxy)ethoxy]propanoate, HATU, DIPEA, DMF, r.t., 19 h, 67%; (g) TFA, r.t., 4 h (h) 6-amino-4-[(3-chloro-4-fluorophenyl)amino]-7-ethoxy-3-quinolinecarbonitrile, HATU, DIPEA, DMF, r.t., 19 h, 10%.

### *N*-{4-[(3-Chloro-4-fluorophenyl)amino]-3-cyano-7-ethoxyquinolin-6-yl}acetamide (LLP001)

*N*-(4-Chloro-3-cyano-7-ethoxy-6*N*-acetylquinoline) (4.00 g, 13.81 mmol, 1.00 eq) and 3-chloro-4-fluoroaniline (2.61 g, 17.95 mmol, 1.30 eq) were suspended in isopropanol (100.0 mL). After addition of methane sulfonic acid (0.06 mL, 0.83 mmol, 0.06 eq), the mixture was refluxed overnight. Subsequently, the resulting precipitate was filtered off and washed with 20% isopropanol in water and water. The product **LLP001** was obtained as a pale-yellow solid (4.59 g, 11.51 mmol) in 84% yield.

$^1$H NMR (500 MHz, DMSO-$d_6$, 300 K): δ (ppm) = 11.02 (bs, 1H), 9.58 (s, 1H), 9.05 (s, 1H), 8.98 (s, 1H), 7.74 (dd, $^4J_{H\text{-}F}$ = 6.6 Hz, $^4J$ = 2.6 Hz, 1H), 7.62 (s, 1H), 7.54 (dd, $^3J_{H\text{-}F}$ = 8.9 Hz, $^3J$ = 8.9 Hz, 1H), 7.46 (ddd, $^3J$ = 8.9 Hz, $^4J_{H\text{-}F}$ = 4.3 Hz, $^4J$ = 2.6 Hz, 1H), 4.33 (q, $^3J$ = 6.9 Hz, 2H), 2.20 (s, 3H), 1.50 (t, $^3J$ = 6.9 Hz, 3H).

$^{13}$C NMR (125 MHz, DMSO-$d_6$, 300 K): δ (ppm) = 169.0, 156.3 (d, $^1J_{C\text{-}F}$ = 247.1 Hz), 155.1, 153.4, 148.2, 138.2, 135.2 (d, $^4J_{C\text{-}F}$ = 2.4 Hz), 129.2, 128.1, 126.9 (d, $^3J_{C\text{-}F}$ = 7.2 Hz), 119.8 (d, $^2J_{C\text{-}F}$ = 19.2 Hz), 117.2 (d, $^2J_{C\text{-}F}$ = 22.8 Hz), 115.8, 114.3, 112.4, 101.8, 86.7, 65.3, 23.8, 14.0.

HR-MS: calculated for $C_{20}H_{16}ClFN_4O_2H$ $(M + H)^+$ 399.1019; found: 399.1005.

### 6-Amino-4-[(3-chloro-4-fluorophenyl)amino]-7-ethoxyquinoline-3-carbonitrile dihydrochloride (DH03)

**LLP001** (4.00 g, 10.03 mmol, 1.00 eq) was suspended in water (50.0 mL), concentrated hydrochloric acid (25.0 mL) added and the reaction mixture was refluxed overnight. Then, the pH was adjusted to pH 14 with NaOH pellets and the resulting precipitate was filtered off. The filter cake was washed with water and dried in a vacuum. **DH03** was obtained as a slightly impure yellow solid (3.58 g, 7.42 mmol) in 74% yield. A small amount was converted into its dihydrochloride to collect analytical data. The slightly impure material was used for further reactions.

$^{1}$H NMR (400 MHz, DMSO-$d_6$, 300 K): δ (ppm) = 10.98 (bs, 1H), 8.83 (s, 1H), 7.75 (dd, $^{4}J_{H\text{-}F}$ = 6.6 Hz, $^{4}J$ = 2.5 Hz, 1H), 7.68 (s, 1H), 7.58 (s, 1H), 7.50 (dd, $^{3}J_{H\text{-}F}$ = 8.9 Hz, $^{3}J$ = 8.9 Hz, 1H), 7.47 (ddd, $^{3}J$ = 8.9 Hz, $^{4}J_{H\text{-}F}$ = 4.4 Hz, $^{4}J$ = 2.5 Hz, 1H), 6.40 (bs, 4H), 4.26 (q, $^{3}J$ = 6.9 Hz, 2H), 1.48 (t, $^{3}J$ = 6.9 Hz, 3H).

$^{13}$C NMR (100 MHz, DMSO-$d_6$, 300 K): δ (ppm) = 156.3 (d, $^{1}J_{C\text{-}F}$ = 246.6 Hz), 153.0, 151.9, 143.3, 140.2, 135.3 (d, $^{4}J_{C\text{-}F}$ = 2.9 Hz), 132.4, 128.2, 127.0 (d, $^{3}J_{C\text{-}F}$ = 7.7 Hz), 119.7 (d, $^{2}J_{C\text{-}F}$ = 19.3 Hz), 117.2 (d, $^{2}J_{C\text{-}F}$ = 22.2 Hz), 114.35, 114.30, 101.9, 99.8, 85.9, 64.9, 14.1.

HR-MS: calculated for $C_{18}H_{14}ClFN_4OH$ $(M + H)^+$ 357.0913; found: 357.0888.

### 2-(2,6-Dioxopiperidin-3-yl)-4-hydroxyisoindoline-1,3-dione (LLP015)

3-Hydroxyphthalic anhydride (3.00 g, 18.28 mmol, 1.00 eq), 3-aminopiperidine-2,6-dione hydrochloride (3.31 g, 20.11 mmol, 1.10 eq) and KOAc (5.56 g, 56.67 mmol, 3.10 eq) were refluxed in acetic acid (60.0 mL) for 24 h. The mixture was added to ice-cold water (300 mL) and the resulting precipitate was subsequently filtered off. The filter cake was washed with cold water (3 x 70 mL) and then dried under high vacuum. The product **LLP015** was obtained as a gray solid (3.85 g, 14.04 mmol) in 77% yield.

$^{1}$H NMR (500 MHz, DMSO-$d_6$, 300 K): δ (ppm) = 11.14 (s, 1H), 11.06 (s, 1H), 7.65 (dd, $^{3}J$ = 8.3 Hz, $^{3}J$ = 7.2 Hz, 1H), 7.32 (dd, $^{3}J$ = 7.2 Hz, $^{4}J$ = 0.6 Hz, 1H), 7.25 (dd, $^{3}J$ = 8.3 Hz, $^{4}J$ = 0.6 Hz, 1H), 5.07 (dd, $^{3}J$ = 12.9 Hz, $^{3}J$ = 5.4 Hz, 1H), 2.88 (ddd, $^{2}J$ = 17.2 Hz, $^{3}J$ = 13.8 Hz, $^{3}J$ = 5.4 Hz, 1H), 2.65 - 2.46 (m, 2H), 2.02 (dddd, $^{2}J$ = 16.6 Hz, $^{3}J$ = 12.9 Hz, $^{3}J$ = 7.7 Hz, $^{3}J$ = 5.4 Hz, 1H).

$^{13}$C NMR (100 MHz, DMSO-$d_6$, 300 K): δ (ppm) = 172.7, 169.9, 167.0, 165.9, 155.4, 136.3, 133.1, 123.5, 114.4, 114.2, 48.6, 30.9, 22.0.

HR-MS: calculated for $C_{13}H_{10}N_2O_5Na$ $(M + Na)^+$ 297.0482; found: 297.0478.

### *tert*-Butyl 2-{[2-(2,6-dioxopiperidin-3-yl)-1,3-dioxoisoindolin-4-yl]oxy}acetate (LLP014)

**LLP015** (2.00 g, 7.29 mmol, 1.00 eq) was dissolved in DMF (73.0 mL), *tert*-butyl bromoacetate (1.08 mL, 7.29 mmol, 1.00 eq) and $K_2CO_3$ (1.51 g, 10.94 mmol, 1.50 eq) were added and the reaction mixture was stirred for 2 h at r.t. Then, ethyl acetate (30 mL) was added to the mixture and the organic phase was washed with 5% LiCl solution (30 mL). The aqueous layer was extracted with ethyl acetate (2 x 50 mL), the combined organic phase was again washed with 5% LiCl solution (5 x 20 mL), brine (20 mL) and finally dried over $MgSO_4$. The solvent was removed under reduced pressure, then the residue was adsorbed on silica gel and purified by column chromatography (cHex → cHex/EtOAc 0:100 over 30 min). The product **LLP014** was obtained as a white solid (2.27 g, 5.84 mmol) in 80% yield.

$^1$H NMR (500 MHz, DMSO-$d_6$, 300 K): δ (ppm) = 11.08 (s, 1H), 7.80 (dd, $^3J$ = 8.6 Hz, $^3J$ = 7.2 Hz, 1H), 7.48 (dd, $^3J$ = 7.2 Hz, $^4J$ = 0.6 Hz, 1H), 7.38 (dd, $^3J$ = 8.6 Hz, $^4J$ = 0.6 Hz, 1H), 5.10 (dd, $^3J$ = 12.9 Hz, $^3J$ = 5.4 Hz, 1H), 4.96 (s, 2H), 2.89 (ddd, $^2J$ = 17.2 Hz, $^3J$ = 13.8 Hz, $^3J$ = 5.4 Hz, 1H), 2.62 - 2.52 (m, 2H), 2.04 (dddd, $^2J$ = 16.0, $^3J$ = 12.9 Hz, $^3J$ = 7.7 Hz, $^3J$ = 5.4 Hz, 1H), 1.43 (s, 9H).

$^{13}$C NMR (125 MHz, DMSO-$d_6$, 300 K): δ (ppm) = 172.7, 169.8, 167.1, 166.7, 165.1, 155.0, 136.7, 133.2, 120.0, 116.5, 115.9, 81.9, 65.6, 47.8, 30.9, 27.6, 22.9.

HR-MS: calculated for $C_{19}H_{19}N_2O_7$ $(M - H)^-$ 387.1198; found: 387.1201.

### 2-{[2-(2,6-Dioxopiperidin-3-yl)-1,3-dioxoisoindolin-4-yl]oxy}acetic acid (LLP016)

**LLP014** (1.00 g, 2.57 mmol, 1.00 eq) was dissolved in TFA (26.0 mL, 332.9 mmol, 129.54 eq) and the mixture was stirred for 4 h at room temperature. Subsequently, DCM (20 mL) was added to the mixture and the solvent was removed under reduced pressure. After drying under high vacuum, the product **LLP016** was obtained as an off-white solid (0.82 g, 2.47 mmol) in 96% yield.

$^1$H NMR (500 MHz, DMSO-$d_6$, 300 K): δ (ppm) = 11.09 (s, 1H), 7.80 (dd, $^3J$ = 8.6 Hz, $^3J$ = 7.2 Hz, 1H), 7.47 (d, $^3J$ = 7.2 Hz, $^4J$ = 0.6 Hz, 1H), 7.39 (dd, $^3J$ = 8.6 Hz, $^4J$ = 0.6 Hz, 1H), 5.10 (dd, $^3J$ = 12.9 Hz, $^3J$ = 5.4 Hz, 1H), 4.98 (s, 2H), 2.89 (ddd, $^2J$ = 17.2 Hz, $^3J$ = 13.8 Hz, $^3J$ = 5.4 Hz, 1H), 2.62 - 2.52 (m, 2H), 2.05 (sm, 1H).

$^{13}$C NMR (125 MHz, DMSO-$d_6$, 300 K): δ (ppm) = 172.7, 169.8, 169.4, 166.7, 165.1, 155.1, 136.7, 133.2, 120.0, 116.3, 115.7, 65.1, 47.8, 30.9, 21.9.

HR-MS: calculated for $C_{15}H_{11}N_2O_7$ $(M - H)^-$ 331.0572; found: 311.0575.

### *tert*-Butyl 3-{2-[2-(2-{[2-(2,6-dioxopiperidin-3-yl)-1,3-dioxoisoindolin-4-yl]oxy}acetamido)ethoxy]ethoxy}propanoate (LLP017)

**LLP016** (0.70 g, 2.11 mmol, 1.00 eq) was dissolved in dry DMF (12.0 mL), mixed with *tert*-butyl 3-[2-(2-aminoethoxy)ethoxy]propanoate (0.54 mL, 2.32 mmol, 1.10 eq), HATU (1.20 g, 3.16 mmol, 1.50 eq) and *N*,*N*-diisopropylethylamine (1.10 mL, 6.32 mmol, 3.00 eq) and stirred under argon atmosphere overnight at r.t. Subsequently, ethyl acetate (50 mL) was added to the mixture and the organic phase was washed with 5% LiCl solution (aq., 30 mL). The aqueous layer was extracted with ethyl acetate (50 mL), the combined organic phase was then washed with 5% LiCl solution (aq., 5 x 30 mL), brine (30 mL) and finally dried over $MgSO_4$. The solvent was removed under reduced pressure, then the residue was adsorbed on silica gel and purified by column chromatography (cHex/EtOAc 70:30 → cHex/EtOAc 0:100 over 25 min). The product **LLP017** was obtained as a whitish solid (0.78 g, 1.42 mmol) in 67% yield.

$^{1}$H NMR (500 MHz, DMSO-$d_6$, 300 K): δ (ppm) = 11.09 (bs, 1H), 7.96 (t, $^{3}J$ = 5.7 Hz, 1H), 7.81 (dd, $^{3}J$ = 8.6 Hz, $^{3}J$ = 7.2 Hz, 1H), 7.50 (dd, $^{3}J$ = 7.2 Hz, $^{4}J$ = 0.6 Hz, 1H), 7.40 (dd, $^{3}J$ = 8.6 Hz, $^{4}J$ = 0.6 Hz, 1H), 5.11 (dd, $^{3}J$ = 5.4 Hz, $^{3}J$ = 12.9 Hz, 1H), 4.78 (s, 2H), 3.58 (t, $^{3}J$ = 6.3 Hz, 2H), 3.50 - 3.49 (m, 4H), 3.46 (t, $^{3}J$ = 5.4 Hz, 2H), 3.31 (sm, $^{3}J$ = 5.7 Hz, $^{3}J$ = 5.4 Hz, 2H), 2.90 (ddd, $^{2}J$ = 17.2 Hz, $^{3}J$ = 13.8 Hz, $^{3}J$ = 5.4 Hz, 1H), 2.63 - 2.52 (m, 2H), 2.41 (t, $^{3}J$ = 6.3 Hz, 2H), 2.04 (sm, 1H), 1.38 (s, 9H).

$^{13}$C NMR (125 MHz, DMSO-$d_6$, 300 K): δ (ppm) = 172.7, 170.4, 169.8, 166.8, 166.7, 165.4, 154.9, 136.9, 133.0, 120.4, 116.8, 116.0, 79.7, 69.6, 69.5, 68.8, 67.6, 66.2, 48.8, 38.4, 35.8, 30.9, 27.7, 22.0.

HR-MS: calculated for $C_{26}H_{32}N_3O_{10}$ (M - H)$^-$ 546.2093; found: 546.2101.

Purity was determined by qNMR using 1,2,4,5-tetrachloro-3-nitrobenzene as internal standard: 95.5%.

Mp: 92.1 °C

#### 3-{2-[2-(2-{[2-(2,6-Dioxopiperidin-3-yl)-1,3-dioxoisoindolin-4-yl]oxy}acetamido)ethoxy]ethoxy}propanoic acid (LLP023)

**LLP017** (1.20 g, 2.19 mmol, 1.00 eq) was dissolved in TFA (22.0 mL, 28.56 mmol, 130.39 eq) and the reaction mixture was stirred at r.t. for 4 h. DCM (20 mL) was added to the mixture. The solvent was removed under reduced pressure. After drying in a high vacuum, the product **LLP023** was obtained as a white solid (1.10 g, 2.19 mmol) in quantitative yield.

$^{1}$H NMR (500 MHz, DMSO-$d_6$, 300 K): δ (ppm) = 12.10 (bs, 1H), 11.09 (s, 1H), 7.98 (t, $^{3}J$ = 5.7 Hz, 1H), 7.81 (dd, $^{3}J$ = 8.6 Hz, $^{3}J$ = 7.2 Hz, 1H), 7.50 (dd, $^{3}J$ = 7.2 Hz, $^{4}J$ = 0.6 Hz, 1H), 7.40 (dd, $^{3}J$ = 8.6 Hz, $^{4}J$ = 0.6 Hz, 1H), 5.11 (dd, $^{3}J$ = 12.9 Hz, $^{3}J$ = 5.4 Hz, 1H), 4.79 (s, 2H), 3.59 (t, $^{3}J$ = 6.3 Hz, 2H), 3.51 - 3.49 (m, 4H), 3.46 (t, $^{3}J$ = 5.4 Hz, 2H), 3.31 (sm, 2H), 2.90 (ddd, $^{2}J$ = 17.2 Hz, $^{3}J$ = 13.8 Hz, $^{3}J$ = 5.4 Hz, 1H), 2.63 - 2.52 (m, 2H), 2.43 (t, $^{3}J$ = 6.3 Hz, 2H), 2.05 (sm, 1H).

$^{13}$C NMR (125 MHz, DMSO-$d_6$, 300 K): δ (ppm) = 172.7, 172.6, 169.8, 166.9, 166.7, 165.4, 155.0, 136.9, 133.0, 120.4, 116.8, 116.0, 69.5, 69.5, 68.8, 67.5, 66.2, 48.8, 38.4, 34.7, 30.9, 22.0.

HR-MS: calculated for $C_{22}H_{24}N_3O_{10}$ (M - H)$^-$ 490.1467; found: 490.1456.

Mp: 162.5 °C

*N*-{4-[(3-Chloro-4-fluorophenyl)amino]-3-cyano-7-ethoxyquinolin-6-yl}-3-{2-[2-(2-{[2-(2,6-dioxopiperidin-3-yl)-1,3-dioxoisoindolin-4-yl]oxy}acetamido)ethoxy]ethoxy}propanamide (LLP019)

**LLP023** (0.54 g, 1.10 mmol, 1.00 eq) was dissolved in dry DMF (11.0 mL), **DH03** (0.47 g, 1.32 mmol, 1.20 eq), HATU (0.63 g, 1.65 mmol, 1.50 eq) and *N*,*N*-diisopropylethylamine (0.58 mL, 3.30 mmol, 3.00 eq) were added and the resulting mixture was stirred under argon atmosphere overnight at r.t. Then, 5% LiCl solution (aq. 30 mL) was added and the mixture was extracted with ethyl acetate (2 x 100 mL). The combined organic phase was then washed with 5% LiCl solution (aq., 4 x 30 mL), brine (30 mL) and finally dried over $MgSO_4$. The solvent was removed under reduced pressure, then the residue was adsorbed on silica gel and purified by column chromatography (cHex/EtOAc 40:60 → 0:100 over 30 min; EtOAc → EtOAc/MeOH 90:10 over 15 min) and preparative HPLC ($H_2O$/MeCN 95:5 → 5:95 over 60 min). The product **LLP019** was obtained as a pale-yellow solid (90 mg, 0.11 mmol) in 10% yield.

$^{1}$H NMR (500 MHz, DMSO-*d$_6$*, 300 K): δ (ppm) = 11.10 (s, 1H), 9.90 (bs, 1H), 9.36 (s, 1H), 8.96 (s, 1H), 8.60 (s, 1H), 7.94 (t, $^3J$ = 5.7 Hz, 1H), 7.77 (dd, $^3J$ = 8.6 Hz, $^3J$ = 7.2 Hz, 1H), 7.47 (dd, $^4J_{H\text{-}F}$ = 6.6 Hz, $^4J$ = 2.6 Hz, 1H), 7.45 (d, $^3J$ = 7.2 Hz, 1H), 7.42 (dd, $^3J_{H\text{-}F}$ = 9.2 Hz, $^3J$ = 8.9 Hz, 1H), 7.38 (s, 1H), 7.36 (d, $^3J$ = 8.6 Hz, 1H), 7.25 (ddd, $^3J$ = 8.9 Hz, $^4J_{H\text{-}F}$ = 4.3 Hz, $^4J$ = 2.6 Hz, 1H), 5.10 (dd, $^3J$ = 12.9 Hz, $^3J$ = 5.4 Hz, 1H), 4.75 (s, 2H), 4.31 (q, $^3J$ = 7.2 Hz, 2H), 3.74 (t, $^3J$ = 6.0 Hz, 2H), 3.59 - 3.53 (m, 4H), 3.46 (t, $^3J$ = 5.4 Hz, 2H), 3.30 (sm, 2H), 2.89 (ddd, $^2J$ = 17.2 Hz, $^3J$ = 13.8 Hz, $^3J$ = 5.4 Hz, 1H), 2.72 (t, $^3J$ = 6.0 Hz, 2H), 2.62 - 2.51 (m, 2H), 2.05 (sm, 1H), 1.46 (t, $^3J$ = 7.2 Hz, 3H).

$^{13}$C NMR (125 MHz, DMSO-*d$_6$*, 300 K): δ (ppm) = 172.7, 169.8, 166.8, 166.6, 165.4, 154.9, 154.3 (d, $^1J_{C\text{-}F}$ = 243.5 Hz), 152.8, 151.6, 150.0, 147.2, 137.9 (d, $^4J_{C\text{-}F}$ = 3.6 Hz), 136.8, 132.9, 128.2, 124.5, 123.3 (d, $^3J_{C\text{-}F}$ = 7.2 Hz), 120.3, 119.5 (d, $^2J_{C\text{-}F}$ = 18.0 Hz), 117.0, 117.0 (d, $^2J_{C\text{-}F}$ = 21.6 Hz), 116.7, 116.0, 113.9, 113.5, 108.3, 89.0, 69.6, 69.5, 68.8, 67.5, 66.5, 64.7, 48.8, 38.4, 37.0, 30.9, 22.0, 14.2.

HR-MS: calculated for $C_{40}H_{36}ClFN_7O_{10}$ $(M - H)^-$ 828.2202; found: 828.2214.

Elemental analysis calculated (%) for $C_{40}H_{37}ClN_7O_{10}$ x 2.00$H_2O$: N: 11.32, C: 55.46, H: 4.77; found: N: 11.17, C: 55.73, H: 5.02.

Mp: 174.0 °C

Scheme S2: Synthesis of DH06[a]

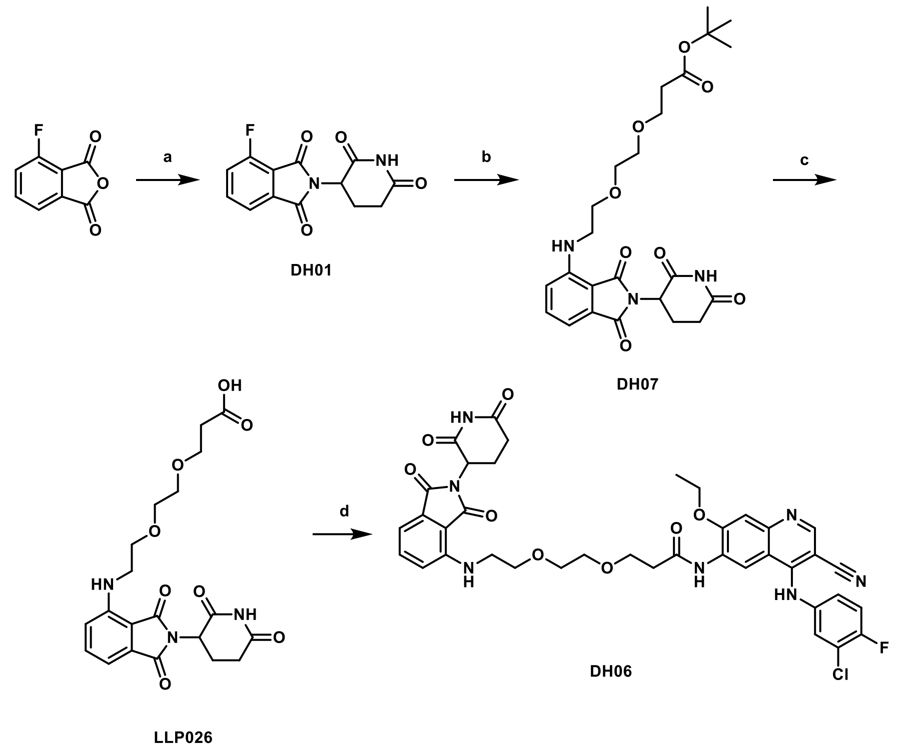

[a] Reagents and conditions: (a) 3-aminopiperidine-2,6-dione hydrochloride, NaOAc, AcOH, reflux, 12 h, 77% [5]; (b) *tert*-butyl 3-[2-(2-aminoethoxy)ethoxy]propanoate, DIPEA, NMP, 90 °C, overnight, 28% [6]; (c) TFA, DCM, rt, 3 h, 95%[6]; (d) **DH03**, HATU, DIPEA, DMF, r.t., overnight, 11%.

### 2-(2,6-Dioxopiperidin-3-yl)-4-fluoroisoindoline-1,3-dione (DH01)

3-Fluorophthalic anhydride (1.00 g, 6.02 mmol, 1.00 eq), 3-aminopiperidine-2,6-dione hydrochloride (0.99 g, 6.02 mmol, 1.00 eq) and NaOAc (0.60 g, 7.20 mmol, 1.20 eq) were dissolved in acetic acid (100.0 mL) and refluxed for 12 h. After cooling to r.t., the reaction solution was poured into water (400 mL) and the aqueous phase was extracted with EtOAc (4 x 150 mL). The combined organic phases were dried over $MgSO_4$, filtered and the solvent was removed under reduced pressure. The remaining residue was adsorbed on silica gel and purified by column chromatography (DCM → DCM/MeOH 90:10 over 20 min). The product **DH01** was obtained as a gray solid (1.28 g, 4.63 mmol) in 77% yield.

$^{1}$H NMR (500 MHz, DMSO-$d_6$, 300 K): δ (ppm) = 11.12 (s, 1H), 7.95 (ddd, $^{3}J$ = 7.3 Hz, $^{3}J$ = 8.4 Hz, $^{4}J_{H-F}$ = 4.6 Hz, 1H), 7.79 (d, $^{3}J$ = 7.5 Hz, 1H), 7.73 (dd, $^{3}J_{H-F}$ = 9.2 Hz, $^{3}J$ = 8.6 Hz, 1H), 5.16 (dd, $^{3}J$ = 5.4 Hz, $^{3}J$ = 12.9 Hz, 1H), 2.89 (ddd, $^{2}J$ = 17.2 Hz, $^{3}J$ = 13.8 Hz, $^{3}J$ = 5.4 Hz, 1H), 2.61 (ddd, $^{2}J$ = 17.2 Hz, $^{3}J$ = 4.6 Hz, $^{3}J$ = 4.3 Hz, 1H), 2.57 - 2.48 (m, 1H), 2.07 (dddd, $^{2}J$ = 16.0 Hz, $^{3}J$ = 12.9 Hz, $^{3}J$ = 7.7 Hz, $^{3}J$ = 5.4 Hz, 1H).

$^{13}$C NMR (125 MHz, DMSO-$d_6$, 300 K): δ (ppm) = 172.6, 169.6, 166.0 (d, $^{3}J_{C-F}$ = 2.4 Hz), 163.9, 156.8 (d, $^{1}J_{C-F}$ = 262.7 Hz), 138.0 (d, $^{3}J_{C-F}$ = 7.2 Hz), 133.4, 122.9 (d, $^{2}J_{C-F}$ = 19.2 Hz), 120.0 (d, $^{4}J_{C-F}$ = 2.4 Hz), 117.0 (d, $^{2}J_{C-F}$ = 13.2 Hz), 49.1, 30.9, 21.8.

HR-MS: calculated for $C_{13}H_9FN_2O_4H$ (M + H)$^+$: 277.0619; found: 277.0597.

### *tert*-Butyl 3-[2-(2-{[2-(2,6-dioxopiperidin-3-yl)-1,3-dioxoisoindolin-4-yl]amino}ethoxy)ethoxy]propanoate (DH07)

**DH01** (1.20 g, 4.34 mmol, 1.00 eq) was dissolved in NMP (12.0 mL), *tert*-butyl 3-[2-(2-aminoethoxy)ethoxy]propanoate (1.11 mL, 4.78 mmol, 1.10 eq) and *N*,*N*-diisopropylethylamine (1.51 mL, 8.68 mmol, 2.00 eq) were added and the reaction mixture stirred overnight at 90 °C. After cooling to room temperature, ethyl acetate (40 mL) was added, and the mixture was washed with water (4 x 10 mL), brine (10 mL) and finally the organic phase was dried over $MgSO_4$. The solvent was removed under reduced pressure, then the residue was adsorbed on silica gel and purified by column chromatography (cHex/EtOAc 70:30 → cHex/EtOAc 0:100 over 30 min). The product **DH07** was obtained as a green, highly viscous oil (600 mg, 1.23 mmol) in 28% yield.

$^1$H NMR (500 MHz, DMSO-$d_6$, 300 K): δ (ppm) = 11.07 (s, 1H), 7.58 (dd, $^3J$ = 8.6 Hz, $^3J$ = 7.2 Hz, 1H), 7.14 (d, $^3J$ = 8.6 Hz, 1H), 7.04 (d, $^3J$ = 7.2 Hz, 1H), 6.59 (t, $^3J$ = 5.7 Hz, 1H), 5.05 (dd, $^3J$ = 12.9 Hz, $^3J$ = 5.4 Hz, 1H), 3.62 (t, $^3J$ = 5.4 Hz, 2H), 3.59 (t, $^3J$ = 6.3 Hz, 2H), 3.56 - 3.54 (m, 2H), 3.52 - 3.50 (m, 2H), 3.46 (sm, 2H), 2.88 (ddd, $^2J$ = 17.2 Hz, $^3J$ = 13.8 Hz, $^3J$ = 5.4 Hz, 1H), 2.61 - 2.51 (m, 2H), 2.39 (t, $^3J$ = 6.3 Hz, 2H), 2.03 (sm, 1H), 1.38 (s, 9H).

$^{13}$C NMR (125 MHz, DMSO-$d_6$, 300 K): δ (ppm) = 172.7, 170.3, 170.0, 168.9, 167.2, 146.4, 136.2, 132.1, 117.4, 110.6, 109.3, 79.6, 69.7, 68.9, 66.3, 48.5, 41.7, 35.8, 30.9, 27.7, 22.1

HR-MS: calculated for $C_{24}H_{31}N_3O_8Na$ (M + Na)$^+$: 512.1983; found: 512.2003.

Purity was determined by qNMR using 1,2,4,5-tetrachloro-3-nitrobenzene as internal standard: 95.6%.

### 3-[2-(2-{[2-(2,6-Dioxopiperidin-3-yl)-1,3-dioxoisoindolin-4-yl]amino}ethoxy)ethoxy]propanoic acid (LLP026)

**DH07** (0.20 g, 0.41 mmol, 1.00 eq) was dissolved in DCM (10.0 mL) and TFA (0.63 mL, 8.17 mmol, 20.00 eq) was added. The mixture was stirred at room temperature for 3 h. The product **LLP026** (0.17 g, 0.39 mmol) was obtained as a yellow solid in 95% yield by removing the solvent and subsequent drying under high vacuum.

$^1$H NMR (500 MHz, DMSO-$d_6$, 300 K): δ (ppm) = 11.06 (s, 1H), 7.58 (dd, $^3J$ = 8.6 Hz, $^3J$ = 6.9 Hz, 1H), 7.14 (d, $^3J$ = 8.6 Hz, 1H), 7.04 (d, $^3J$ = 6.9 Hz, 1H), 6.60 (bs, 1H), 5.05 (dd, $^3J$ = 12.9 Hz, $^3J$ = 5.4 Hz, 1H), 3.63 - 3.59 (m, 4H), 3.56 - 3.54 (m, 2H), 3.52 - 3.51 (m, 2H), 3.48 - 3.45 (sm, 2H), 2.88 (ddd, $^2J$ = 17.2 Hz, $^3J$ = 13.8 Hz, $^3J$ = 5.4 Hz, 1H), 2.62 - 2.51 (m, 2H), 2.43 (t, $^3J$ = 6.3 Hz, 2H), 2.06 - 2.00 (sm, 1H).

$^{13}C$ NMR (125 MHz, DMSO-$d_6$, 300 K): δ (ppm) = 172.7, 170.3, 170.0, 168.9, 167.2, 146.4, 136.2, 132.1, 117.4, 110.6, 109.3, 79.6, 69.7, 68.9, 66.3, 48.5, 41.7, 35.8, 30.9, 27.7, 22.1.

HR-MS: calculated for $C_{20}H_{22}N_3O_8$ $(M - H)^-$: 432.1412; found: 432.1407.

Mp: 97.7 °C

### *N*-{4-[(3-Chloro-4-fluorophenyl)-amino]-3-cyano-7-ethoxyquinolin-6-yl}-3-[2-(2-{[2-(2,6-dioxopiperidin-3-yl)-1,3-dioxoisoindolin-4-yl]amino}ethoxy)-ethoxy]propenamide (DH06)

**LLP026** (0.36 g, 0.84 mmol, 1.00 eq) was dissolved in dry DMF (5.0 mL), **DH03** (0.30 g, 0.84 mmol, 1.00 eq), HATU (0.64 g, 1.68 mmol, 2.00 eq) and *N*,*N*-diisopropylethylamine (0.74 mL, 0.44 mmol, 5.20 eq) were added and the reaction mixture stirred under an argon atmosphere at r.t. overnight. After this time, ethyl acetate (40 mL) was added, and the mixture was successively washed with 5% LiCl solution (aq., 4 x 15 mL), brine (15 mL), and finally the organic phase was dried over $MgSO_4$. The solvent was removed under reduced pressure, the remaining residue was adsorbed on silica gel and purified by column chromatography (cHex → cHex/EtOAc 0:100 over 30 min; EtOAc → EtOAc/MeOH 90:10 over 10 min). The product **DH06** was obtained as a yellow solid (68 mg, 0.09 mmol) in 11% yield.

$^1H$ NMR (500 MHz, DMSO-$d_6$, 300 K): δ (ppm) = 11.07 (s, 1H), 9.66 (s, 1H), 9.33 (s, 1H), 8.94 (s, 1H), 8.53 (s, 1H), 7.52 (dd, $^3J$ = 8.6 Hz, $^3J$ = 7.2 Hz, 1H), 7.42 - 7.36 (m, 3H), 7.22 - 7.19 (m, 1H), 7.05 (d, $^3J$ = 8.6 Hz, 1H), 6.99 (d, $^3J$ = 7.2 Hz, 1H), 6.55 (t, $^3J$ = 5.7 Hz, 1H), 5.03 (dd, $^3J$ = 12.9 Hz, $^3J$ = 5.4 Hz, 1H), 4.30 (q, $^3J$ = 7.2 Hz, 2H), 3.75 (t, $^3J$ = 6.0 Hz, 2H), 3.61 (t, $^3J$ = 5.4 Hz, 2H), 3.60 (ps, 4H, 22-$H_2$), 3.43 - 3.39 (sm, 2H), 2.86 (ddd, $^2J$ = 17.2 Hz, $^3J$ = 13.8 Hz, $^3J$ = 5.4 Hz, 1H), 2.71 (t, $^3J$ = 6.0 Hz, 2H), 2.59 - 2.53 (m, 2H), 2.05 - 2.00 (sm, 1H), 1.45 (t, $^3J$ = 7.2 Hz, 3H).

$^{13}C$ NMR (125 MHz, DMSO-$d_6$, 300 K): δ (ppm) = 172.7, 170.0, 169.8, 168.9, 167.2, 154.2 (d, $^1J_{C-F}$ = 243.5 Hz), 152.7, 151.7, 149.9, 147.5, 146.3, 137.9 (d, $^4J_{C-F}$ = 3.6 Hz), 136.1, 132.0, 128.2, 124.4, 123.2 (d, $^3J_{C-F}$ = 6.0 Hz), 119.5 (d, $^2J_{C-F}$ = 18.0 Hz), 117.3, 117.1, 117.0 (d, $^2J_{C-F}$ = 24.0 Hz), 113.9, 113.4, 110.6, 109.2, 108.5, 89.1, 69.6, 69.6, 68.9, 66.5, 64.6, 48.5, 41.7, 37.0, 30.9, 22.1, 14.2.

HR-MS: calculated for $C_{38}H_{35}ClFN_7O_8Na$ $(M + Na)^+$: 794.2107; found: 794.2112.

Elemental analysis calculated (%) for $C_{38}H_{35}ClN_7O_8$ x 1.00 $H_2O$: N: 12.41, C: 57.76, H: 4.72; found: N: 12.26, C: 57.54, H: 5.11.

Mp: 154.7 °C

Scheme S3: Synthesis of LLP037[a]

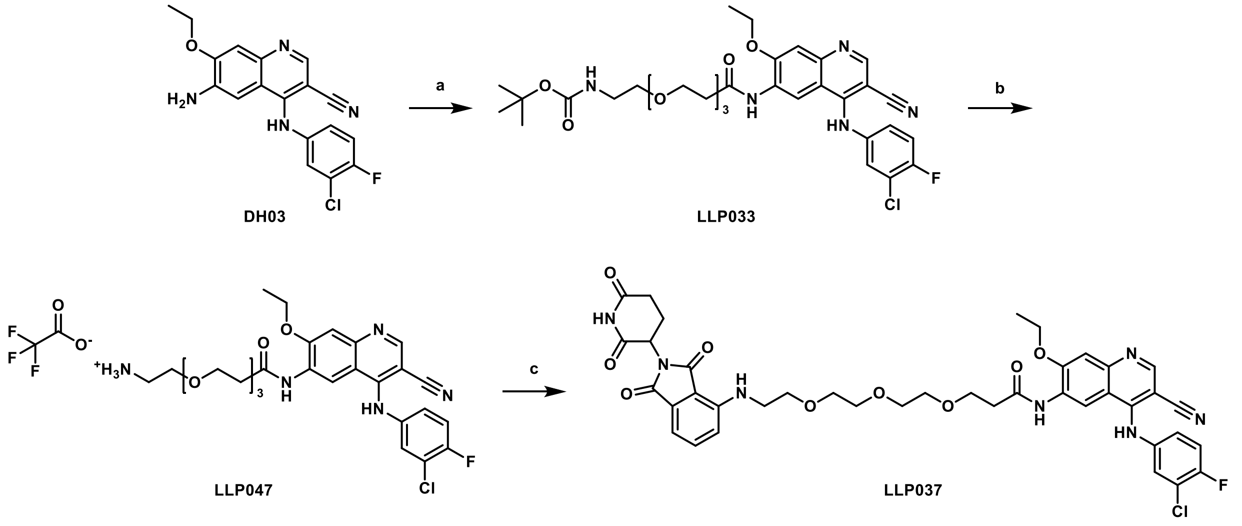

[a] Reagents and conditions: (a) 12-[(*tert*-butoxycarbonyl)amino]-4,7,10-trioxadodecanoic acid, HATU, DIPEA, DMSO, r.t., overnight, 41%; (b) TFA, DCM, r.t., 2 h, 100%; (c) **DH01**, DIPEA, NMP, 90 °C, overnight, 18%.

### *tert*-Butyl [2-(2-{2-[3-({4-[(3-chloro-4-fluorophenyl)amino]-3-cyano-7-ethoxyquinolin-6-yl}amino)-3-oxopropoxy]ethoxy}ethoxy)ethyl]carbamate (LLP033)

**DH03** (0.49 g, 1.37 mmol, 1.10 eq) was dissolved in dry DMSO (12.0 mL), mixed with 12-[(*tert*-butoxycarbonyl)amino]-4,7,10-trioxadodecanoic acid (0.40 g, 1.24 mmol, 1.00 eq), HATU (0.71 g, 1.86 mmol, 1.50 eq) and *N*,*N*-diisopropylethylamine (0.65 mL, 3.72 mmol, 3.00 eq) and stirred overnight at r.t. After this time, EtOAc (70 mL) was added, the reaction mixture was washed with water (30 mL) and subsequently, the aqueous phase was extracted with EtOAc (2 x 50 mL). The combined organic phase was washed with water (3 x 30 mL), brine (30 mL) and dried over $MgSO_4$. The solvent was removed under reduced pressure, then the residue was adsorbed on silica gel and purified by column chromatography (cHex/EtOAc 50:50 → cHex/EtOAc 0:100 over 20 min; EtOAc → EtOAc/MeOH 90:10 over 18 min). The product **LLP033** was obtained as a yellow solid (0.33 g, 0.50 mmol) in 41% yield.

$^1$H NMR (500 MHz, DMSO-$d_6$, 300 K): δ (ppm) = 9.76 (s, 1H), 9.36 (s, 1H), 9.00 (s, 1H), 8.56 (s, 1H), 7.44 (dd, $^4J_{H\text{-}F}$ = 6.3 Hz, $^4J$ = 2.9 Hz, 1H), 7.41 (s, 1H), 7.40 (dd, $^3J_{H\text{-}F}$ = 9.2 Hz, $^3J$ = 8.9 Hz, 1H), 7.23 (ddd, $^3J$ = 8.9 Hz, $^4J_{H\text{-}F}$ = 4.3 Hz, $^4J$ = 2.9 Hz, 1H), 6.69 (t, $^3J$ = 6.0 Hz, 1H), 4.32 (q, $^3J$ = 6.9 Hz, 2H), 3.74 (t, $^3J$ = 6.0 Hz, 2H), 3.58 - 3.45 (m, 8H), 3.34 (t, $^3J$ = 6.0 Hz, 2H), 3.03 (sm, 2H), 2.73 (t, $^3J$ = 6.0 Hz, 2H), 1.47 (t, $^3J$ = 6.9 Hz, 3H), 1.36 (s, 9H).

$^{13}$C NMR (125 MHz, DMSO-$d_6$, 300 K): δ (ppm) = 169.8, 155.5, 154.3 (d, $^1J_{C\text{-}F}$ = 238.7 Hz), 152.9, 151.6, 150.1, 147.0, 137.9 (d, $^4J_{C\text{-}F}$ = 3.6 Hz), 128.2, 124.6, 123.3 (d, $^3J_{C\text{-}F}$ = 7.2 Hz), 119.5 (d, $^2J_{C\text{-}F}$ = 19.2 Hz),

117.0 (d, $^{2}J_{C-F}$ = 22.8 Hz), 117.0, 113.9, 113.6, 108.1, 89.0, 77.5, 69.7, 69.7, 69.6, 69.4, 69.1, 66.5, 64.7, 39.5, 37.0, 28.2, 14.2.

HR-MS: calculated for $C_{32}H_{39}ClFN_5O_7H$ $(M + H)^+$: 660.2595; found: 660.2585.

Purity was determined by qNMR using dimethyl terephthalate as internal standard: 88.7%.

Mp: 61.8 °C

### 2-(2-{2-[3-({4-[(3-Chloro-4-fluorophenyl)amino]-3-cyano-7-ethoxyquinolin-6-yl}amino)-3-oxopropoxy]ethoxy}ethoxy)ethan-1-aminium 2,2,2-trifluoroacetate (LLP047)

**LLP033** (0.40 g, 0.61 mmol, 1.00 eq) was dissolved in DCM (10.0 mL), TFA (0.93 mL, 12.12 mmol, 20.00 eq) was added and the mixture was stirred at r.t. for 2 h. Removal of the solvent followed by drying under high vacuum rendered **LLP047** (0.41 g, 0.61 mmol) as an orange oil in quantitative yield.

$^{1}$H NMR (500 MHz, DMSO-*d*$_6$, 300 K): δ (ppm) = 9.76 (s, 1H), 9.38 (s, 1H), 8.94 (s, 1H), 8.56 (s, 1H), 7.83 (s, 3H), 7.44 (dd, $^{4}J_{H-F}$ = 6.3 Hz, $^{4}J$ = 2.9 Hz, 1H), 7.42 (s, 1H), 7.41 (dd, $^{3}J_{H-F}$ = 9.2 Hz, $^{3}J$ = 8.9 Hz, 1H), 7.23 (ddd, $^{3}J$ = 8.9 Hz, $^{4}J_{H-F}$ = 4.3 Hz, $^{4}J$ = 2.9 Hz, 1H), 4.32 (q, $^{3}J$ = 6.9 Hz, 2H), 3.74 (t, $^{3}J$ = 6.0 Hz, 2H), 3.57 - 3.54 (m, 10H), 3.03 (ps, 2H), 2.73 (t, $^{3}J$ = 6.0 Hz, 2H), 1.47 (t, $^{3}J$ = 6.9 Hz, 3H).

$^{13}$C NMR (125 MHz, DMSO-*d*$_6$, 300 K): δ (ppm) = 169.8, 158.1 (q, $^{2}J_{C-F}$ = 31.2 Hz), 154.3 (d, $^{1}J_{C-F}$ = 238.7 Hz), 153.0, 151.6, 150.1, 147.0, 138.0 (d, $^{4}J_{C-F}$ = 3.6 Hz), 128.2, 124.6, 123.3 (d, $^{3}J_{C-F}$ = 7.2 Hz), 119.5 (d, $^{2}J_{C-F}$ = 19.2 Hz), 117.2 (q, $^{1}J_{C-F}$ = 300.0 Hz), 117.0 (d, $^{2}J_{C-F}$ = 22.8 Hz), 116.9, 113.9, 113.8, 108.1, 89.0, 69.7, 69.6, 69.6, 66.6, 66.5, 64.7, 38.5, 36.9, 14.2.

HR-MS: calculated for $C_{27}H_{32}ClFN_5O_5$ $(M)^+$: 560.2071; found: 560.2047.

### *N*-{4-[(3-Chloro-4-fluorophenyl)amino]-3-cyano-7-ethoxyquinolin-6-yl}-3-{2-[2-(2-{[2-(2,6-dioxopiperidin-3-yl)-1,3-dioxoisoindolin-4-yl]amino}ethoxy)ethoxy]ethoxy}propenamide (LLP037)

**LLP047** (0.25 g, 0.37 mmol, 1.00 eq) was dissolved in NMP (3.0 mL), **DH01** (0.12 g, 0.45 mmol, 1.20 eq) and *N*,*N*-diisopropylethylamine (0.20 mL, 1.11 mmol, 3.00 eq) were added and the mixture stirred overnight at 90 °C. After this time, EtOAc (30 mL) was added and the mixture was washed with water (2 x 10 mL). The combined aqueous phase was extracted with ethyl acetate (2 x 20 mL). The combined organic phase was washed with brine (10 mL) and dried over $MgSO_4$. The solvent was removed under reduced pressure, then the residue was adsorbed on silica gel and purified first by column chromatography (DCM → DCM/MeOH 90:10 over 30 min) and then by preparative HPLC ($H_2O$/MeCN 95:5 → 5:95 over 60 min). The product **LLP037** was obtained as a yellow solid (53 mg, 0.06 mmol) in 18% yield.

$^1H$ NMR (500 MHz, DMSO-$d_6$, 300 K): δ (ppm) = 11.07 (s, 1H), 9.67 (s, 1H), 9.33 (s, 1H), 8.95 (s, 1H), 8.53 (s, 1H), 7.54 (dd, $^3J$ = 8.6 Hz, $^3J$ = 7.2 Hz, 1H), 7.42 - 7.37 (m, 3H), 7.23 - 7.20 (m, 1H), 7.08 (d, $^3J$ = 8.6 Hz, 1H), 7.01 (d, $^3J$ = 7.2 Hz, 1H), 6.55 (t, $^3J$ = 5.7 Hz, 1H), 5.04 (dd, $^3J$ = 12.9 Hz, $^3J$ = 5.4 Hz, 1H), 4.31 (q, $^3J$ = 6.9 Hz, 2H), 3.73 (t, $^3J$ = 6.0 Hz, 2H), 3.59 - 3.53 (m, 10H), 3.41 (sm, 2H), 2.87 (ddd, $^2J$ = 17.2 Hz, $^3J$ = 13.8 Hz, $^3J$ = 5.4 Hz, 1H), 2.71 (t, $^3J$ = 6.0 Hz, 2H), 2.60 - 2.53 (m, 2H), 2.02 (sm, 1H), 1.46 (t, $^3J$ = 6.9 Hz, 3H).

$^{13}C$ NMR (125 MHz, DMSO-$d_6$, 300 K): δ (ppm) = 172.7, 170.0, 169.8, 168.9, 167.2, 154.2 (d, $^1J_{C-F}$ = 243.5 Hz), 152.8, 151.8, 149.9, 147.6, 146.3, 137.9 (d, $^4J_{C-F}$ = 3.6 Hz), 136.1, 132.0, 128.2, 124.4, 123.2 (d, $^3J_{C-F}$ = 7.2 Hz), 119.5 (d, $^2J_{C-F}$ = 19.2 Hz), 117.3, 117.1, 117.0 (d, $^2J_{C-F}$ = 22.0 Hz), 113.9, 113.5, 110.6, 109.2, 108.5, 89.1, 69.8, 69.7, 69.7, 68.8, 66.5, 64.6, 48.5, 41.7, 37.0, 30.9, 22.1, 14.2.

HR-MS: calculated for $C_{40}H_{39}ClFN_7O_9H$ $(M + H)^+$: 816.2555; found: 816.2531.

Elemental analysis calculated (%) for $C_{40}H_{39}ClFN_7O_9$ x 0.50 $H_2O$: N: 11.88, C: 58.22, H: 4.89; found: N: 11.70, C: 58.05, H: 4.85.

Mp: 101.1 °C

### Scheme S4: Synthesis of LLP038[a]

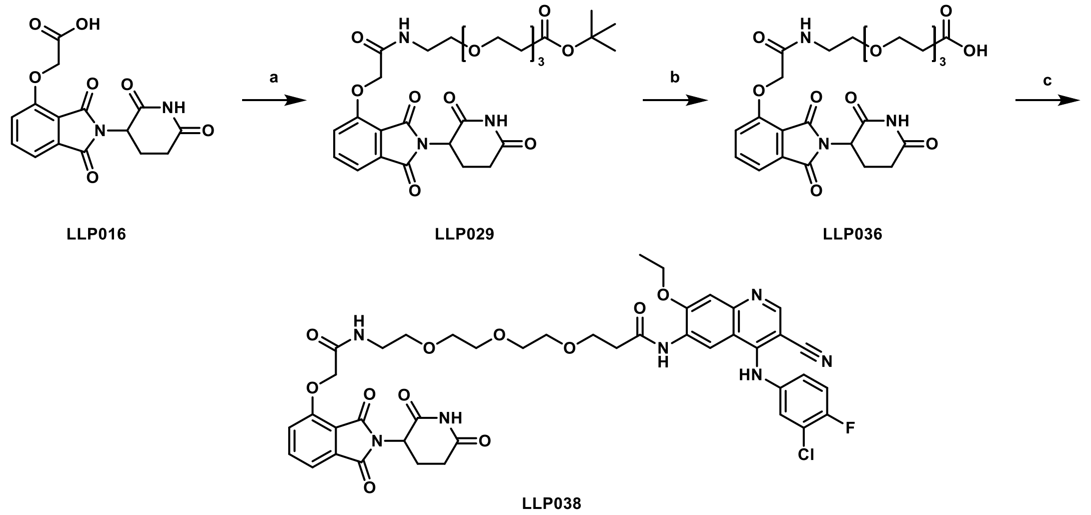

[a] Reagents and conditions: (a) *tert*-butyl 12-amino-4,7,10-trioxadodecanoate, HATU, DIPEA, DMF, r.t., overnight, 60%; (b) TFA, r.t., 3 h; (c) **DH03**, HATU, DIPEA, DMSO, r.t., overnight, 23%.

#### *tert*-Butyl 1-{[2-(2,6-dioxopiperidin-3-yl)-1,3-dioxoisoindolin-4-yl]oxy}-2-oxo-6,9,12-trioxa-3-azapentadecan-15-oate (LLP029)

**LP11** (0.60 g, 1.80 mmol, 1.00 eq) was dissolved in dry DMF (10.0 mL), mixed with *tert*-butyl 12-amino-4,7,10-trioxadodecanoate (0.49 mL, 1.80 mmol, 1.00 eq), HATU (1.03 g, 2.71 mmol, 1.50 eq) and *N*,*N*-diisopropylethylamine (0.95 mL, 5.42 mmol, 3.00 eq) and the mixture stirred under argon atmosphere overnight at r.t. Then, EtOAc (50 mL) was added and the organic phase was washed with 5% LiCl solution (aq., 30 mL). The aqueous layer was extracted with EtOAc (2 x 50 mL), the combined organic phase was again washed with 5% LiCl solution (aq., 4 x 30 mL), brine (30 mL) and finally dried over $MgSO_4$. The solvent was removed under reduced pressure, then the residue was adsorbed on silica gel and purified by column chromatography (cHex/EtOAc 50:50 → cHex/EtOAc 0:100 over 25 min; EtOAc → EtOAc/MeOH 90:10 over 15 min). The product **LLP029** was obtained as a gray solid (0.63 g, 1.07 mmol) in 60% yield.

$^1$H NMR (500 MHz, DMSO-$d_6$, 300 K): δ (ppm) = 11.09 (s, 1H), 7.98 (t, $^3J$ = 5.7 Hz, 1H7.81 (dd, $^3J$ = 8.6 Hz, $^3J$ = 7.5 Hz, 1H), 7.50 (d, $^3J$ = 7.5 Hz, 1H), 7.40 (d, $^3J$ = 8.6 Hz, 1H), 5.11 (dd, $^3J$ = 12.9 Hz, $^3J$ = 5.4 Hz, 1H), 4.78 (s, 2H), 3.57 (t, $^3J$ = 6.3 Hz, 2H), 3.51 (ps, 4H), 3.49 - 3.45 (m, 6H), 3.33 - 3.30 (m, 2H), 2.90 (ddd, $^2J$ = 17.2 Hz, $^3J$ = 13.8 Hz, $^3J$ = 5.4 Hz, 1H), 2.62 - 2.52 (m, 2H), 2.40 (t, $^3J$ = 6.3 Hz, 2H), 2.07 - 2.02 (sm, 1H), 1.39 (s, 9H).

$^{13}$C NMR (125 MHz, DMSO-$d_6$, 300 K): δ (ppm) = 172.7, 170.3, 169.8, 166.8, 166.7, 165.4, 154.9, 136.9, 133.0, 120.4, 116.8, 116.0, 79.7, 69.7, 69.6, 68.8, 67.5, 66.2, 48.8, 38.4, 35.8, 30.9, 27.7, 22.0.

HR-MS: calculated for $C_{28}H_{37}N_3O_{11}Na$ $(M + Na)^+$: 614.2320; found: 614.2297.

Purity was determined by qNMR using 1,2,4,5-tetrachloro-3-nitrobenzene as internal standard: 98.7%.

Mp: 85.6 °C

### 1-{[2-(2,6-Dioxopiperidin-3-yl)-1,3-dioxoisoindolin-4-yl]oxy}-2-oxo-6,9,12-trioxa-3-azapentadecan-15-oic acid (LLP036)

**LLP029** (0.40 g, 0.68 mmol, 1.00 eq) and TFA (1.30 mL, 16.90 mmol, 25.00 eq) were dissolved in DCM (10.0 mL) and the reaction mixture was stirred at r.t. for 3 h. After removal of the solvent under reduced pressure and drying under high vacuum, the product **LLP036** was obtained as a white resin (0.36 g, 0.68 mmol) in quantitative yield.

$^1$H NMR (500 MHz, DMSO-$d_6$, 300 K): δ (ppm) = 11.07 (bs, 1H), 7.99 (t, $^3J$ = 5.7 Hz, 1H), 7.81 (dd, $^3J$ = 8.6 Hz, $^3J$ = 7.5 Hz, 1H), 7.50 (d, $^3J$ = 7.5 Hz, 1H), 7.40 (d, $^3J$ = 8.6 Hz, 1H), 5.11 (dd, $^3J$ = 12.9 Hz, $^3J$ = 5.4 Hz, 1H), 4.79 (s, 2H), 3.59 (t, $^3J$ = 6.3 Hz, 2H), 3.51 (ps, 4H), 3.49 - 3.46 (m, 6H), 3.32 (sm, 2H), 2.90 (ddd, $^2J$ = 17.2 Hz, $^3J$ = 13.8 Hz, $^3J$ = 5.4 Hz, 1H), 2.63 - 2.53 (m, 2H), 2.42 (t, $^3J$ = 6.3 Hz, 2H), 2.06 (sm, 1H).

$^{13}$C NMR (125 MHz, DMSO-$d_6$, 300 K): δ (ppm) = 172.7, 172.6, 169.8, 166.9, 166.7, 165.4, 155.0, 136.9, 133.0, 120.4, 116.8, 116.0, 69.7, 69.6, 68.8, 67.6, 66.3, 48.8, 38.4, 34.8, 30.9, 22.0.

HR-MS: calculated for $C_{24}H_{28}N_3O_{11}$ $(M - H)^-$: 534.1729; found: 534.1725.

### *N*-{4-[(3-Chloro-4-fluorophenyl)amino]-3-cyano-7-ethoxyquinolin-6-yl}-3-(2-{2-[2-(2-{[2-(2,6-dioxopiperidin-3-yl)-1,3-dioxoisoindolin-4-yl]oxy}acetamido)ethoxy]ethoxy}ethoxy)propanamide (LLP038)

**LLP036** (0.30 g, 0.56 mmol, 1.00 eq) was dissolved in dry DMSO (6.0 mL), **DH03** (0.24 g, 0.67 mmol, 1.20 eq), HATU (0.84 g, 0.84 mmol, 1.50 eq) and *N*,*N*-diisopropylethylamine (0.30 mL, 1.68 mmol, 3.00 eq) were added and the reaction mixture was stirred under an argon atmosphere overnight at r.t. After this time, EtOAc (40 mL) was added, the mixture was washed with water (2 x 20 mL) and brine (20 mL) and dried over $MgSO_4$. The solvent was removed under reduced pressure, then the residue was adsorbed on silica gel and purified first by column chromatography (cHex/EtOAc 50:50 → cHex/EtOAc 0:100 over 25 min; EtOAc → EtOAc/MeOH 90:10 over 20 min) and then by preparative HPLC ($H_2O$/MeCN 95:5 → 5:95 over 60 min). The product **LLP038** was obtained as a beige solid (113 mg, 0.13 mmol) in 23% yield.

$^1$H NMR (500 MHz, DMSO-$d_6$, 300 K): δ (ppm) = 11.10 (s, 1H), 9.67 (s, 1H), 9.34 (s, 1H), 8.94 (s, 1H), 8.53 (s, 1H), 7.95 (t, $^3J$ = 5.7 Hz, 1H), 7.79 (dd, $^3J$ = 8.6 Hz, $^3J$ = 7.2 Hz, 1H), 7.47 (d, $^3J$ = 7.2 Hz, 1H), 7.42 (dd, $^4J_{H-F}$ = 6.6 Hz, $^4J$ = 2.6 Hz, 1H), 7.40 (s, 1H, 8-H), 7.40 (dd, $^3J_{H-F}$ = 8.9 Hz, $^3J$ = 8.9 Hz, 1H), 7.38 (d, $^3J$ = 8.6 Hz, 1H), 7.22 (ddd, $^3J$ = 8.9 Hz, $^4J_{H-F}$ = 4.3 Hz, $^4J$ = 2.6 Hz, 1H), 5.10 (dd, $^3J$ = 12.9 Hz, $^3J$ = 5.4 Hz, 1H),

4.77 (s, 2H), 4.31 (q, $^{3}J$ = 7.2 Hz, 2H), 3.73 (t, $^{3}J$ = 6.0 Hz, 2H), 3.56 - 3.48 (m, 8H), 3.46 (t, $^{3}J$ = 5.4 Hz, 2H), 3.30 - 3.27 (m, 2H), 2.89 (ddd, $^{2}J$ = 17.2 Hz, $^{3}J$ = 13.8 Hz, $^{3}J$ = 5.4 Hz, 1H), 2.72 (t, $^{3}J$ = 6.0 Hz, 2H), 2.62 - 2.52 (m, 2H), 2.04 (sm, 1H), 1.46 (t, $^{3}J$ = 7.2 Hz, 3H).

$^{13}$C NMR (125 MHz, DMSO-$d_6$, 300 K): δ (ppm) = 172.7, 169.8, 166.8, 166.6, 165.4, 154.9, 154.2 (d, $^{1}J_{C\text{-}F}$ = 243.5 Hz), 152.8, 151.8, 149.9, 147.6, 137.9 (d, $^{4}J_{C\text{-}F}$ = 3.6 Hz), 136.8, 133.0, 128.2, 124.4, 123.2 (d, $^{3}J_{C\text{-}F}$ = 7.2 Hz), 120.3, 119.5 (d, $^{2}J_{C\text{-}F}$ = 18.0 Hz), 117.1, 117.0 (d, $^{2}J_{C\text{-}F}$ = 21.6 Hz), 116.7, 116.0, 113.9, 113.5, 108.5, 89.1, 69.7, 69.7, 69.6, 69.6, 68.7, 67.5, 66.5, 64.6, 48.8, 38.3, 37.0, 30.9, 22.0, 14.2.

HR-MS: calculated for $C_{42}H_{41}ClFN_7O_{11}H$ (M + H)$^+$: 874.2609; found: 874.2590.

Elemental analysis calculated (%) for $C_{42}H_{41}ClFN_7O_{11}$ x 1.00 $H_2O$: N: 10.99, C: 56.54, H: 4.86; found: N: 10.96, C: 56.61, H: 4.85.

Mp: 145.1 °C

Scheme S5: Synthesis of LLP031[a]

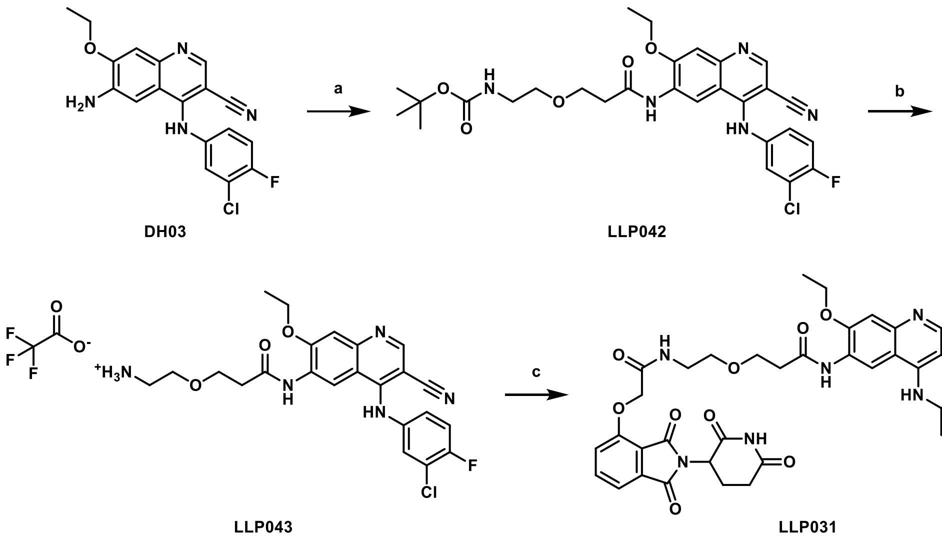

[a] Reagents and conditions: (a) 3-[2-(*tert*-butoxycarbonylamino)ethoxy]propionic acid, HATU, DIPEA, DMSO, r.t., overnight, 63%; (b) TFA, DCM, r.t., 2 h, 100%; (c) LLP016, HATU, DIPEA, DMF, r.t., overnight, 36%.

### *tert*-Butyl {2-[3-({4-[(3-chloro-4-fluorophenyl)amino]-3-cyano-7-ethoxyquinolin-6-yl}amino)-3-oxopropoxy]ethyl}carbamate (LLP042)

**DH03** (0.79 g, 2.21 mmol, 1.00 eq) was dissolved in dry DMSO (23.0 mL), mixed with 3-[2-(*tert*-butoxycarbonylamino)ethoxy]propionic acid (0.57 g, 2.46 mmol, 1.10 eq), HATU (1.34 g, 3.51 mmol, 1.50 eq) and *N*,*N*-diisopropylethylamine (1.23 mL, 7.02 mmol, 3.00 eq) and the reaction mixture stirred under an argon atmosphere overnight at r.t. Then, EtOAc (100 mL) was added, the reaction mixture was washed with water (5 x 30 mL), brine (30 mL) and the organic phase was dried over $MgSO_4$. The solvent was removed under reduced pressure, then the residue was adsorbed on silica gel and purified by column chromatography (cHex/EtOAc 30:70 → cHex/EtOAc 0:100 over 40 min). The product **LLP042** was obtained as a yellow solid (0.80 g, 1.40 mmol) in 63% yield.

$^1$H NMR (500 MHz, DMSO-*d$_6$*, 300 K): δ (ppm) = 9.68 (s, 1H), 9.35 (s, 1H), 8.94 (s, 1H), 8.54 (s, 1H), 7.43 (dd, $^4J_{H\text{-}F}$ = 6.6 Hz, $^4J$ = 2.9 Hz, 1H), 7.41 (s, 1H), 7.40 (dd, $^3J_{H\text{-}F}$ = 9.2 Hz, $^3J$ = 8.9 Hz, 1H), 7.22 (ddd, $^3J$ = 8.9 Hz, $^4J_{H\text{-}F}$ = 4.3 Hz, $^4J$ = 2.9 Hz, 1H), 6.73 (t, $^3J$ = 6.0 Hz, 1H), 4.32 (q, $^3J$ = 6.9 Hz, 2H), 3.72 (t, $^3J$ = 6.0 Hz, 2H), 3.43 (t, $^3J$ = 6.3 Hz, 2H), 3.10 (sm, 2H), 2.72 (t, $^3J$ = 6.3 Hz, 2H), 1.47 (t, $^3J$ = 6.9 Hz, 3H), 1.35 (s, 9H).

$^{13}$C NMR (125 MHz, DMSO-*d$_6$*, 300 K): δ (ppm) = 169.7, 155.5, 154.2 (d, $^1J_{C\text{-}F}$ = 243.5 Hz), 152.9, 151.8, 150.0, 147.6, 137.9 (d, $^4J_{C\text{-}F}$ = 3.6 Hz), 128.2, 124.4, 123.2 (d, $^3J_{C\text{-}F}$ = 7.2 Hz), 119.5 (d, $^2J_{C\text{-}F}$ = 19.2 Hz), 117.1, 117.0 (d, $^2J_{C\text{-}F}$ = 21.6 Hz), 113.9, 113.6, 108.6, 89.1, 77.6, 69.0, 66.1, 64.6, 39.5, 36.9, 28.1, 14.2.

HR-MS: calculated for $C_{28}H_{31}ClFN_5O_5Na$ (M + Na)$^+$: 594.1890; found: 594.1871.

Purity was determined by qNMR using dimethyl terephthalate as internal standard: 95.3%.

Mp: 87.1 °C

### 2-[3-({4-[(3-Chloro-4-fluorophenyl)amino]-3-cyano-7-ethoxyquinolin-6-yl}amino)-3-oxopropoxy]ethan-1-aminium 2,2,2-trifluoroacetate (LLP043)

**LLP042** (0.40 g, 0.70 mmol, 1.00 eq) was dissolved in DCM (10.0 mL), TFA (1.08 mL, 13.99 mmol, 20.00 eq) was added and the mixture was stirred for 2 h at r.t. **LLP043** (0.41 g, 0.70 mmol) was obtained as a pale-yellow solid in quantitative yield after removal of the solvent under reduced pressure and drying under high vacuum.

$^{1}$H NMR (500 MHz, DMSO-$d_6$, 300 K): δ (ppm) = 9.73 (bs, 1H), 9.41 (s, 1H), 8.90 (s, 1H), 8.56 (s, 1H), 7.87 (bs, 3H), 7.45 - 7.43 (m, 1H), 7.42 (s, 1H), 7.41 (dd, $^{3}J_{H\text{-}F}$ = 9.2 Hz, $^{3}J$ = 8.9 Hz, 1H), 7.24 - 7.21 (m, 1H), 4.32 (q, $^{3}J$ = 6.9 Hz, 2H), 3.77 (t, $^{3}J$ = 6.0 Hz, 2H), 3.63 (t, $^{3}J$ = 5.4 Hz, 2H), 3.00 (ps, 2H), 2.78 (t, $^{3}J$ = 6.3 Hz, 2H), 1.47 (t, $^{3}J$ = 6.9 Hz, 3H).

$^{13}$C NMR (125 MHz, DMSO-$d_6$, 300 K): δ (ppm) = 169.6, 158.0 (d, $^{2}J_{C\text{-}F}$ = 31.2 Hz), 154.3 (d, $^{1}J_{C\text{-}F}$ = 242.3 Hz), 153.1, 151.7, 150.1, 147.4, 137.9 (d, $^{4}J_{C\text{-}F}$ = 3.6 Hz), 128.0, 124.6, 123.4 (d, $^{3}J_{C\text{-}F}$ = 7.2 Hz), 119.5 (d, $^{2}J_{C\text{-}F}$ = 19.2 Hz), 117.2 (d, $^{1}J_{C\text{-}F}$ = 299.9 Hz), 117.0 (d, $^{2}J_{C\text{-}F}$ = 21.6 Hz), 117.0, 114.4, 113.9, 108.4, 89.0, 66.3, 66.3, 64.6, 38.4, 36.5, 14.2.

HR-MS: calculated for $C_{23}H_{24}ClFN_5O_3$ $(M)^+$: 472.1546; found: 472.1538.

Mp: 117.5 °C

### *N*-{4-[(3-Chloro-4-fluorophenyl)amino]-3-cyano-7-ethoxyquinolin-6-yl}-3-[2-(2-{[2-(2,6-dioxopiperidin-3-yl)-1,3-dioxoisoindolin-4-yl]oxy}acetamido)ethoxy]propenamide (LLP031)

**LLP043** (130 mg, 0.26 mmol, 1.00 eq) was dissolved in dry DMF (3.0 mL), mixed with **LLP016** (93 mg, 0.28 mmol, 1.10 eq), HATU (148 mg, 0.39 mmol, 1.50 eq) and *N*,*N*-diisopropylethylamine (0.14 mL, 0.78 mmol, 3.00 eq) and the reaction mixture was stirred under an argon atmosphere overnight at r.t. Then, EtOAc (40 mL) was added and the mixture was washed with 5% LiCl solution (aq., 20 mL). The aqueous layer was extracted with EtOAc (2 x 20 mL). The combined organic phase was washed with 5% LiCl solution (aq., 4 x 20 mL), brine (10 mL) and finally dried over $MgSO_4$. The solvent was removed under reduced pressure, then the residue was adsorbed on silica gel and purified by column chromatography (cHex/EtOAc 50:50 → cHex/EtOAc 0:100; EtOAc/MeOH 100:0 over 25 min; EtOAc → EtOAc/MeOH 90:10 over 20 min). The product **LLP031** was obtained as a beige solid (75 mg, 0.09 mmol) in 36% yield.

$^{1}$H NMR (500 MHz, DMSO-$d_6$, 300 K): δ (ppm) = 11.10 (s, 1H), 9.71 (bs, 1H), 9.33 (s, 1H), 8.91 (s, 1H), 8.54 (s, 1H), 7.96 (t, $^{3}J$ = 5.7 Hz, 1H), 7.72 (dd, $^{3}J$ = 8.6 Hz, $^{3}J$ = 7.2 Hz, 1H), 7.42 (dd, $^{4}J_{H\text{-}F}$ = 6.6 Hz, $^{4}J$ = 2.6 Hz, 1H), 7.40 (d, $^{3}J$ = 7.2 Hz, 1H), 7.40 (dd, $^{3}J_{H\text{-}F}$ = 9.2 Hz, $^{3}J$ = 8.9 Hz, 1H), 7.35 (s, 1H), 7.32 (d, $^{3}J$ =

8.6 Hz, 1H), 7.22 (ddd, $^3J$ = 8.9 Hz, $^4J_{H-F}$ = 4.3 Hz, $^4J$ = 2.6 Hz, 1H), 5.09 (dd, $^3J$ = 5.4 Hz, $^3J$ = 12.9 Hz, 1H), 4.73 (s, 2H), 4.29 (q, $^3J$ = 6.9 Hz, 2H), 3.76 (t, $^3J$ = 6.0 Hz, 2H), 3.54 (t, $^3J$ = 5.7 Hz, 2H), 3.37 (sm, 2H), 2.89 (ddd, $^2J$ = 17.2 Hz, $^3J$ = 13.8 Hz, $^3J$ = 5.4 Hz, 1H), 2.73 (t, $^3J$ = 6.0 Hz, 2H), 2.60 - 2.52 (m, 2H), 2.05 (sm, 1H), 1.45 (t, $^3J$ = 6.9 Hz, 3H).

$^{13}$C NMR (125 MHz, DMSO-$d_6$, 300 K): δ (ppm) = 172.7, 169.8, 169.7, 166.9, 166.6, 165.4, 154.7, 154.4 (d, $^1J_{C-F}$ = 243.5 Hz), 152.8, 151.5, 150.1, 147.2, 137.9 (d, $^4J_{C-F}$ = 3.6 Hz), 136.7, 132.8, 128.2, 124.8, 123.6 (d, $^3J_{C-F}$ = 7.2 Hz), 120.2, 119.5 (d, $^2J_{C-F}$ = 19.2 Hz), 117.0 (d, $^2J_{C-F}$ = 21.6 Hz), 116.9, 116.6, 115.9, 113.7, 113.4, 108.1, 88.7, 69.6, 69.5, 68.5, 67.5, 66.2, 64.7, 48.8, 38.2, 36.9, 30.9, 21.9, 14.2.

HR-MS: calculated for $C_{38}H_{33}ClFN_7O_9H$ (M + H)$^+$: 786.2085; found: 786.2062.

Elemental analysis calculated (%) for $C_{38}H_{33}ClN_7O_9$ x 0.50 $H_2O$: N: 12.33, C: 57.40, H: 4.31; found: N: 12.17, C: 57.13, H: 4.42.

Mp: 226.8 °C

Scheme S6: Synthesis of LLP049[a]

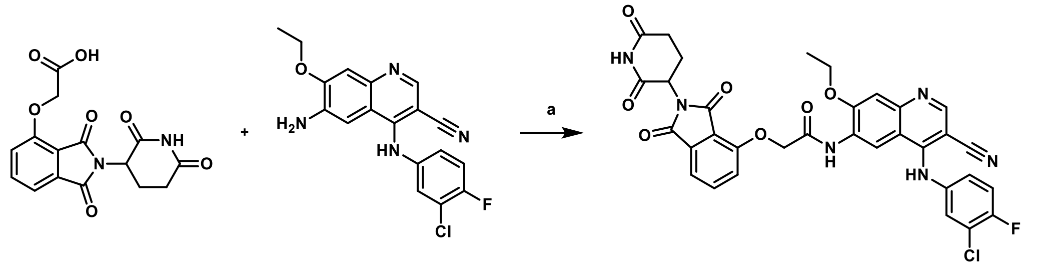

[a] Reagents and conditions: (a) HATU, DIPEA, DMF, r.t., overnight, 20%.

### *N*-{4-[(3-Chloro-4-fluorophenyl)amino]-3-cyano-7-ethoxyquinolin-6-yl}-2-{[2-(2,6-dioxopiperidin-3-yl)-1,3-dioxoisoindolin-4-yl]oxy}acetamide (LLP049)

**DH03** (0.40 g, 1.12 mmol, 1.00 eq) was dissolved in dry DMF (11.0 mL), **LLP016** (0.37 g, 1.12 mmol, 1.00 eq), HATU (0.64 g, 1.68 mmol, 1.50 eq) and *N*,*N*-diisopropylethylamine (0.59 mL, 3.36 mmol, 3.00 eq) were added and the reaction mixture was stirred under an argon atmosphere overnight at r.t. Subsequently, EtOAc (50 mL) was added and the mixture was washed with 5% LiCl solution (aq., 25 mL). A whitish precipitate formed in the organic phase, which was filtered off. The filter cake was washed with EtOAc (2 x 10 mL), $H_2O$ (2 x 10 mL) and DEE (2 x 10 mL). The product **LLP049** was obtained as a beige solid (150 mg, 0.22 mmol) in 20% yield.

$^1$H NMR (500 MHz, DMSO-$d_6$, 300 K): δ (ppm) = 11.13 (s, 1H), 9.79 (bs, 1H), 9.56 (s, 1H), 9.07 (s, 1H), 8.57 (s, 1H), 7.86 (dd, $^3J$ = 8.6 Hz, $^3J$ = 7.2 Hz, 1H), 7.58 (d, $^3J$ = 8.6 Hz, 1H), 7.54 (d, $^3J$ = 7.2 Hz, 1H), 7.35 (s, 1H), 7.44 (dd, $^4J_{H\text{-}F}$ = 6.6 Hz, $^4J$ = 2.6 Hz, 1H), 7.40 (dd, $^3J_{H\text{-}F}$ = 9.2 Hz, $^3J$ = 8.9 Hz, 1H), 7.24 - 7.21 (m, 1H), 5.14 (dd, $^3J$ = 12.9 Hz, $^3J$ = 5.4 Hz, 1H), 5.12 (s, 2H), 4.38 (q, $^3J$ = 6.9 Hz, 2H), 2.92 (ddd, $^2J$ = 17.2 Hz, $^3J$ = 13.8 Hz, $^3J$ = 5.4 Hz, 1H), 2.64 - 2.54 (m, 2H), 2.10 - 2.05 (sm, 1H), 1.43 (t, $^3J$ = 6.9 Hz, 3H).

$^{13}$C NMR (125 MHz, DMSO-$d_6$, 300 K): δ (ppm) = 172.7, 169.8, 166.7, 165.8, 165.3, 154.3 (d, $^1J_{C\text{-}F}$ = 243.5 Hz), 154.2, 152.2, 151.9, 150.2, 147.5, 137.9 (d, $^4J_{C\text{-}F}$ = 3.6 Hz), 137.0, 133.1, 127.2, 124.6, 123.4 (d, $^3J_{C\text{-}F}$ = 7.2 Hz), 120.2, 119.5 (d, $^2J_{C\text{-}F}$ = 19.2 Hz), 117.0 (d, $^2J_{C\text{-}F}$ = 22.8 Hz), 116.9, 116.6, 116.3, 113.9, 112.9, 108.4, 89.2, 67.4, 64.9, 48.8, 30.9, 22.0, 13.9.

HR-MS: calculated for $C_{33}H_{24}ClFN_6O_7H$ $(M + H)^+$: 671.1452; found: 671.1430.

Purity was determined by qNMR using dimethyl terephthalate as internal standard: 96.1%.

Mp: 218.2 °C

Scheme S7: Synthesis of LLP041[a]

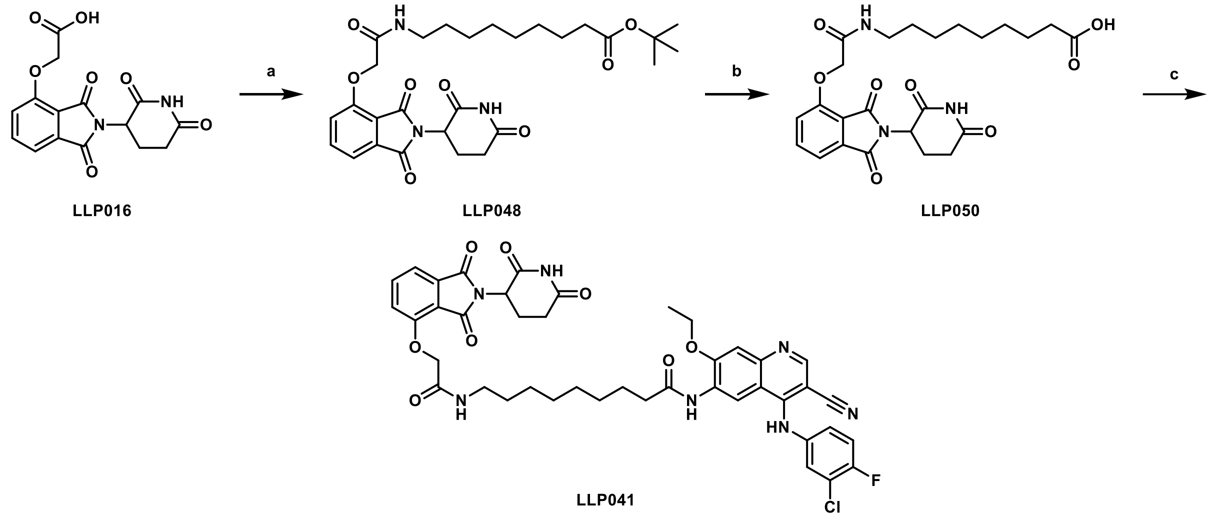

[a] Reagents and conditions: (a) *tert*-butyl 9-aminononanoate, HATU, DIPEA, DMF, r.t., 72 h, 50%; (b) TFA, r.t., 3 h; (c) **DH03**, HATU, DIPEA, DMSO, r.t., overnight, 46%.

### *tert*-Butyl 9-(2-{[2-(2,6-dioxopiperidin-3-yl)-1,3-dioxoisoindolin-4-yl]oxy}acetamido)nonanoate (LLP048)

**LLP016** (0.60 g, 2.18 mmol, 1.00 eq) was dissolved in dry DMF (15.0 mL), mixed with *tert*-butyl 9-aminononanoate (0.50 g, 2.18 mmol, 1.00 eq), HATU (1.24 g, 3.27 mmol, 1.50 eq) and *N*,*N*-diisopropylethylamine (1.14 mL, 6.54 mmol, 3.00 eq) and the reaction mixture stirred under an argon atmosphere for 72 h at r.t. Then, EtOAc (50 mL) was added and the reaction mixture was washed with 5% LiCl solution (aq., 5 x 30 mL), brine (20 mL) and dried over $MgSO_4$. The solvent was removed under reduced pressure, then the residue was adsorbed on silica gel and purified by column chromatography (cHex/EtOAc 80:20 → cHex/EtOAc 0:100 over 30 min). The product **LLP048** was obtained as a gray solid (0.59 g, 1.09 mmol) in 50% yield.

$^1$H NMR (500 MHz, DMSO-*d*$_6$, 300 K): δ (ppm) = 11.09 (bs, 1H), 7.90 (t, $^3J$ = 5.7 Hz, 1H), 7.81 (dd, $^3J$ = 8.6 Hz, $^3J$ = 7.2 Hz, 1H), 7.50 (dd, $^3J$ = 7.2 Hz, 1H), 7.40 (dd, $^3J$ = 8.6 Hz, 1H), 5.12 (dd, $^3J$ = 12.9 Hz, $^3J$ = 5.4 Hz, 1H), 4.76 (s, 2H), 3.14 (dt, $^3J$ = 7.2 Hz, $^3J$ = 5.7 Hz, 2H), 2.90 (ddd, $^2J$ = 17.2 Hz, $^3J$ = 13.8 Hz, $^3J$ = 5.4 Hz, 1H), 2.62 - 2.52 (m, 2H), 2.15 (t, $^3J$ = 7.5 Hz, 2H), 2.07 - 2.02 (m, 1H), 1.48 - 1.42 (m, 4H), 1.39 (s, 9H), 1.24 (ps, 8H).

$^{13}$C NMR (125 MHz, DMSO-*d*$_6$, 300 K): δ (ppm) = 172.6, 172.2, 169.8, 166.7, 166.5, 165.5, 155.0, 136.8, 133.0, 120.4, 116.8, 116.0, 79.3, 67.7, 48.8, 38.3, 34.7, 30.9, 28.9, 28.6, 28.5, 28.3, 27.7, 26.2, 24.5, 22.0.

HR-MS: calculated for $C_{28}H_{37}N_3O_8Na$ $(M + Na)^+$: 566.2473; found: 566.2457.

Purity was determined by qNMR using 1,2,4,5-tetrachloro-3-nitrobenzene as internal standard: 98.6%.

Mp: 99.9 °C

#### 9-(2-{[2-(2,6-Dioxopiperidin-3-yl)-1,3-dioxoisoindolin-4-yl]oxy}acetamido)nonanoic acid (LLP050)

**LLP048** (0.45 g, 0.83 mmol, 1.00 eq) and TFA (1.59 mL, 20.69 mmol, 25.00 eq) were dissolved in DCM (10.0 mL) and the reaction mixture was stirred for 3 h at r.t. Removal of the solvent under reduced pressure followed by drying under high vacuum yielded **LLP050** as an off-white solid (0.41 g, 0.83 mmol) in quantitative yield.

$^1$H NMR (500 MHz, DMSO-$d_6$, 300 K): δ (ppm) = 11.93 (bs, 1H), 11.09 (s, 1H), 7.90 (t, $^3J$ = 5.7 Hz, 1H), 7.81 (dd, $^3J$ = 8.6 Hz, $^3J$ = 7.2 Hz, 1H), 7.50 (dd, $^3J$ = 7.2 Hz, 1H), 7.40 (dd, $^3J$ = 8.6 Hz, 1H), 5.12 (dd, $^3J$ = 12.9 Hz, $^3J$ = 5.4 Hz, 1H), 4.76 (s, 2H), 3.14 (sm, 2H), 2.90 (ddd, $^2J$ = 17.2 Hz, $^3J$ = 13.8 Hz, $^3J$ = 5.4 Hz, 1H), 2.62 - 2.52 (m, 2H), 2.18 (t, $^3J$ = 7.5 Hz, 2H), 2.07 - 2.02 (m, 1H), 1.49 - 1.42 (m, 4H), 1.24 (ps, 8H).

$^{13}$C NMR (125 MHz, DMSO-$d_6$, 300 K): δ (ppm) = 174.4, 172.7, 169.8, 166.7, 166.5, 165.5, 155.0, 136.9, 133.0, 120.4, 116.8, 116.0, 67.7, 48.8, 38.3, 33.6, 30.9, 28.9, 28.6, 28.5, 28.4, 26.2, 24.4, 22.0.

HR-MS: calculated for $C_{24}H_{28}N_3O_8$ $(M - H)^-$: 486.1882; found: 486.1879.

Mp: 135.7 °C

#### *N*-{4-[(3-Chloro-4-fluorophenyl)amino]-3-cyano-7-ethoxyquinolin-6-yl}-9-(2-{[2-(2,6-dioxopiperidin-3-yl)-1,3-dioxoisoindolin-4-yl]oxy}acetamido)nonanamide (LLP041)

**LLP050** (0.30 g, 0.62 mmol, 1.00 eq) was dissolved in dry DMSO (8.0 mL), **DH03** (0.24 g, 0.68 mmol, 1.10 eq), HATU (0.35 g, 0.92 mmol, 1.50 eq) and *N*,*N*-diisopropylethylamine (0.32 mL, 1.85 mmol, 3.00 eq) were added and the resulting mixture was stirred under an argon atmosphere overnight at r.t. After addition of EtOAc (50 mL), the mixture was washed with $H_2O$ (5 x 20 mL), brine (15 mL) and dried over $MgSO_4$. The solvent was removed under reduced pressure, then the residue was adsorbed on silica gel and purified by column chromatography (cHex/EtOAc 70:30 → cHex/EtOAc 0:100 over 20 min; EtOAc → EtOAc/MeOH 90:10 over 13 min). The product **LLP041** was obtained as a pale-yellow solid (0.24 g, 0.29 mmol) in 46% yield.

$^1$H NMR (400 MHz, DMSO-$d_6$, 300 K): δ (ppm) = 11.10 (s, 1H), 9.66 (s, 1H), 9.22 (s, 1H), 8.84 (s, 1H), 8.53 (s, 1H), 7.90 (s, $^3J$ = 5.7 Hz, 1H), 7.80 (dd, $^3J$ = 8.5 Hz, $^3J$ = 7.3 Hz, 1H), 7.48 (d, $^3J$ = 7.3 Hz, 1H), 7.44 - 7-42 (m, 1H), 7.42 - 7.38 (m, 1H), 7.40 (s, 1H), 7.39 (d, $^3J$ = 8.5 Hz, 1H), 7.24 - 7.21 (m, 1H), 5.11 (dd, $^3J$ = 12.8 Hz, $^3J$ = 5.3 Hz, 1H), 4.76 (s, 2H), 4.30 (q, 2H, $^3J$ = 6.9 Hz), 3.14 (sm, 2H),2.90 (ddd, $^2J$ = 17.2 Hz, $^3J$ =

13.7 Hz, $^{3}J$ = 5.3 Hz, 1H), 2.63 - 2.52 (m, 2H), 2.45 (t, $^{3}J$ = 7.3 Hz, 2H), 2.08 - 2.01 (m, 1H), 1.64 - 1.57 (m, 2H), 1.46 (t, $^{3}J$ = 6.9 Hz, 2H), 1.48 - 1.42 (m, 2H), 1.34 - 1.24 (m, 8H).

$^{13}$C NMR (100 MHz, DMSO-$d_6$, 300 K): δ (ppm) = 172.7, 171.7, 169.8, 166.7, 166.6, 165.5, 155.0, 154.3 (d, $^{1}J_{C-F}$ = 243.7 Hz), 153.4, 151.8, 150.0, 147.7, 137.8, 136.9, 133.0, 128.2, 124.7, 123.4 (d, $^{3}J_{C-F}$ = 5.8 Hz), 120.4, 119.5 (d, $^{2}J_{C-F}$ = 19.3 Hz), 117.1, 117.0 (d, $^{2}J_{C-F}$ = 21.2 Hz), 116.8, 116.0, 114.5, 113.8, 108.6, 88.8, 67.7, 64.5, 48.8, 38.3, 36.0, 30.9, 28.9, 28.8, 28.6, 28.5, 26.2, 25.1, 22.0, 14.2.

HR-MS: calculated for $C_{42}H_{41}ClFN_7O_8H$ $(M + H)^+$: 826.2762; found: 826.2747.

Purity was determined by qNMR using maleic acid as internal standard: 91.9%.

Mp: 182.2 °C

Scheme S8: Synthesis of LP15[a]

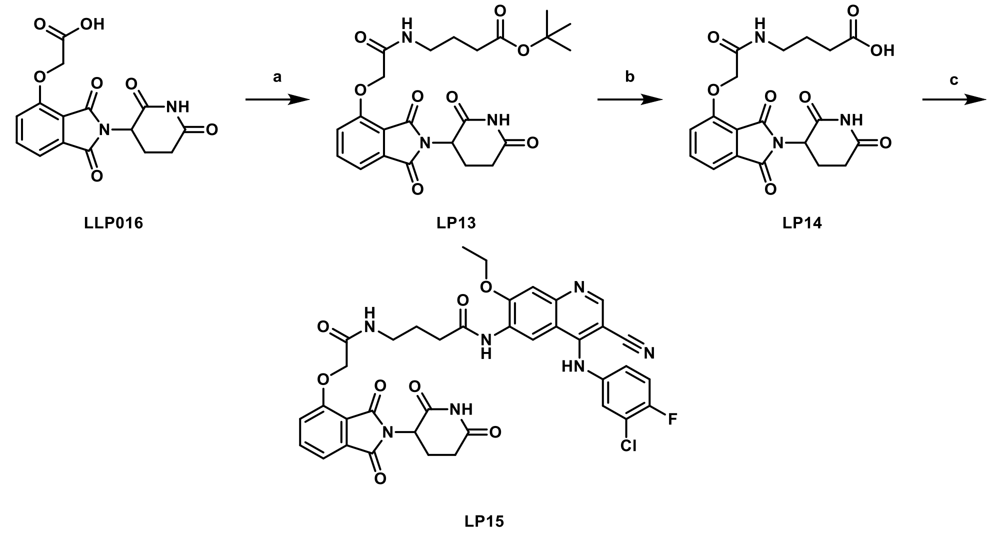

[a] Reagents and conditions: (a) *tert*-butyl 4-aminobutanoate hydrochloride, HATU, DIPEA, DMF, r.t., overnight, 62%; (b) TFA, DCM, r.t., overnight, 100%; (c) **DH03**, HATU, TEA, DMSO, r.t., 72 h, 19%.

### *tert*-Butyl 4-(2-{[2-(2,6-dioxopiperidin-3-yl)-1,3-dioxoisoindolin-4-yl]oxy}acetamido)butanoate (LP13)

**LLP016** (1.00 g, 3.01 mmol, 1.00 eq) was dissolved in dry DMF (17.1 mL), mixed with *tert*-butyl 4-aminobutanoate hydrochloride (0.77 g, 3.31 mmol, 1.10 eq), HATU (1.72 g, 4.52 mmol, 1.50 eq) and *N*,*N*-diisopropylethylamine (1.58 mL, 9.03 mmol, 3.00 eq) and the reaction mixture was stirred under an argon atmosphere overnight at r.t. After addition of EtOAc (70 mL), the mixture was washed with 5% LiCl solution (aq., 30 mL). The aqueous phase was extracted with EtOAc (3 x 50 mL). The combined organic phase was washed with 5% LiCl solution (aq., 5 x 40 mL), brine (30 mL) and finally dried over $MgSO_4$. The solvent was removed under reduced pressure, then the residue was adsorbed on silica gel and purified by column chromatography (cHex/EtOAc 70:30 → cHex/EtOAc 0:100 over 50 min). The product **LP13** was obtained as a yellowish solid (0.88 g, 1.85 mmol) in 62% yield.

$^1$H NMR (500 MHz, DMSO-$d_6$, 300 K): δ (ppm) = 11.09 (s, 1H), 7.97 (t, $^3J$ = 5.7 Hz, 1H), 7.81 (dd, $^3J$ = 8.6 Hz, $^3J$ = 7.5 Hz, 1H), 7.50 (d, $^3J$ = 7.5 Hz, 1H), 7.40 (d, $^3J$ = 8.6 Hz, 1H), 5.12 (dd, $^3J$ = 12.9 Hz, $^3J$ = 5.4 Hz, 1H), 4.77 (s, 2H), 3.16 (sm, 2H), 2.90 (ddd, $^2J$ = 17.2 Hz, $^3J$ = 13.8 Hz, $^3J$ = 5.4 Hz, 1H), 2.63 - 2.52 (m, 2H), 2.21 (t, $^3J$ = 7.5 Hz, 2H), 2.07 - 2.02 (m, 1H), 1.65 (sm, 2H), 1.39 (s, 9H).

$^{13}$C NMR (125 MHz, DMSO-$d_6$, 300 K): δ (ppm) = 172.7, 171.8, 169.8, 166.8, 166.7, 165.4, 155.1, 136.8, 133.0, 120.4, 116.8, 116.0, 79.5, 67.7, 48.8, 37.7, 32.1, 30.9, 27.7, 24.5, 22.0.

HR-MS: calculated for $C_{23}H_{26}N_3O_8$ $(M - H)^-$: 472.1725; found: 472.1718.

Purity was determined by qNMR using maleic acid as internal standard: 98.2%.

Mp: 141.8 °C

#### 4-(2-{[2-(2,6-Dioxopiperidin-3-yl)-1,3-dioxoisoindolin-4-yl]oxy}acetamido)butanoic acid (LP14)

**LP13** (0.70 g, 1.48 mmol, 1.00 eq) and TFA (1.38 mL, 17.97 mmol, 12.14 eq) were dissolved in DCM (7.0 mL) and the mixture was stirred overnight at r.t. **LP14** was obtained as an off-white solid (0.62 g, 1.48 mmol) in quantitative yield after removal of the solvent under reduced pressure and drying under high vacuum.

$^{1}$H NMR (500 MHz, DMSO-*d*$_6$, 300 K): δ (ppm) = 11.92 (bs, 1H), 11.09 (s, 1H), 7.98 (t, $^{3}J$ = 5.7 Hz, 1H), 7.81 (dd, $^{3}J$ = 8.6 Hz, $^{3}J$ = 7.5 Hz, 1H), 7.49 (d, $^{3}J$ = 7.5 Hz, 1H), 7.39 (d, $^{3}J$ = 8.6 Hz, 1H), 5.12 (dd, $^{3}J$ = 12.9 Hz, $^{3}J$ = 5.4 Hz, 1H), 4.77 (s, 2H), 3.16 (sm, 2H), 2.90 (ddd, $^{2}J$ = 17.2 Hz, $^{3}J$ = 13.8 Hz, $^{3}J$ = 5.4 Hz, 1H), 2.63 - 2.52 (m, 2H), 2.23 (t, $^{3}J$ = 7.5 Hz, 2H), 2.06 - 2.02 (m, 1H), 1.66 (sm, 2H).

$^{13}$C NMR (125 MHz, DMSO-*d*$_6$, 300 K): δ (ppm) = 174.1, 172.7, 169.8, 166.8, 166.7, 165.4, 155.1, 136.9, 133.0, 120.4, 116.8, 116.0, 67.7, 48.8, 37.8, 30.9, 24.5, 22.0.

HR-MS: calculated for $C_{19}H_{18}N_3O_8$ (M - H)$^-$: 416.1099; found: 416.1095.

Mp: 206.5 °C

#### *N*-{4-[(3-Chloro-4-fluorophenyl)amino]-3-cyano-7-ethoxyquinolin-6-yl}-4-(2-{[2-(2,6-dioxopiperidin-3-yl)-1,3-dioxoisoindolin-4-yl]oxy}acetamido)butanamide (LP15)

**LP14** (0.50 g, 1.20 mmol, 1.00 eq) was dissolved in dry DMSO (31.2 mL), **DH03** (0.47 g, 1.32 mmol, 1.10 eq), HATU (0.68 g, 1.80 mmol, 1.50 eq) and triethylamine (0.67 mL, 4.80 mmol, 4.00 eq) were added and the resulting mixture was stirred under an argon atmosphere for 72 h at r.t. Then, EtOAc (60 mL) was added and the mixture was washed with $H_2O$ (5 x 30 mL), brine (20 mL) and finally dried over $MgSO_4$. The solvent was removed under reduced pressure, then the residue was adsorbed on silica gel and purified by column chromatography (cHex/EtOAc 50:50 → cHex/EtOAc 0:100 over 25 min; EtOAc → EtOAc/MeOH 90:10 over 15 min). The product **LP15** was obtained as a yellow solid (168 mg, 0.11 mmol) in 19% yield.

$^{1}$H NMR (400 MHz, DMSO-*d*$_6$, 300 K): δ (ppm) = 11.09 (s, 1H), 9.71 (bs, 1H), 9.31 (s, 1H), 8.86 (s, 1H), 8.54 (s, 1H), 8.02 (t, $^{3}J$ = 5.7 Hz, 1H), 7.80 (dd, $^{3}J$ = 8.5 Hz, $^{3}J$ = 7.3 Hz, 1H), 7.47 (d, $^{3}J$ = 7.3 Hz, 1H), 7.44 (dd, $^{4}J_{H-F}$ = 6.6 Hz, $^{4}J$ = 2.5 Hz, 1H), 7.41 (dd, $^{3}J_{H-F}$ = 9.2 Hz, $^{3}J$ = 8.2 Hz, 1H), 7.40 (d, $^{3}J$ = 8.5 Hz, 1H), 7.39 (s, 1H), 7.23 (ddd, $^{3}J$ = 8.9 Hz, $^{4}J_{H-F}$ = 3.9 Hz, $^{4}J$ = 2.7 Hz, 1H), 5.11 (dd, $^{3}J$ = 12.8 Hz, $^{3}J$ = 5.5 Hz, 1H), 4.79 (s, 2H), 4.30 (q, 2H, $^{3}J$ = 6.9 Hz), 4.32 - 4.27 (m, 2H), 2.88 (ddd, $^{2}J$ = 17.2 Hz, $^{3}J$ = 13.7 Hz, $^{3}J$ = 5.5 Hz, 1H), 2.61 - 2.48 (m, 4H), 2.07 - 2.00 (m, 1H), 1.80 (sm, 2H), 1.45 (t, $^{3}J$ = 6.9 Hz, 3H).

$^{13}$C NMR (100 MHz, DMSO-*d*$_6$, 300 K): δ (ppm) = 172.7, 171.3, 169.8, 166.9, 166.7, 165.4, 155.0, 154.4 (d, $^{1}J_{C-F}$ = 243.7 Hz), 153.4, 151.7, 150.1, 147.4, 137.7, 136.9, 133.0, 128.2, 124.8, 123.6 (d, $^{3}J_{C-F}$ = 6.7 Hz),

120.4, 119.5 (d, $^2J_{C-F}$ = 19.3 Hz), 117.0, 117.0 (d, $^2J_{C-F}$ = 22.2 Hz), 116.8, 116.0, 114.5, 113.8, 108.4, 88.7, 67.7, 64.6, 48.8, 38.0, 33.5, 30.9, 25.2, 22.0, 14.2.

HR-MS: calculated for $C_{37}H_{31}ClFN_7O_8H$ $(M + H)^+$: 756.1979; found: 756.1947.

Purity was determined by qNMR using maleic acid as internal standard: 86.6%.

Mp: 157.8 °C

### Scheme S9: Synthesis of LP08[a]

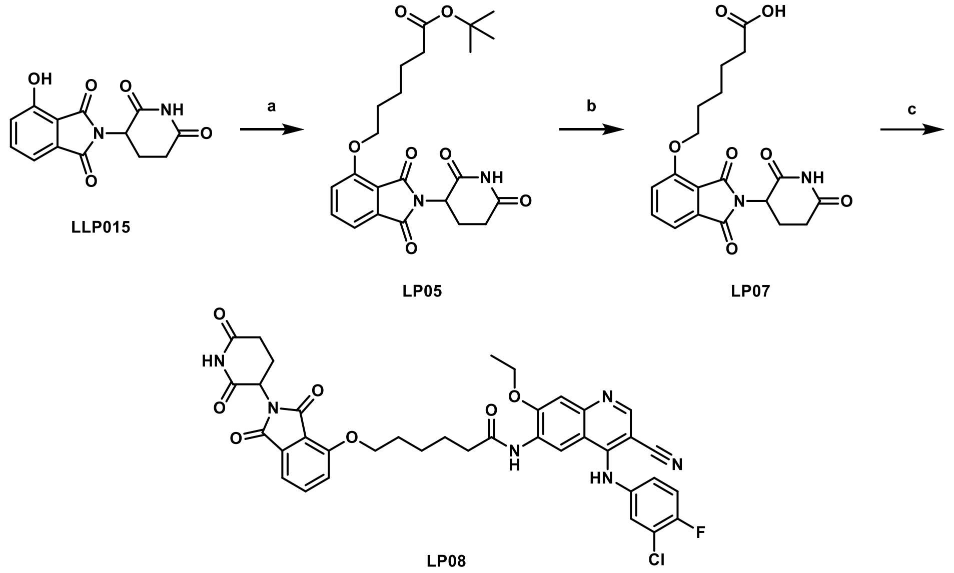

[a] Reagents and conditions: (a) *tert*-butyl 4-bromohexanoate, KI, $KHCO_3$, DMF, 60 °C, overnight, 69%[6]; (b) TFA, DCM, r.t., 3 h, 100%[6]; (c) **DH03**, HATU, DIPEA, DMSO, r.t., overnight, 8%.

### *tert*-Butyl 6-{[2-(2,6-dioxopiperidin-3-yl)-1,3-dioxoisoindolin-4-yl]oxy}hexanoate (LP05)

**LLP015** (1.50 g, 5.47 mmol, 1.00 eq) was dissolved in DMF (8.0 mL), *tert*-butyl 4-bromohexanoate (1.38 mL, 6.56 mmol, 1.20 eq), KI (0.09 g, 0.55 mmol, 0.10 eq) and $KHCO_3$ (0.83 g, 8.21 mmol, 1.50 eq) were added and the mixture stirred at 60 °C overnight. After cooling down to r.t., EtOAc (50 mL) was added and the mixture was washed with 5% LiCl solution (aq., 30 mL). The aqueous phase was extracted with EtOAc (2 x 50 mL). The combined organic phase was then washed with 5% LiCl solution (aq., 5 x 20 mL), brine (20 mL) and subsequently dried over $MgSO_4$. The solvent was removed under reduced pressure, then the residue was adsorbed on silica gel and purified by column chromatography (cHex/EtOAc 90:10 → cHex/EtOAc 50:50 over 30 min). The product **LP05** was obtained as a white solid (1.66 g, 3.74 mmol) in 69% yield.

$^1$H NMR (500 MHz, $CDCl_3$, 300 K): δ (ppm) = 8.23 (s, 1H), 7.66 (dd, $^3J$ = 8.6 Hz, $^3J$ = 7.5 Hz, 1H), 7.44 (dd, $^3J$ = 7.5 Hz, 1H), 7.20 (dd, $^3J$ = 8.6 Hz, 1H), 4.95 (dd, $^3J$ = 12.3 Hz, $^3J$ = 5.4 Hz, 1H), 4.17 (t, $^3J$ = 6.6 Hz, 2H), 2.90 - 2.69 (m, 3H), 2.25 (t, $^3J$ = 7.5 Hz, 2H), 2.13 - 2.09 (m, 1H), 1.89 (sm, 2H), 1.69 - 1.63 (m, 2H), 1.56 - 1.52 (m, 2H), 1.43 (s, 9H).

$^{13}$C NMR (125 MHz, $CDCl_3$, 300 K): δ (ppm) = 173.1, 171.1, 168.2, 167.2, 165.8, 156.8, 136.6, 133.9, 119.0, 117.2, 115.9, 80.3, 69.3, 49.2, 35.5, 31.5, 28.8, 28.2, 25.4, 24.8, 22.7.

HR-MS: calculated for $C_{23}H_{27}N_2O_7$ $(M - H)^-$: 443.1824; found: 443.1820.

### 6-{[2-(2,6-Dioxopiperidin-3-yl)-1,3-dioxoisoindolin-4-yl]oxy}hexanoic acid (LP07)

**LP05** (1.30 g, 2.92 mmol, 1.00 eq) and TFA (2.71 mL, 35.45 mmol, 12.14 eq) were dissolved in DCM (14.0 mL) and the mixture was stirred for 3 h at r.t. After removal of the solvent under reduced pressure and drying under high vacuum, the product **LP07** was obtained as a white solid (1.13 g, 2.92 mmol) in quantitative yield.

$^{1}$H NMR (500 MHz, DMSO-$d_6$, 300 K): δ (ppm) = 11.08 (s, 1H), 7.80 (dd, $^{3}J$ = 8.6 Hz, $^{3}J$ = 7.5 Hz, 1H), 7.50 (dd, $^{3}J$ = 8.6 Hz, 1H), 7.43 (dd, $^{3}J$ = 7.5 Hz, 1H), 5.07 (dd, $^{3}J$ = 12.3 Hz, $^{3}J$ = 5.4 Hz, 1H), 4.20 (t, $^{3}J$ = 6.6 Hz, 2H), 2.88 (ddd, $^{2}J$ = 17.2 Hz, $^{3}J$ = 13.8 Hz, $^{3}J$ = 5.4 Hz, 1H), 2.62 - 2.51 (m, 2H), 2.23 (t, $^{3}J$ = 7.5 Hz, 2H), 2.06 - 2.01 (m, 1H), 1.77 (sm, 2H), 1.58 (sm, 2H), 1.50 - 1.43 (m, 2H).

$^{13}$C NMR (125 MHz, DMSO-$d_6$, 300 K): δ (ppm) = 174.3, 172.7, 169.9, 166.8, 165.3, 156.0, 137.0, 133.2, 119.8, 116.2, 115.1, 68.7, 48.7, 33.6, 30.9, 28.1, 24.9, 24.1, 22.0.

HR-MS: calculated for $C_{19}H_{19}N_2O_7$ (M - H)$^-$: 387.1198; found: 387.1194.

### *N*-{4-[(3-Chloro-4-fluorophenyl)amino]-3-cyano-7-ethoxyquinolin-6-yl}-6-{[2-(2,6-dioxopiperidin-3-yl)-1,3-dioxoisoindolin-4-yl]oxy}hexanamide (LP08)

**LP07** (0.80 g, 2.06 mmol, 1.00 eq) was dissolved in dry DMSO (20.6 mL), **DH03** (0.81 g, 2.23 mmol, 1.10 eq), HATU (1.18 g, 3.09 mmol, 1.50 eq) and *N*,*N*-diisopropylethylamine (1.09 mL, 6.18 mmol, 3.00 eq) were added and the resulting mixture was stirred under an argon atmosphere for 72 h at r.t. After addition of EtOAc (100 mL), the reaction mixture was washed with $H_2O$ (5 x 40 mL), brine (30 mL) and finally dried over $MgSO_4$. The solvent was removed under reduced pressure, then the residue was adsorbed on silica gel and purified first by column chromatography (cHex/EtOAc 40:60 → cHex/EtOAc 0:100 over 30 min; EtOAc → EtOAc/MeOH 90:10 over 15 min) and then by preparative HPLC ($H_2O$/MeOH 95:5 → 5:95 over 60 min). The product **LP08** was obtained as a gray solid (114 mg, 0.16 mmol) in 8% yield.

$^{1}$H NMR (500 MHz, DMSO-$d_6$, 300 K): δ (ppm) = 11.08 (s, 1H), 9.67 (s, 1H), 9.27 (s, 1H), 8.86 (s, 1H), 8.53 (s, 1H), 7.80 (dd, $^{3}J$ = 8.6 Hz, $^{3}J$ = 7.5 Hz, 1H), 7.51 (d, $^{3}J$ = 8.6 Hz, 1H), 7.43 (d, $^{3}J$ = 7.5 Hz, 1H), 7.44 - 7-42 (m, 1H), 7.40 (dd, $^{3}J_{H-F}$ = 9.2 Hz, $^{3}J$ = 8.9 Hz, 1H), 7.39 (s, 1H), 7.23 - 7.22 (m, 1H), 5.06 (dd, $^{3}J$ = 12.6 Hz, $^{3}J$ = 5.4 Hz, 1H), 4.29 (q, $^{3}J$ = 7.2 Hz, 2H), 4.22 (t, $^{3}J$ = 6.3 Hz, 2H), 2.87 (ddd, $^{2}J$ = 17.2 Hz, $^{3}J$ = 13.8 Hz, $^{3}J$ = 5.4 Hz, 1H), 2.60 - 2.52 (m, 2H), 2.52 - 2.51 (m, 2H), 2.03 - 1.99 (m, 1H), 1.82 (sm, 2H), 1.71 (sm, 2H), 1.56 - 1.53 (m, 2H), 1.45 (t, $^{3}J$ = 7.2 Hz, 3H).

$^{13}$C NMR (125 MHz, DMSO-$d_6$, 300 K): δ (ppm) = 172.7, 171.6, 169.9, 166.8, 165.3, 156.0, 154.3 (d, $^{1}J_{C-F}$ = 244.7 Hz), 153.3, 151.9, 150.0, 147.7, 137.8 (d, $^{4}J_{C-F}$ = 3.6 Hz), 137.0, 133.2, 128.2, 124.7, 123.4 (d, $^{3}J_{C-}$

$_{F}$ = 6.0 Hz), 119.8, 119.5 (d, $^{2}J_{C\text{-}F}$ = 19.2 Hz), 117.1, 117.0 (d, $^{2}J_{C\text{-}F}$ = 21.6 Hz), 116.2, 115.1, 114.5, 113.8, 108.6, 88.8, 68.7, 64.5, 48.7, 36.0, 30.9, 28.2, 24.9, 24.8, 22.0, 14.2.

HR-MS: calculated for $C_{37}H_{31}ClFN_6O_7$ (M - H)$^{-}$: 725.1932; found: 725.1934.

Purity was determined by qNMR using maleic acid as internal standard: 90.1%.

Mp: 152.9 °C

### Scheme S10: Synthesis of LP04[a]

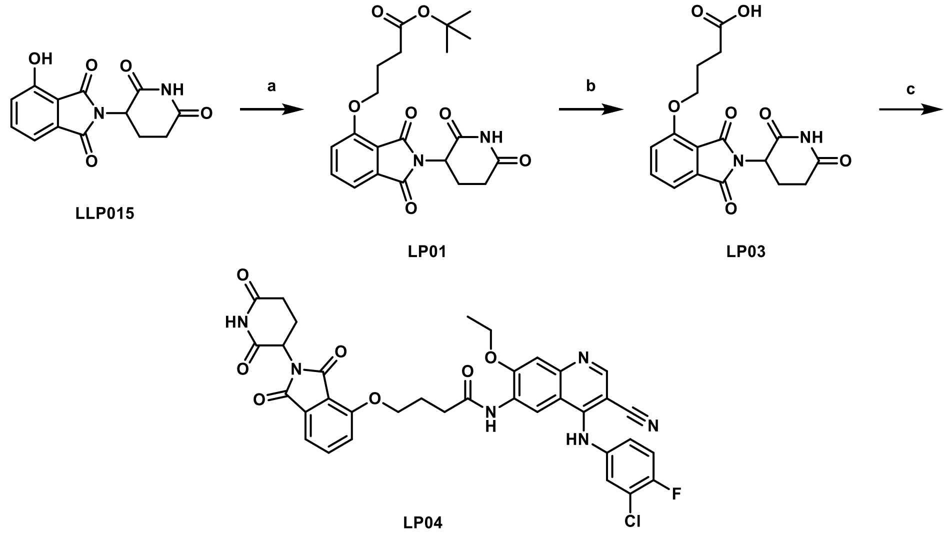

[a] Reagents and conditions: (a) *tert*-butyl 4-bromobutanoate, KI, $K_2CO_3$, DMF, 60 °C, overnight, 66% [7]; (b) TFA, DCM, r.t., 3 h, 100% [7]; (c) **DH03**, HATU, DIPEA, DMSO, r.t., overnight, 33%.

### *tert*-Butyl 4-((2-(2,6-dioxopiperidin-3-yl)-1,3-dioxoisoindolin-4-yl)oxy)butanoate (LP01)

**LLP015** (2.00 g, 7.29 mmol, 1.00 eq) was dissolved in DMF (10.0 mL), *tert*-butyl 4-bromobutanoate (1.55 mL, 8.75 mmol, 1.20 eq), KI (0.12 g, 0.73 mmol, 0.10 eq) and $K_2CO_3$ (1.51 g, 10.94 mmol, 1.50 eq) were added and the mixture stirred at 60 °C overnight. Then, EtOAc (50 mL) was added and the mixture was washed with 5% LiCl solution (aq., 30 mL). The aqueous phase was extracted with EtOAc (2 x 50 mL). The combined organic phase was then washed again with 5% LiCl solution (aq., 5 x 20 mL), brine (20 mL) and at the end dried over $MgSO_4$. The solvent was removed under reduced pressure, then the residue was adsorbed on silica gel and purified by column chromatography (cHex/EtOAc 90:10 → cHex/EtOAc 0:100 over 60 min). The product **LP01** was obtained as a white solid (2.01 g, 4.82 mmol) in 66% yield.

$^1$H NMR (500 MHz, DMSO-$d_6$, 300 K): δ (ppm) = 11.08 (s, 1H), 7.81 (dd, $^3J$ = 8.6 Hz, $^3J$ = 7.2 Hz, 1H), 7.50 (dd, $^3J$ = 8.6 Hz, 1H), 7.45 (dd, $^3J$ = 7.2 Hz, 1H), 5.08 (dd, $^3J$ = 12.9 Hz, $^3J$ = 5.4 Hz, 1H), 4.22 (t, $^3J$ = 6.3 Hz, 2H), 2.88 (ddd, $^2J$ = 17.2 Hz, $^3J$ = 13.8 Hz, $^3J$ = 5.4 Hz, 1H), 2.62 - 2.51 (m, 2H), 2.43 (t, $^3J$ = 7.2 Hz, 2H), 2.06 - 2.01 (m, 1H), 1.97 (sm, 2H), 1.39 (s, 9H).

$^{13}$C NMR (125 MHz, DMSO-$d_6$, 300 K): δ (ppm) = 172.7, 171.8, 169.8, 166.8, 165.3, 155.8, 137.0, 133.2, 119.8, 116.4, 115.3, 79.7, 67.8, 48.7, 30.9, 30.9, 27.7, 24.1, 22.0.

HR-MS: calculated for $C_{21}H_{23}N_2O_7$ $(M - H)^-$: 415.1511; found: 415.1507.

#### 4-{[2-(2,6-Dioxopiperidin-3-yl)-1,3-dioxoisoindolin-4-yl]oxy}butanoic acid (LP03)

**LP01** (1.80 g, 4.32 mmol, 1.00 eq) and TFA (4.02 mL, 52.48 mmol, 12.14 eq) were dissolved in DCM (20.0 mL) and the mixture was stirred for 3 h at r.t. The product, **LP03**, was obtained as an off-white solid (1.56 g, 4.32 mmol) in quantitative yield after removal of the solvent under reduced pressure and drying under high vacuum.

$^{1}$H NMR (500 MHz, DMSO-$d_6$, 300 K): δ (ppm) = 11.08 (s, 1H), 7.81 (dd, $^{3}J$ = 8.6 Hz, $^{3}J$ = 7.2 Hz, 1H), 7.51 (dd, $^{3}J$ = 8.6 Hz, 1H), 7.45 (dd, $^{3}J$ = 7.2 Hz, 1H), 5.08 (dd, $^{3}J$ = 12.9 Hz,$^{3}J$ = 5.4 Hz, 1H), 4.23 (t, $^{3}J$ = 6.3 Hz, 2H), 2.88 (ddd, $^{2}J$ = 17.2 Hz, $^{3}J$ = 13.8 Hz, $^{3}J$ = 5.4 Hz, 1H), 2.62 - 2.51 (m, 2H), 2.45 (t, $^{3}J$ = 7.5 Hz, 2H), 2.06 - 2.01 (m, 1H), 1.98 (sm, 2H).

$^{13}$C NMR (125 MHz, DMSO-$d_6$, 300 K): δ (ppm) = 174.0, 172.7, 169.9, 166.8, 165.3, 155.8, 137.0, 133.2, 119.8, 116.4, 115.3, 67.9, 48.8, 30.9, 29.7, 24.0, 22.0.

HR-MS: calculated for $C_{17}H_{16}N_2O_7H$ (M + H)$^+$: 361.1030; found: 361.1017.

#### *N*-{4-[(3-Chloro-4-fluorophenyl)amino]-3-cyano-7-ethoxyquinolin-6-yl}-4-{[2-(2,6-dioxopiperidin-3-yl)-1,3-dioxoisoindolin-4-yl]oxy}butanamide (LP04)

**LP03** (0.40 g, 1.11 mmol, 1.00 eq) was dissolved in dry DMSO (11.1 mL), **DH03** (0.44 g, 1.22 mmol, 1.10 eq), HATU (0.63 g, 1.67 mmol, 1.50 eq) and *N*,*N*-diisopropylethylamine (0.58 mL, 3.33 mmol, 3.00 eq) were added and the resulting mixture was stirred under an argon atmosphere overnight at r.t. Upon addition of EtOAc (50 mL), the mixture was washed with $H_2O$ (5 x 30 mL), brine (20 mL) and finally dried over $MgSO_4$. The solvent was removed under reduced pressure, then the residue was adsorbed on silica gel and purified by column chromatography (cHex/EtOAc 30:70 → 0:100 over 18 min; EtOAc → EtOAc/MeOH 90:10 over 15 min). The product **LP04** was obtained as a yellowish solid (254 mg, 0.36 mmol) in 33% yield.

$^{1}$H NMR (500 MHz, DMSO-$d_6$, 300 K): δ (ppm) = 11.08 (s, 1H), 9.67 (bs, 1H), 9.38 (s, 1H), 8.85 (s, 1H), 8.54 (s, 1H), 7.81 (dd, $^{3}J$ = 8.6 Hz, $^{3}J$ = 7.2 Hz, 1H), 7.53 (d, $^{3}J$ = 8.6 Hz, 1H), 7.45 (d, $^{3}J$ = 7.2 Hz, 1H), 7.45 (dd, $^{4}J_{H\text{-}F}$ = 6.6 Hz, $^{4}J$ = 2.9 Hz, 1H), 7.41 (dd, $^{3}J$ = 9.2 Hz, $^{3}J$ = 8.9 Hz, 1H), 7.40 (s, 1H), 7.24 (ddd, $^{3}J$ = 8.9 Hz, $^{4}J_{H\text{-}F}$ = 4.3 Hz, $^{4}J$ = 2.6 Hz, 1H), 5.08 (dd, $^{3}J$ = 12.9 Hz, $^{3}J$ = 5.4 Hz, 1H), 4.32 (m, 4H), 2.88 (ddd, $^{2}J$ = 17.2 Hz, $^{3}J$ = 13.8 Hz, $^{3}J$ = 5.4 Hz, 1H), 2.71 (t, $^{3}J$ = 7.2 Hz, 2H), 2.61 - 2.53 (m, 2H), 2.14 - 2.09 (m, 2H), 2.02 (sm, 1H), 1.43 (t, $^{3}J$ = 7.2 Hz, 3H).

$^{13}$C NMR (125 MHz, DMSO-$d_6$, 300 K): δ (ppm) = 172.7, 171.1, 169.9, 166.8, 165.3, 155.8, 154.3 (d, $^{1}J_{C\text{-}F}$ = 243.5 Hz), 153.5, 151.9, 150.0, 147.8, 137.7 (d, $^{4}J_{C\text{-}F}$ = 3.6 Hz), 137.0, 133.2, 128.1, 124.7, 123.5 (d, $^{3}J_{C\text{-}F}$ = 7.2 Hz), 119.8, 119.5 (d, $^{2}J_{C\text{-}F}$ = 18.0 Hz), 117.1, 117.0 (d, $^{2}J_{C\text{-}F}$ = 21.6 Hz), 116.3, 115.3, 114.7, 113.7, 108.6, 88.7, 68.1, 64.5, 48.7, 32.1, 30.9, 24.4, 22.0, 14.2.

HR-MS: calculated for $C_{35}H_{27}ClFN_6O_7$ $(M - H)^-$: 697.1619; found: 697.1624.

Purity was determined by qNMR using maleic acid as internal standard: 93.4%.

Mp: 157.0 °C

### Scheme S11: Synthesis of LB06[a]

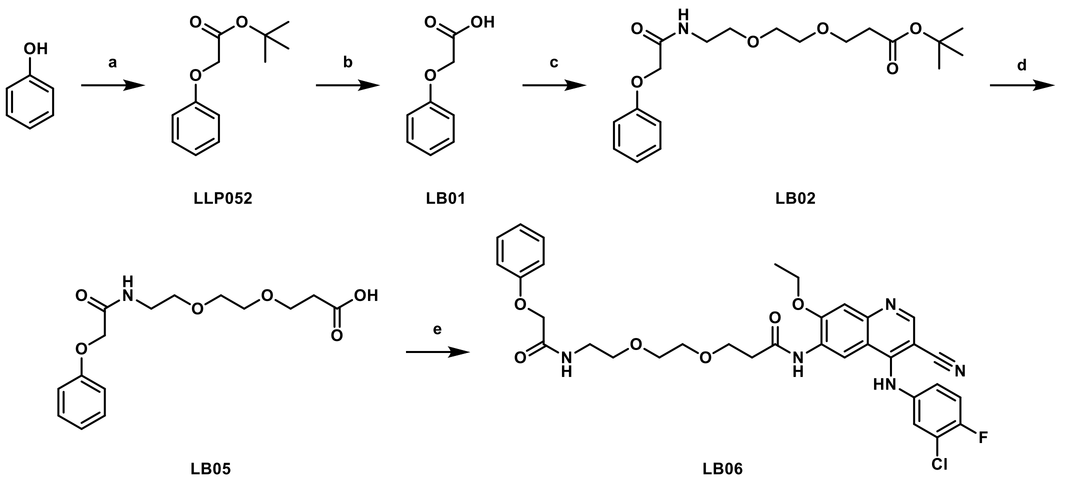

[a] Reagents and conditions: (a) *tert*-butyl 2-bromoacetate, KI, $KHCO_3$, DMF, r.t., 2 h, 80% [7]; (b) TFA, r.t., 2 h, 87%[6]; (c) *tert*-butyl 3-[2-(2-aminoethoxy)ethoxy]propanoate, HATU, DIPEA, DMF, r.t., overnight, 49%; (d) TFA, r.t., 2 h; (e) **DH03**, HATU, DIPEA, DMF, r.t., overnight, 7%.

#### *tert*-Butyl 2-phenoxyacetate (LLP052)

Phenol (0.69 g, 7.29 mmol, 1.00 eq) was dissolved in DMF (10.0 mL), *tert*-butyl bromoacetate (1.08 mL, 7.29 mmol, 1.00 eq), KI (0.12 g, 0.73 mmol, 0.10 eq) and $KHCO_3$ (1.11 g, 10.94 mmol, 1.50 eq) were added and the mixture stirred for 5 h at 60 °C. After cooling to r.t., EtOAc (50 mL) was added and the mixture was washed with 5% LiCl solution (aq., 5 x 30 mL), brine (20 mL) and at the end dried over $MgSO_4$. The solvent was removed under reduced pressure, then the residue was adsorbed on silica gel and purified by column chromatography (cHex → cHex/EtOAc 80:20 over 20 min). The product **LLP052** was obtained as a colorless oil (1.28 g, 6.14 mmol) in 84% yield.

$^1$H NMR (400 MHz, DMSO-*d₆*, 300 K): δ (ppm) = 7.31 - 7.26 (m, 2H), 6.95 (sm, 1H), 6.91 - 6.87 (m, 2H), 4.63 (s, 2H), 1.42 (s, 9H).

$^{13}$C NMR (100 MHz, DMSO-$d_6$, 300 K): δ (ppm) = 167.8, 157.6, 129.4, 121.0, 114.4, 81.3, 64.9, 27.7.

HR-MS: calculated for $C_{12}H_{16}O_3Na$ $(M + Na)^+$: 231.0992; found: 231.0989.

Spectral data are in accordance with literature [8].

#### 2-Phenoxyacetic acid (LB01)

**LLP052** (1.10 g, 5.28 mmol, 1.00 eq) was dissolved in DCM (10.0 mL), TFA (8.14 mL, 105.60 mmol, 20.00 eq) was added and stirred for 2 h at r.t. After addition of DCM (20 mL), the mixture was washed with $H_2O$ (20 mL) and dried over $MgSO_4$. The solvent was then removed under reduced pressure and the product **LB01** was obtained as a white solid (0.70 g, 4.59 mmol) in 87% yield.

$^{1}H$ NMR (300 MHz, $CDCl_3$, 300 K): δ (ppm) = 8.59 (bs, 1H), 7.36 - 7.29 (m, 2H), 7.06 - 7.00 (sm, 1H), 6.95 - 6.91 (m, 2H), 4.70 (s, 2H).

HR-MS: calculated for $C_8H_7O_3$ $(M - H)^-$: 151.0401; found: 151.0388.

Spectral data are in accordance with literature [8].

### *tert*-Butyl 3-{2-[2-(2-phenoxyacetamido)ethoxy]-ethoxy}propanoate (LB02)

**LB01** (0.60 g, 3.94 mmol, 1.00 eq) was dissolved in dry DMF (5.0 mL), mixed with *tert*-butyl 3-[2-(2-aminoethoxy)ethoxy]propanoate (1.00 mL, 4.34 mmol, 1.10 eq), HATU (2.25 g, 5.92 mmol, 1.50 eq) and *N*,*N*-diisopropylethylamine (2.01 mL, 11.83 mmol, 3.00 eq) and stirred under an argon atmosphere overnight at r.t. After addition of EtOAc (50 mL), the mixture was washed with 5% LiCl solution (aq., 5 x 30 mL), brine (30 mL) and then dried over $MgSO_4$. The solvent was removed under reduced pressure, the residue was adsorbed on silica gel and purified by column chromatography (cHex/EtOAc 60:40 → cHex/EtOAc 20:80 over 30 min). The product **LB02** was obtained as a colorless oil (0.70 g, 1.91 mmol) in 49% yield.

$^{1}H$ NMR (400 MHz, DMSO-*d*$_6$, 300 K): δ (ppm) = 8.01 (t, $^{3}J$ = 5.7 Hz, 1H), 7.32 - 7.28 (m, 2H, 3-H), 6.98 - 6.94 (m, 3H), 4.46 (s, 2H), 3.58 (t, $^{3}J$ = 6.2 Hz, 2H), 3.48 (ps, 4H), 3.44 (t, $^{3}J$ = 6.0 Hz, 2H), 3.31 - 3.26 (m, 2H), 2.41 (t, $^{3}J$ = 6.2 Hz, 2H), 1.39 (s, 9H).

$^{13}C$ NMR (100 MHz, DMSO-*d*$_6$, 300 K): δ (ppm) = 170.3, 167.6, 157.7, 129.4, 121.1, 114.7, 79.7, 69.6, 69.4, 68.7, 66.9, 66.2, 38.2, 35.8, 27.7.

HR-MS: calculated for $C_{19}H_{29}NO_6Na$ $(M + Na)^+$: 390.1887; found: 390.1879.

Purity was determined by qNMR using 1,2,4,5-tetrachloro-3-nitrobenzene as internal standard: 98.8%.

### 3-{2-[2-(2-Phenoxyacetamido)ethoxy]-ethoxy}propanoic acid (LB05)

**LB02** (0.60 g, 1.63 mmol, 1.00 eq) and TFA (3.14 mL, 40.75 mmol, 25.00 eq) were dissolved in DCM (10.0 mL) and the reaction mixture was stirred for 2 h at r.t. **LB05** was obtained as an off-white solid (0.51 g, 1.63 mmol) in quantitative yield after removal of the solvent under reduced pressure and drying under high vacuum.

$^{1}H$ NMR (400 MHz, DMSO-*d*$_6$, 300 K): δ (ppm) = 8.02 (t, $^{3}J$ = 5.3 Hz, 1H), 7.32 - 7.28 (m, 2H), 6.98 - 6.94 (m, 3H), 4.47 (s, 2H), 3.60 (t, $^{3}J$ = 6.4 Hz, 2H), 3.48 (ps, 4H), 3.45 (t, $^{3}J$ = 6.0 Hz, 2H), 3.29 (sm, 2H), 2.43 (t, $^{3}J$ = 6.4 Hz, 2H).

$^{13}C$ NMR (100 MHz, DMSO-*d*$_6$, 300 K): δ (ppm) = 172.6, 167.7, 157.7, 129.4, 121.2, 114.7, 69.5, 69.4, 68.8, 66.9, 66.2, 38.2, 34.7.

HR-MS: calculated for $C_{15}H_{20}NO_6$ $(M - H)^-$: 310.1296; found: 310.1294.

### *N*-{4-[(3-Chloro-4-fluorophenyl)amino]-3-cyano-7-ethoxyquinolin-6-yl}-3-{2-[2-(2-phenoxyacetamido)ethoxy]ethoxy}propanamide (LB06)

**LB05** (0.40 g, 1.28 mmol, 1.00 eq) was dissolved in dry DMSO (12.8 mL), **DH03** (0.55 g, 1.54 mmol, 1.20 eq), HATU (0.73 g, 1.92 mmol, 1.50 eq) and *N*,*N*-diisopropylethylamine (0.65 mL, 3.85 mmol, 3.00 eq) were added and the resulting mixture was stirred under an argon atmosphere overnight at r.t. After addition of $H_2O$ (30 mL), the mixture was extracted with EtOAc (2 x 70 mL). The combined organic phase was then washed with $H_2O$ (4 x 30 mL) and Brine (30 mL) and dried over $MgSO_4$. The solvent was removed under reduced pressure, then the residue was adsorbed on silica gel and purified first by column chromatography (cHex/EtOAc 50:50 → 0:100 over 25 min; EtOAc → EtOAc/MeOH 90:10 over 15 min) and then by preparative HPLC ($H_2O$/MeCN 95:5 → 5:95 over 60 min). The product **LB06** was obtained as a light-yellow solid (60 mg, 0.09 mmol) in 7% yield.

$^1$H NMR (400 MHz, DMSO-$d_6$, 300 K): δ (ppm) = 9.86 (bs, 1H), 9.38 (s, 1H), 8.97 (s, 1H), 8.60 (s, 1H), 8.01 (t, $^3J$ = 5.7 Hz, 1H), 7.46 (dd, $^4J_{H-F}$ = 6.6 Hz, $^4J$ = 2.7 Hz, 1H), 7.41 (dd, $^3J_{H-F}$ = 9.2 Hz, $^3J$ = 8.9 Hz, 1H), 7.40 (s, 1H), 7.30 - 7.22 (m, 3H), 6.97 - 6.92 (m, 3H), 4.44 (s, 2H), 4.32 (q, $^3J$ = 6.9 Hz, 2H), 3.74 (t, $^3J$ = 6.2 Hz, 2H), 3.57 - 3.51 (m, 4H), 3.45 (t, $^3J$ = 6.0 Hz, 2H), 3.27 (sm, 2H), 2.89 (ddd, $^2J$ = 17.2 Hz, $^3J$ = 13.8 Hz, $^3J$ = 5.4 Hz, 1H), 2.73 (t, $^3J$ = 6.2 Hz, 2H), 1.47 (t, $^3J$ = 6.9 Hz, 3H).

$^{13}$C NMR (100 MHz, DMSO-$d_6$, 300 K): δ (ppm) = 169.8, 167.7, 157.6, 154.5 (d, $^1J_{C-F}$ = 243.7 Hz), 153.1, 151.3, 150.4, 146.2, 137.7 (d, $^4J_{C-F}$ = 3.9 Hz), 129.4, 128.3, 124.9, 123.7 (d, $^3J_{C-F}$ = 6.7 Hz), 121.1, 119.5 (d, $^2J_{C-F}$ = 19.3 Hz), 117.0 (d, $^2J_{C-F}$ = 22.2 Hz), 116.7, 114.6, 113.8, 107.6, 88.8, 69.6, 69.4, 68.8, 66.9, 66.5, 64.7, 38.2, 36.9, 14.2.

HR-MS: calculated for $C_{33}H_{33}ClFN_5O_6H$ $(M + H)^+$: 650.2176; found: 650.2173.

Purity was determined by qNMR using methyl sulfone as internal standard: 95.2%.

Mp: 80.1 °C

$^1H$ and $^{13}C$ NMR spectra of the PROTACs

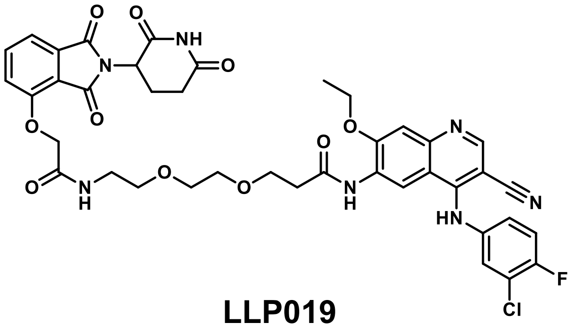

**LLP019**

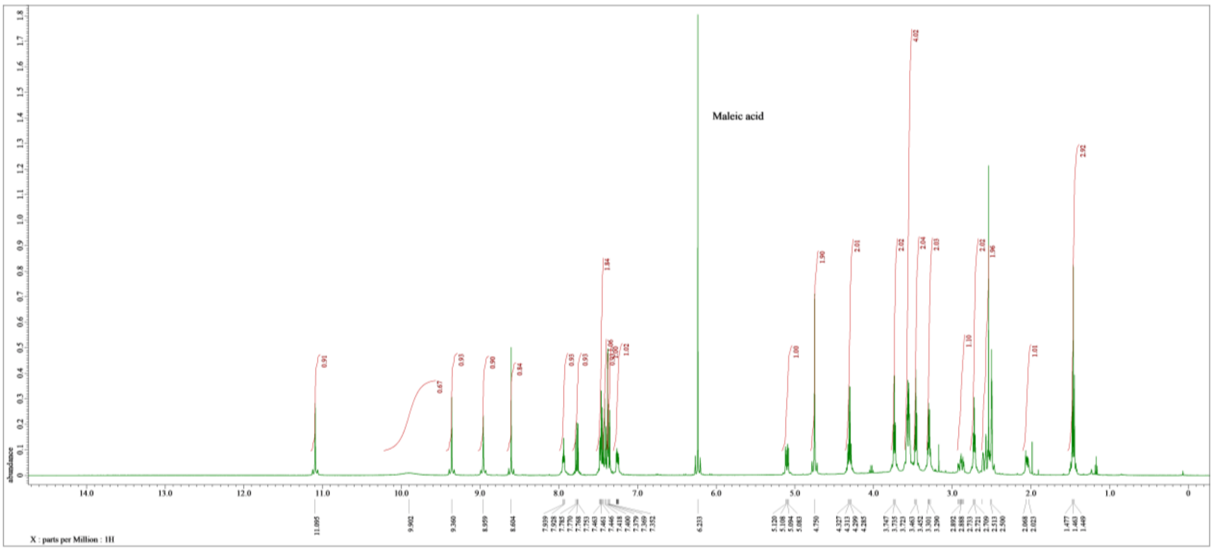

**Figure S8:** $^1H$ NMR spectrum of **LLP019**, 500 MHz, solvent: DMSO-$d_6$, with internal standard maleic acid

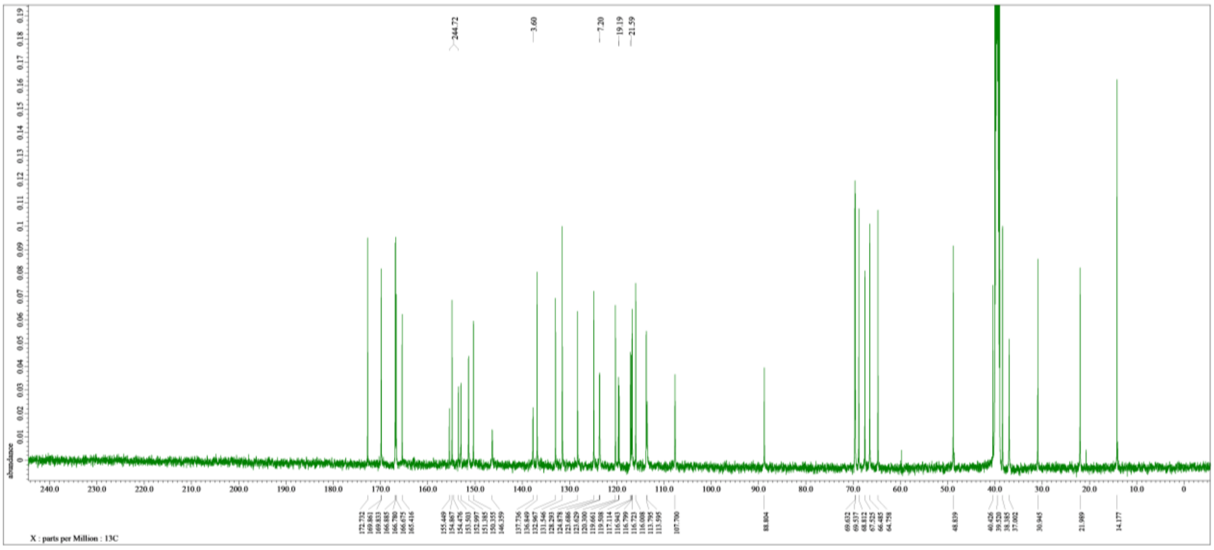

**Figure S9:** $^{13}C$ NMR spectrum of **LLP019**, 125 MHz, solvent: DMSO-$d_6$

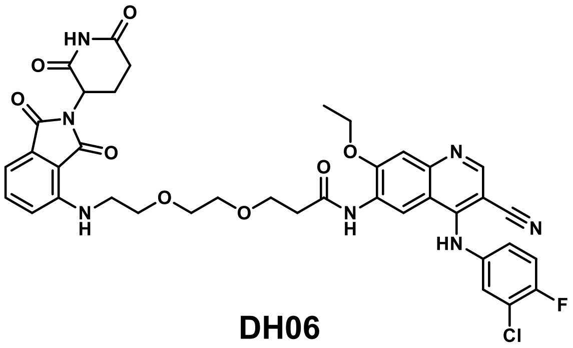

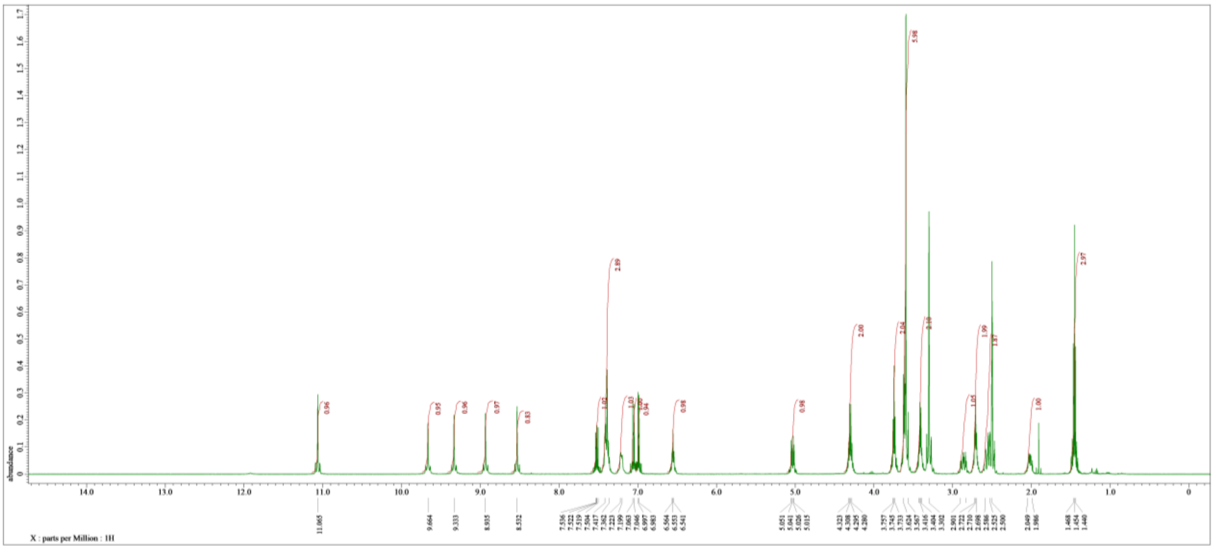

**Figure S10:**[1]H NMR spectrum of **DH06**, 500 MHz, solvent: DMSO-*d₆*

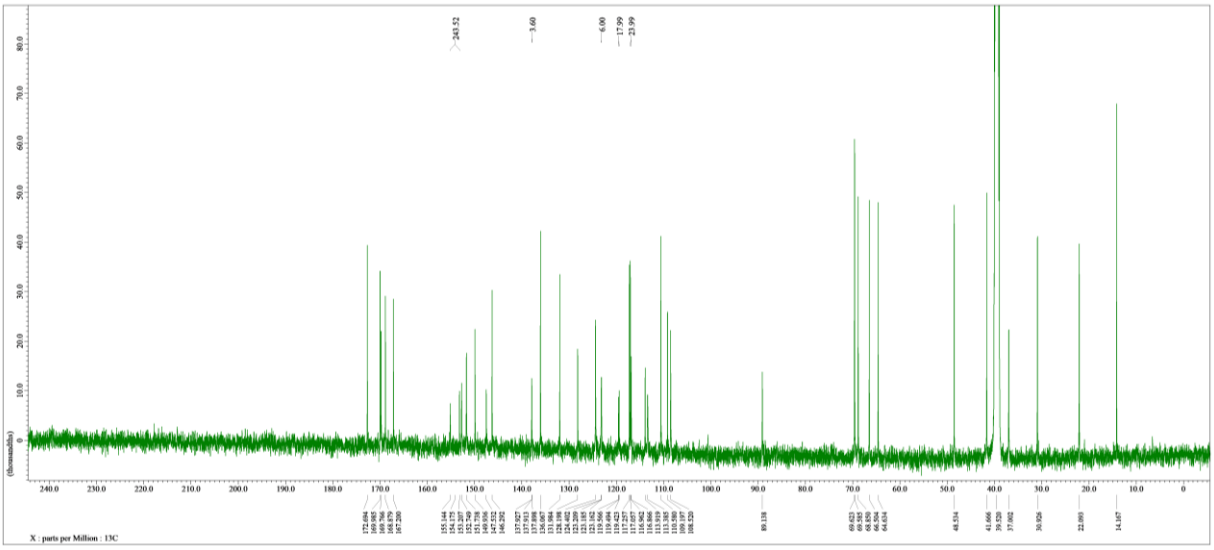

**Figure S11:** [13]C NMR spectrum of **DH06**, 125 MHz, solvent: DMSO-*d₆*

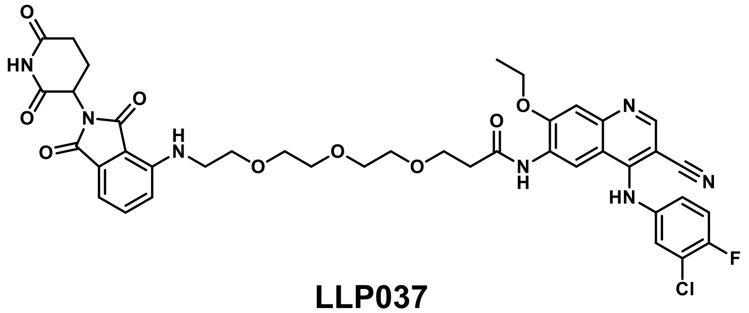

**LLP037**

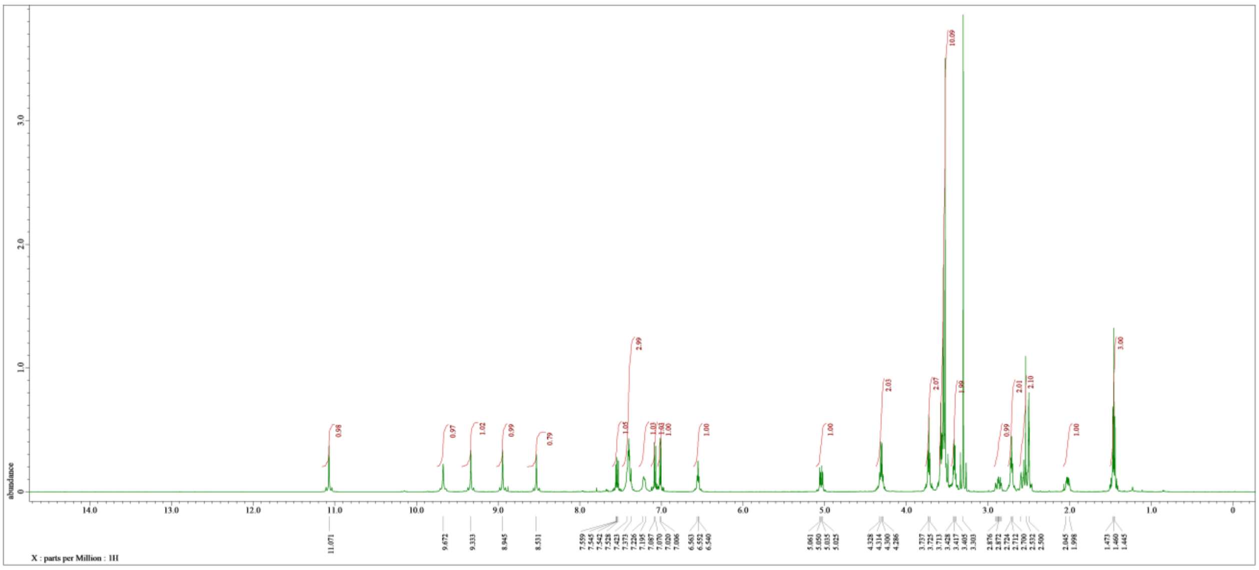

**Figure S12:** $^{1}$H NMR spectrum of **LLP037**, 500 MHz, solvent: DMSO-$d_6$

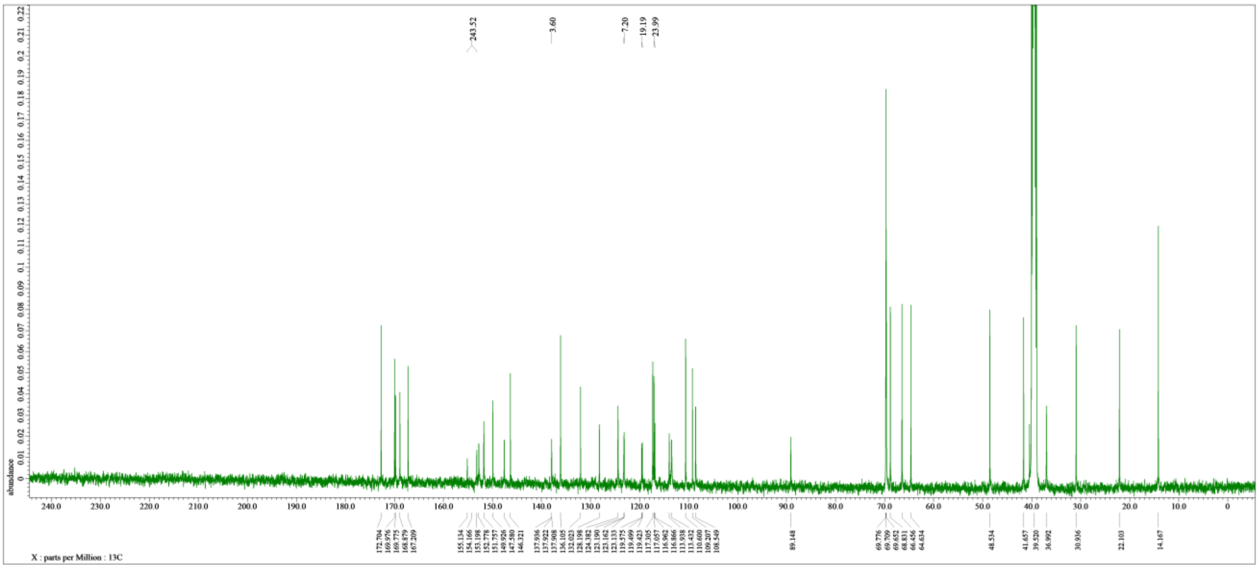

**Figure S13:** $^{13}$C NMR spectrum of **LLP037**, 125 MHz, solvent: DMSO-$d_6$

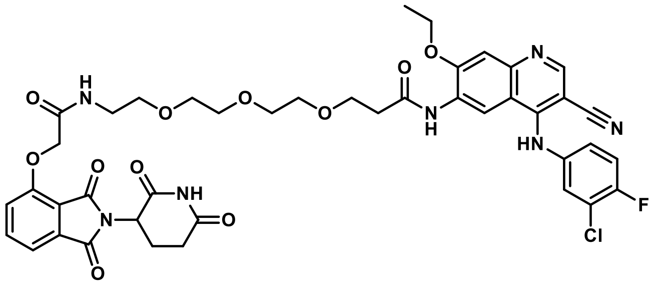

**LLP038**

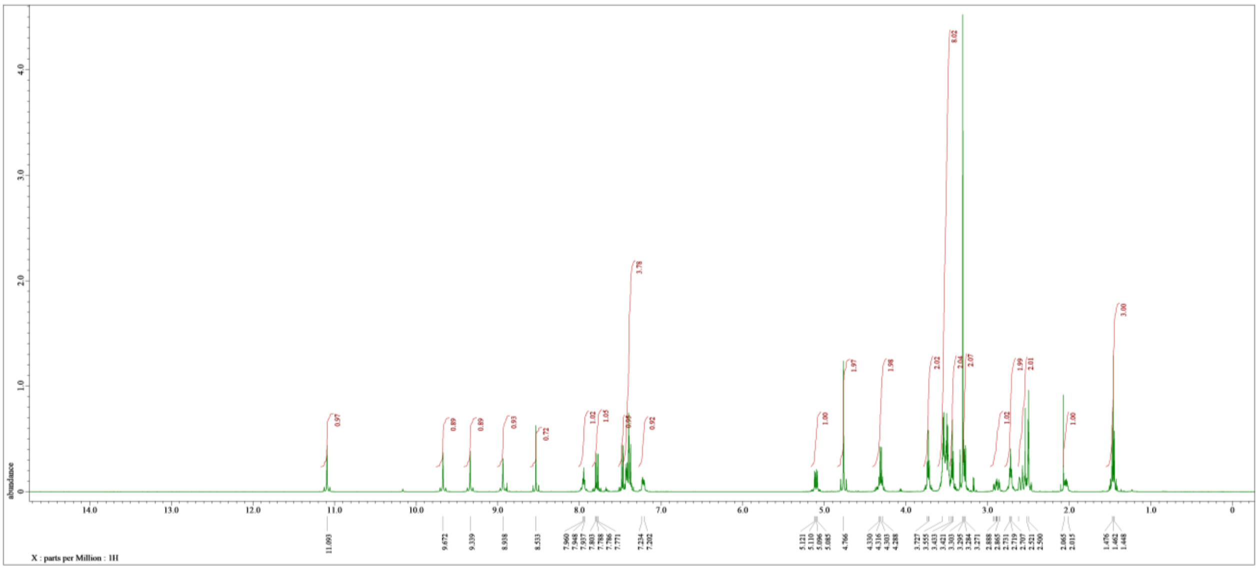

**Figure S14:** $^{1}$H NMR spectrum of **LLP038**, 500 MHz, solvent: DMSO-$d_6$

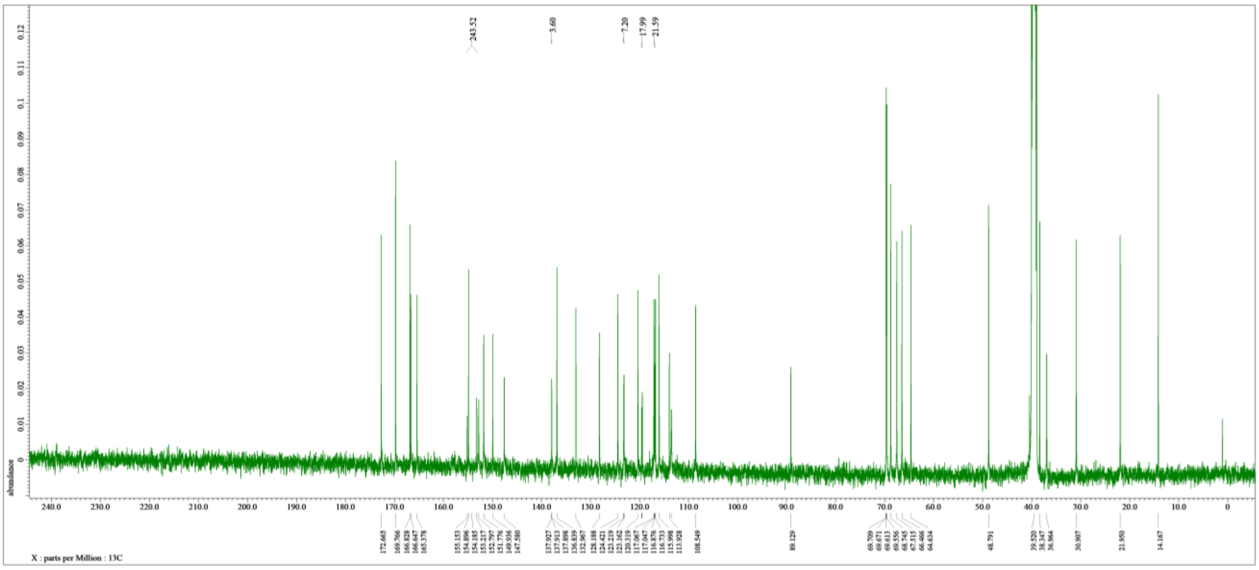

**Figure S15:** $^{13}$C NMR spectrum of **LLP038**, 125 MHz, solvent: DMSO-$d_6$

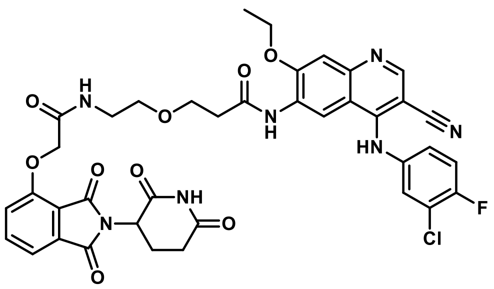

**LLP031**

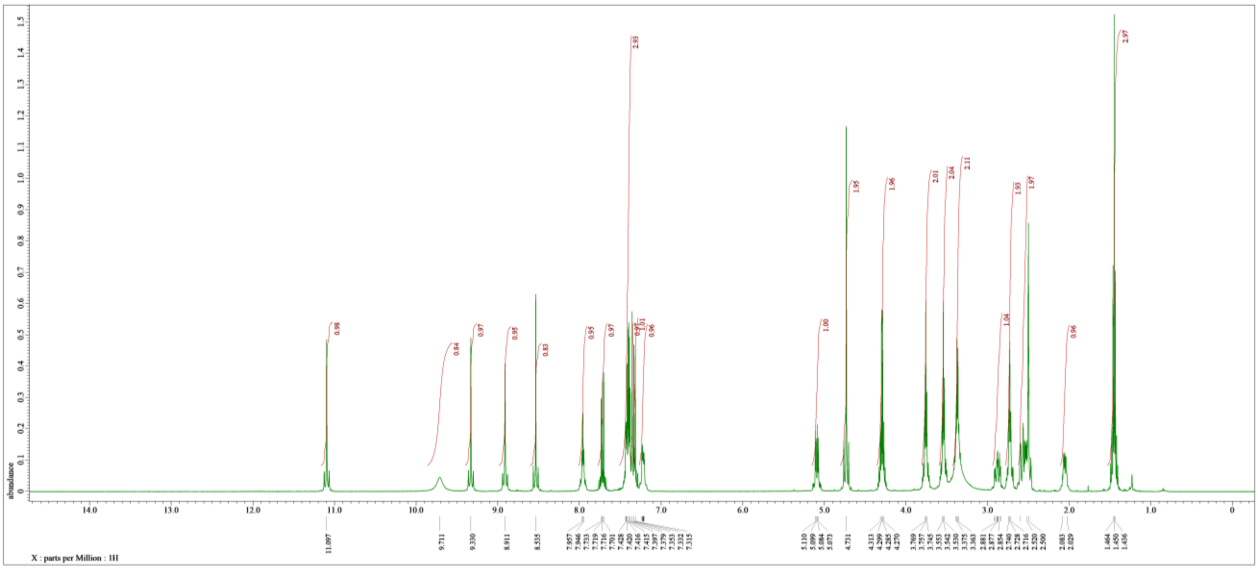

**Figure S16:** $^{1}$H NMR spectrum of **LLP031**, 500 MHz, solvent: DMSO-$d_6$

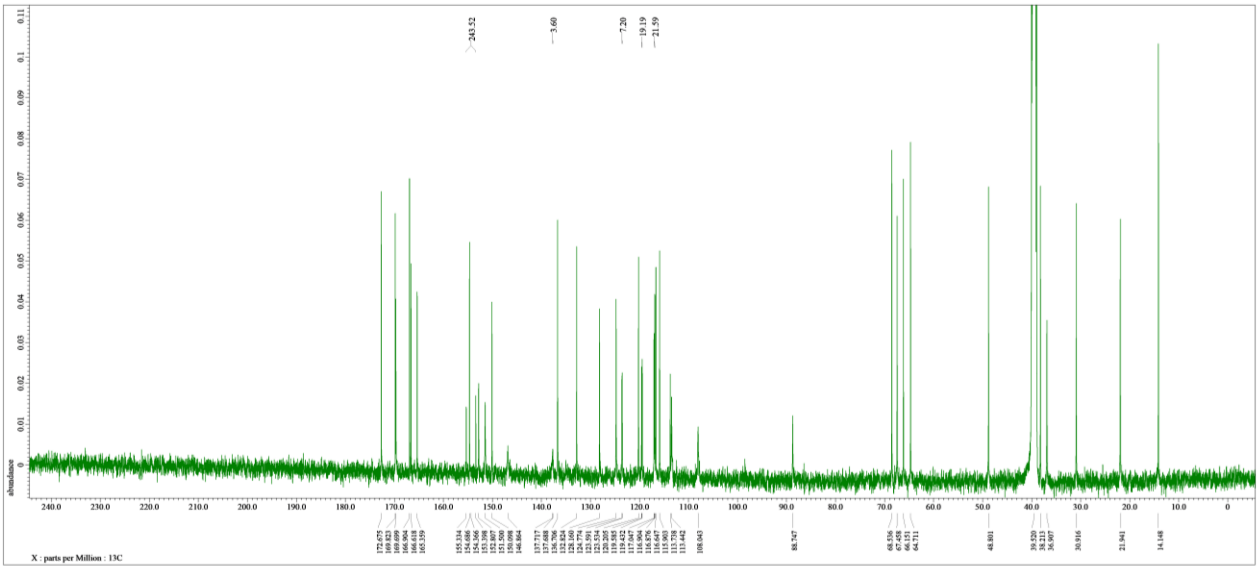

**Figure S17:** $^{13}$C NMR spectrum of **LLP031**, 125 MHz, solvent: DMSO-$d_6$

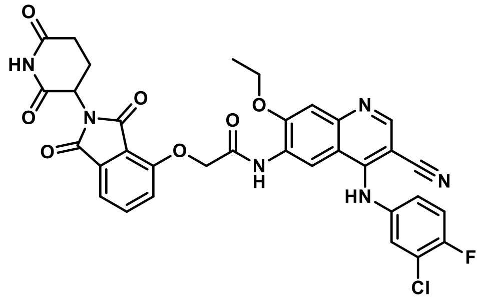

**LLP049**

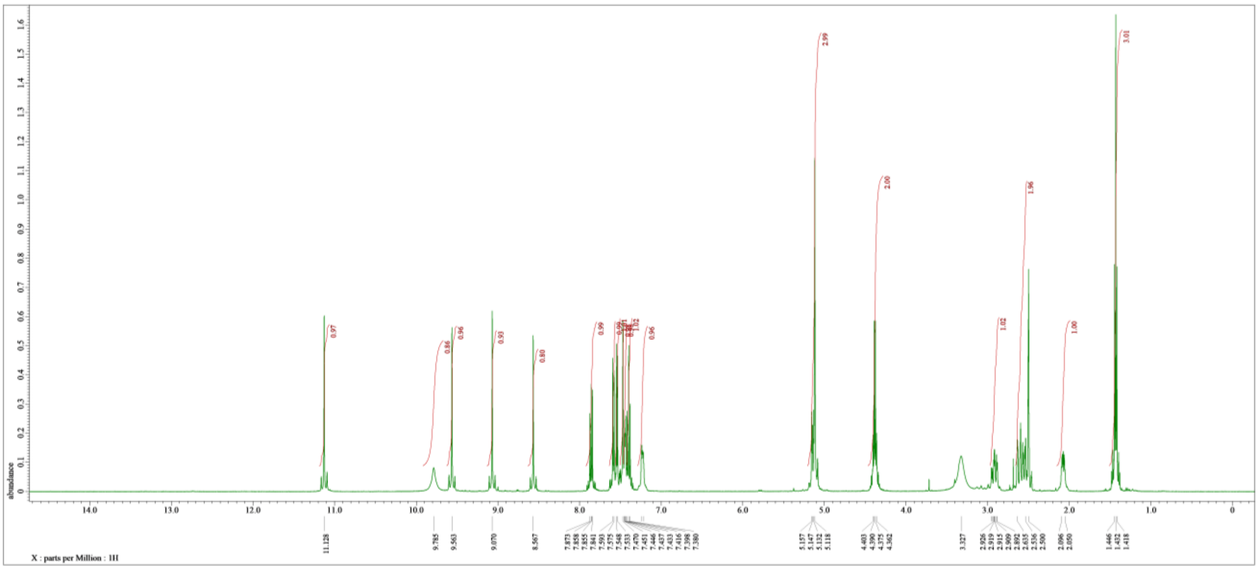

**Figure S18:** $^{1}$H NMR spectrum of **LLP049**, 500 MHz, solvent: DMSO-$d_6$

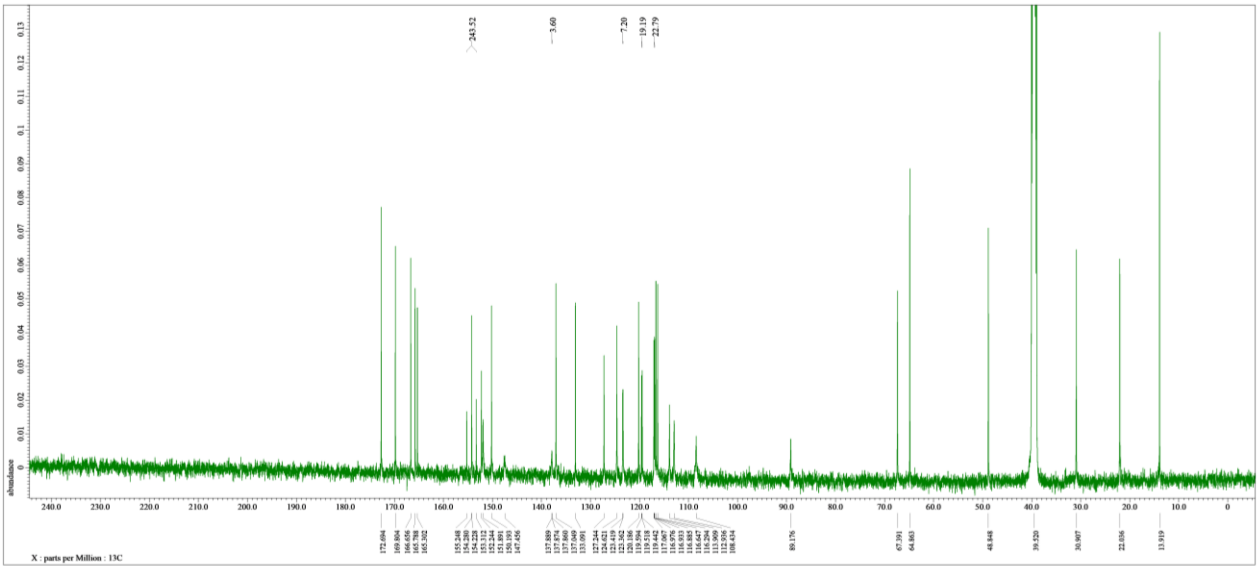

**Figure S19:** $^{13}$C NMR spectrum of **LLP049**, 125 MHz, solvent: DMSO-$d_6$

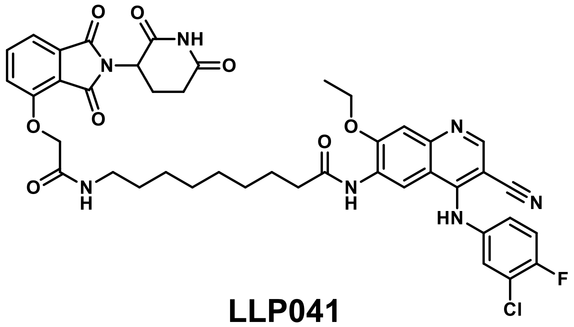

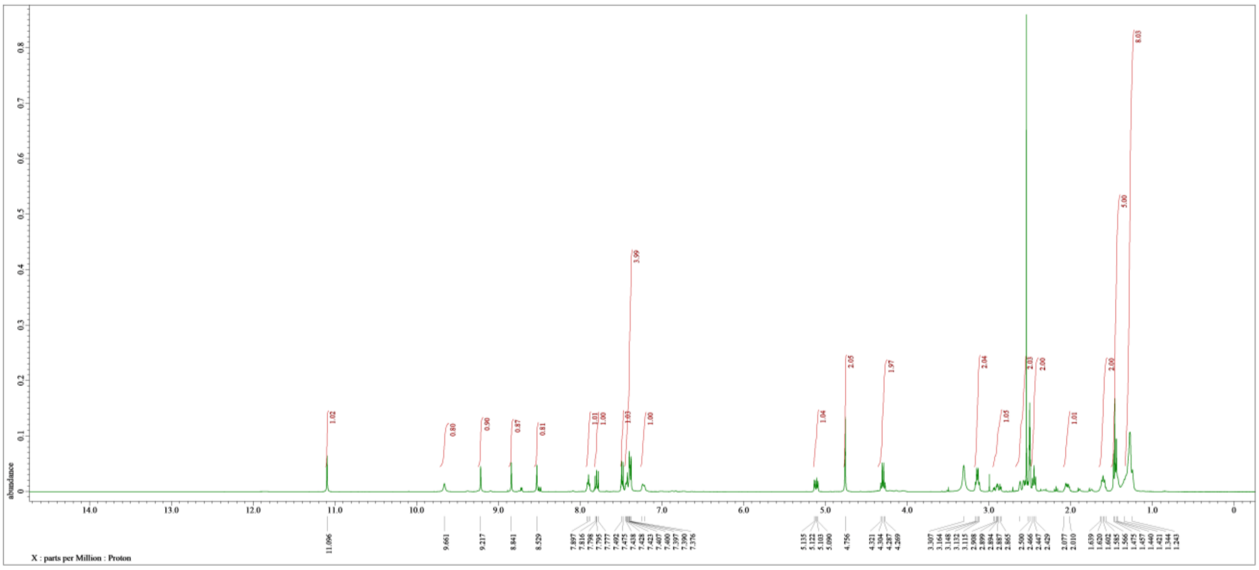

**Figure S20:** $^{1}$H NMR spectrum of **LLP041**, 400 MHz, solvent: DMSO-$d_6$

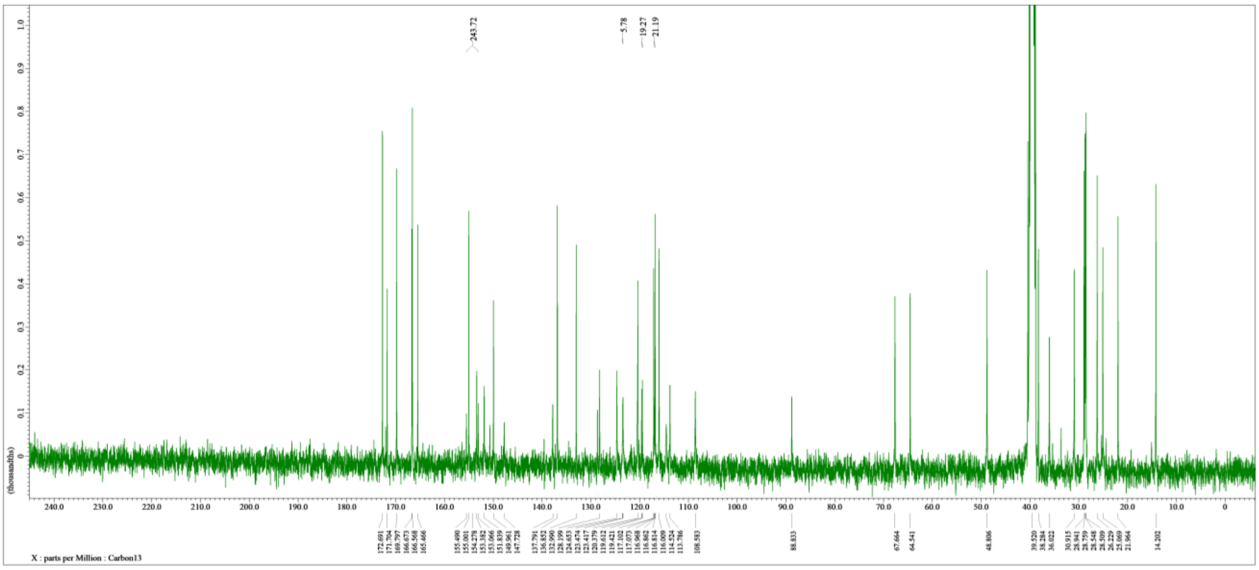

**Figure S21:** $^{13}$C NMR spectrum of **LLP041**, 100 MHz, solvent: DMSO-$d_6$

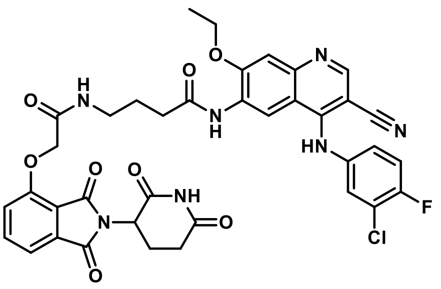

**LP15**

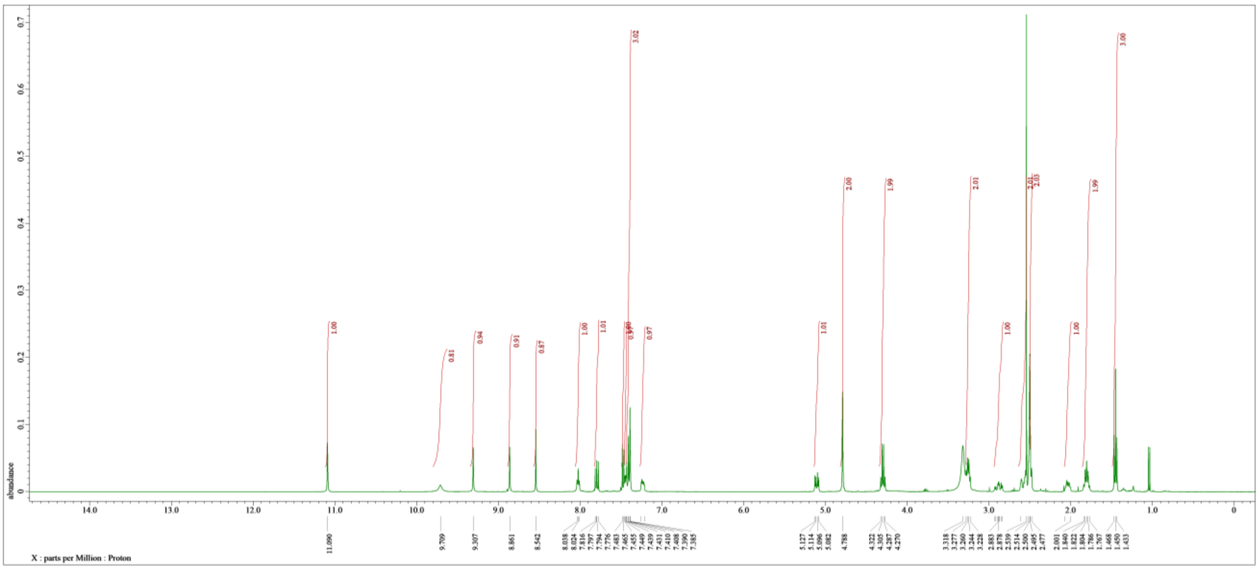

**Figure S22:** $^1$H NMR spectrum of **LP15**, 400 MHz, solvent: DMSO-$d_6$

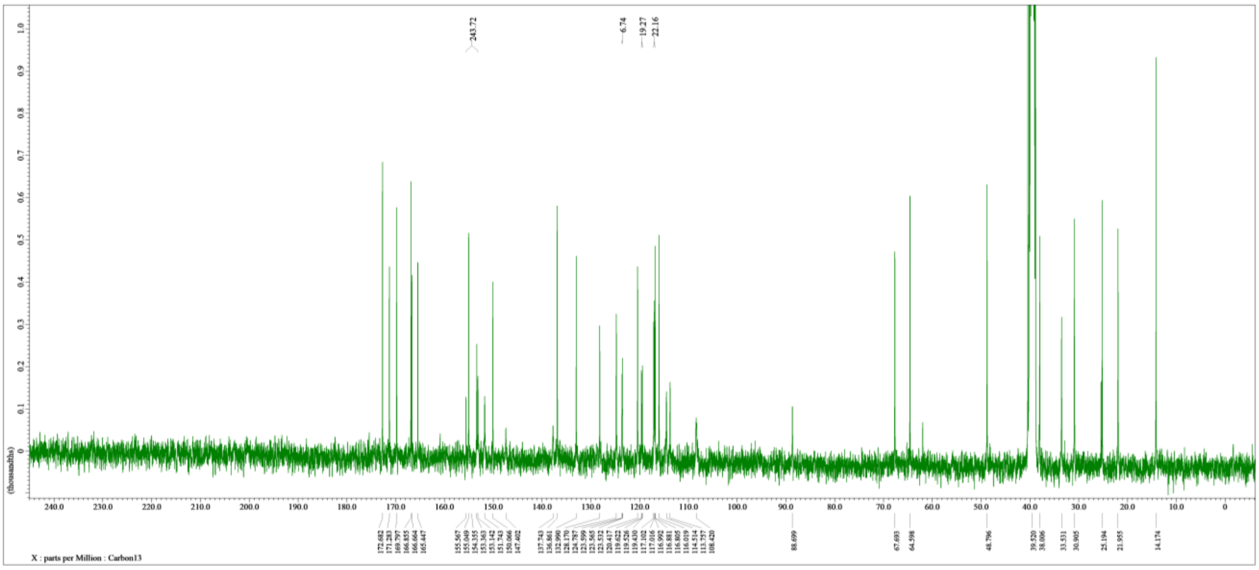

**Figure S23:** $^{13}$C NMR spectrum of **LP15**, 100 MHz, solvent: DMSO-$d_6$

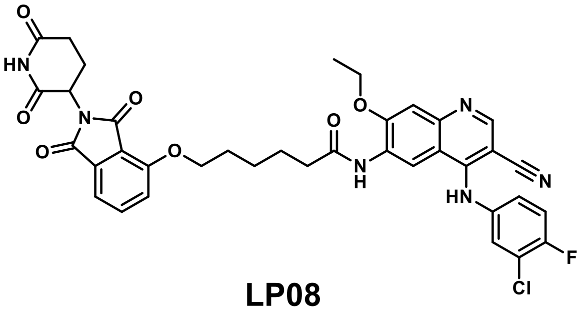

**LP08**

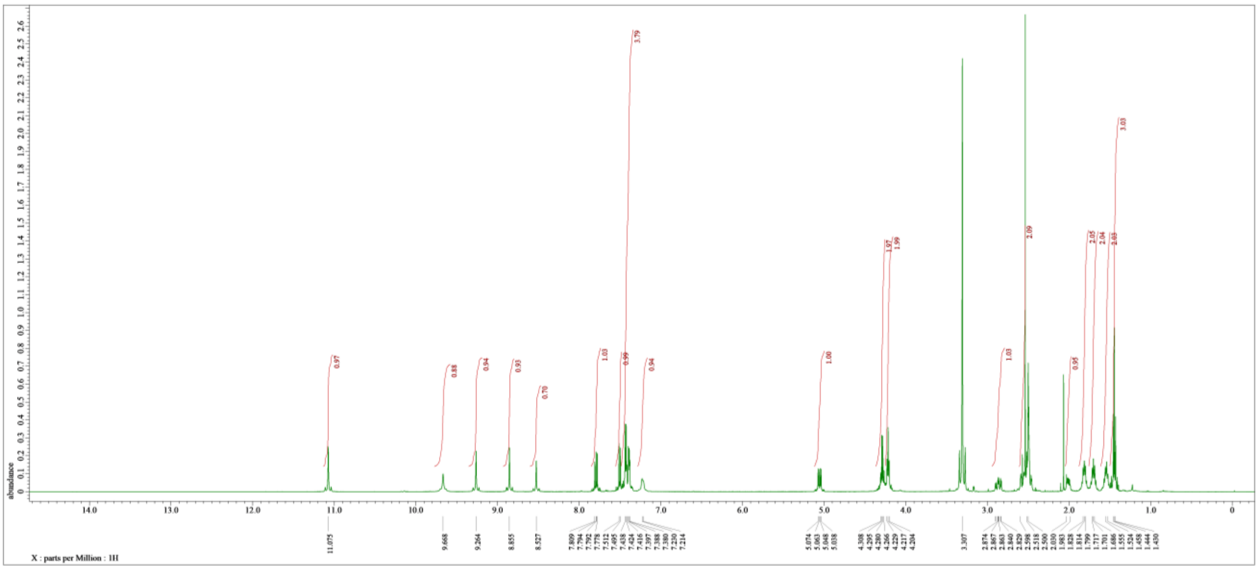

**Figure S24:** $^{1}H$ NMR spectrum of **LP08**, 500 MHz, solvent: DMSO-$d_6$

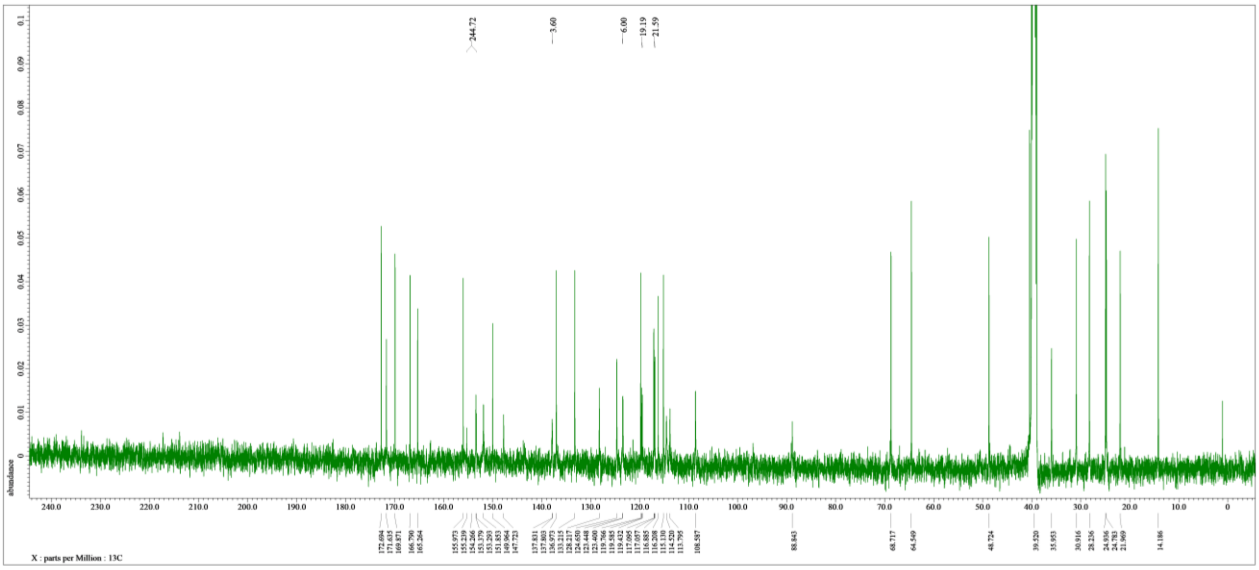

**Figure S25:** $^{13}C$ NMR spectrum of **LP08**, 125 MHz, solvent: DMSO-$d_6$

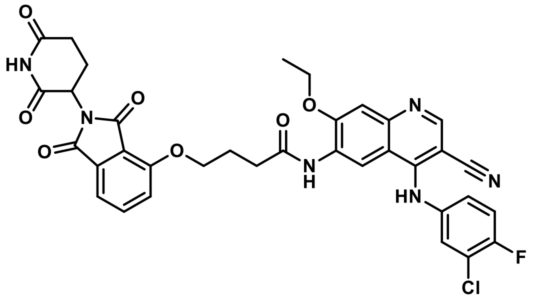

**LP04**

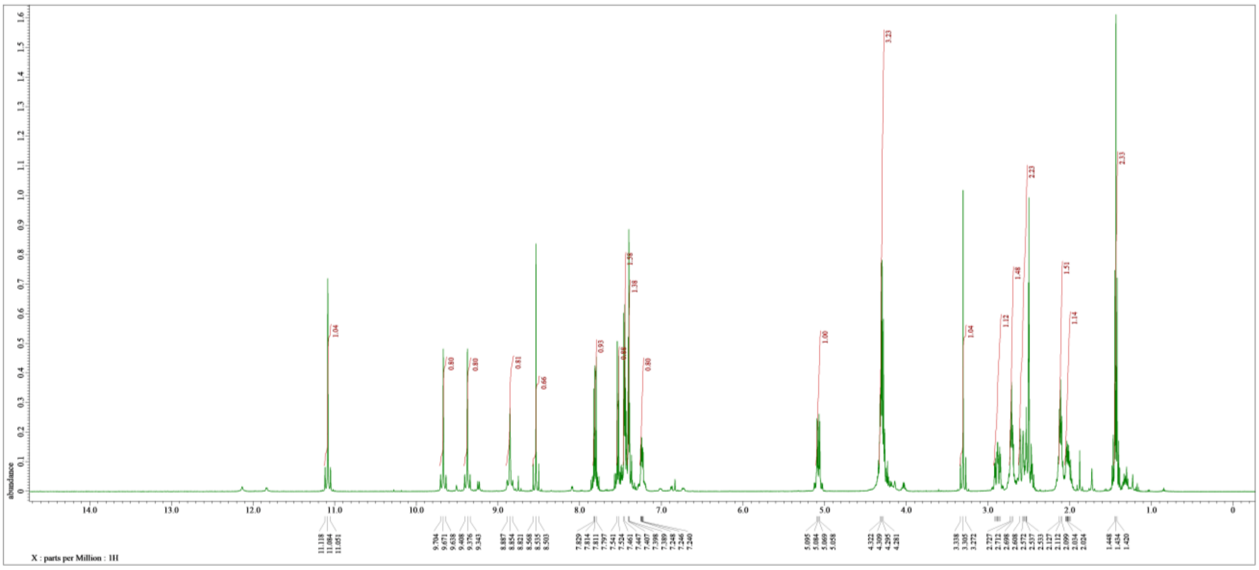

**Figure S26:** $^{1}$H NMR spectrum of **LP04**, 500 MHz, solvent: DMSO-$d_6$

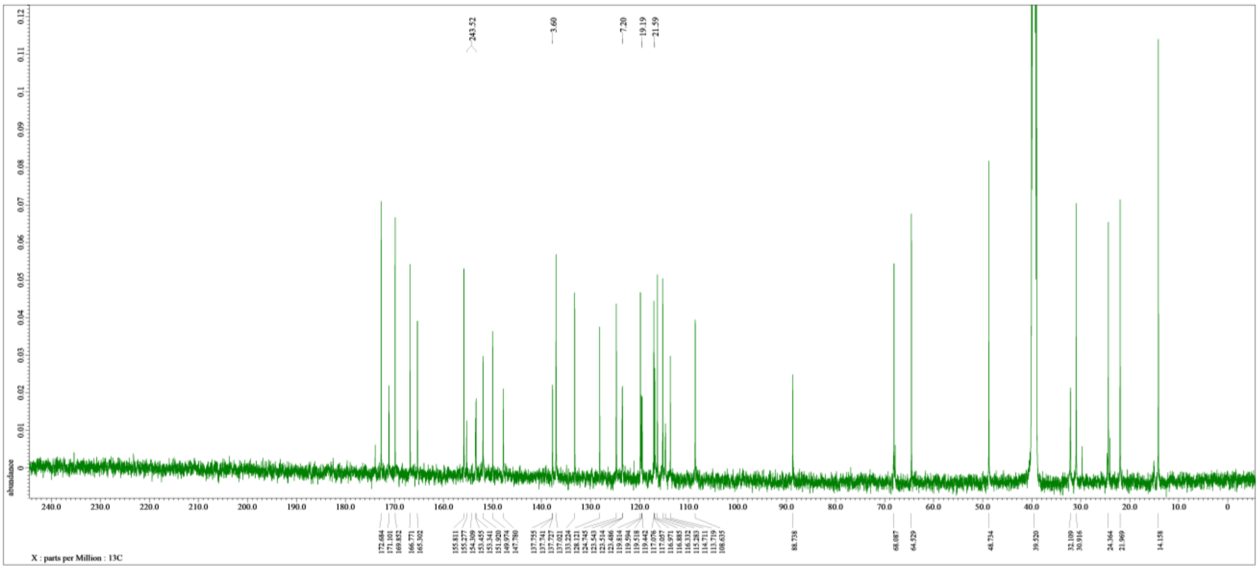

**Figure S27:** $^{13}$C NMR spectrum of **LP04**, 125 MHz, solvent: DMSO-$d_6$

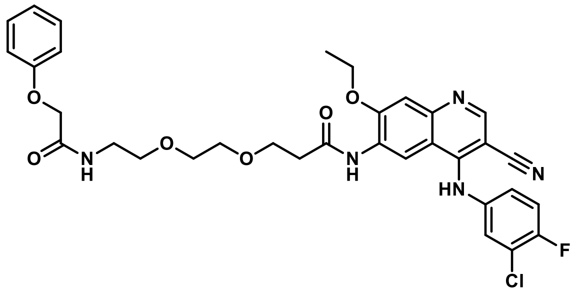

**LB06**

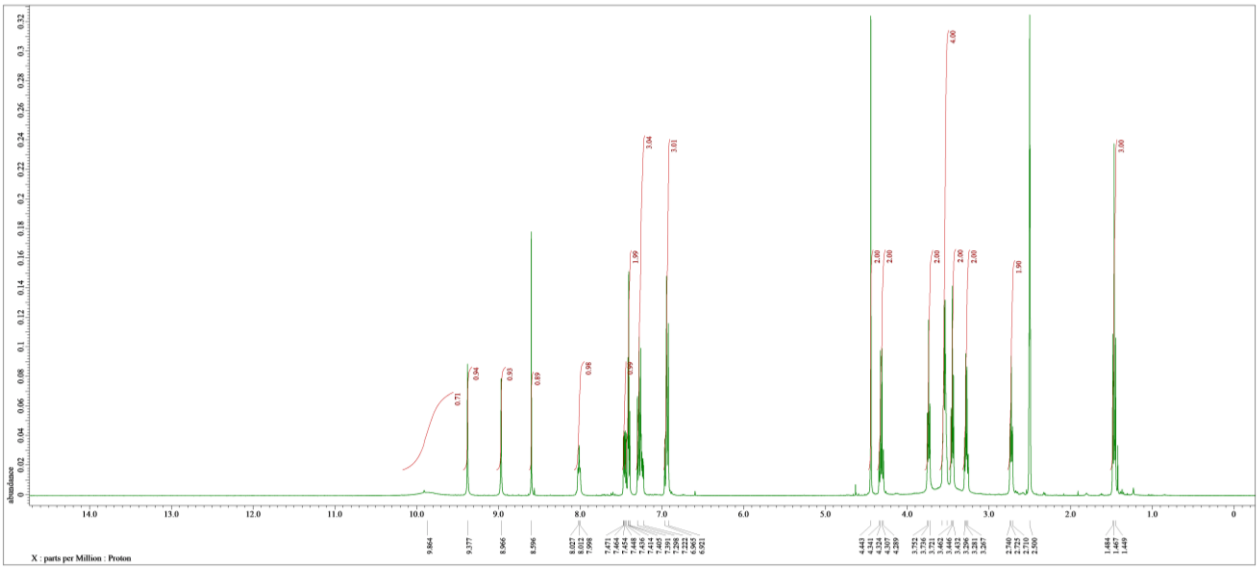

**Figure S28:** $^{1}$H NMR spectrum of **LB06**, 400 MHz, solvent: DMSO-$d_6$

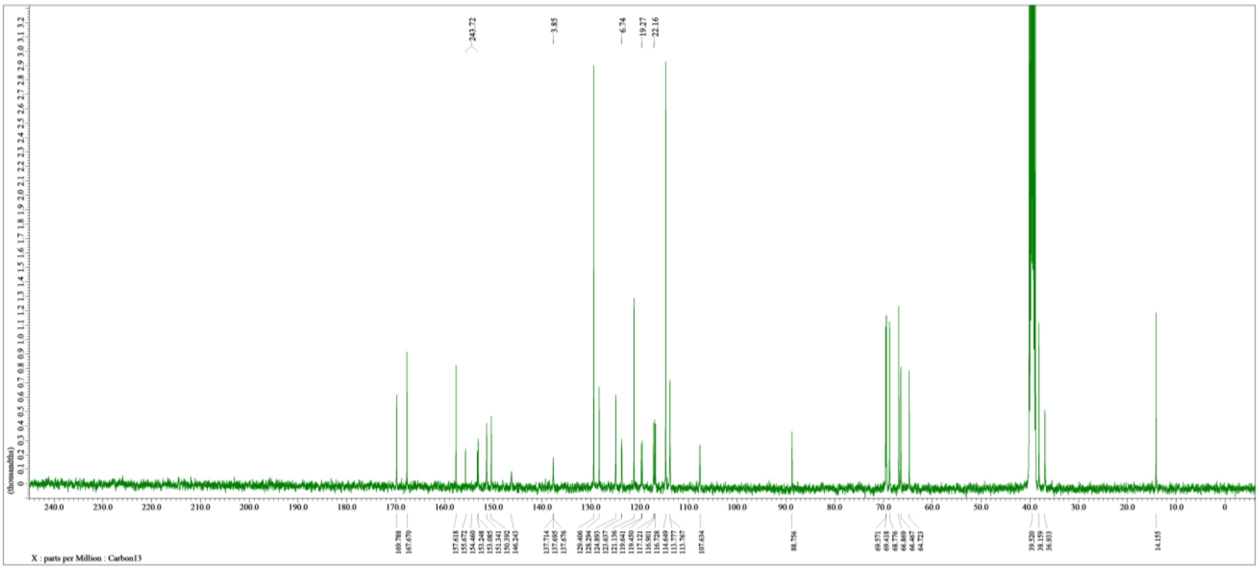

**Figure S29:** $^{13}$C NMR spectrum of **LB06**, 100 MHz, solvent: DMSO-$d_6$
